# Evolutionary branching points in multi-dimensional trait spaces

**DOI:** 10.1101/2025.10.20.683593

**Authors:** Hiroshi C. Ito, Akira Sasaki

**Affiliations:** Research Center for Integrative Evolutionary Science, The Graduate University for Advanced Studies, SOKENDAI, Hayama, Kanagawa 240-0193, Japan; Evolution and Ecology Program, International Institute for Applied Systems Analysis, A-2361 Laxenburg, Austria

**Keywords:** adaptive dynamics, convergence stability, canonical equation, Lyapunov function, multivariate trait space, adaptive evolution

## Abstract

Ecological interactions play a crucial role in driving speciation and adaptive radiation. According to the adaptive dynamics theory of evolution in one-dimensional trait spaces, such evolutionary diversifications can occur when evolutionary branching points (convergence stable and evolutionarily unstable singular points) exist in the trait spaces. However, these evolutionary diversifications may occur across multiple traits at once, as exemplified by various within-species ecotypic diversifications involving multiple traits or genes. This paper attempts to extend this branching point condition to evolution in multi-dimensional trait spaces. Two criteria, termed branching possibility and branching inevitability, are developed to evaluate the likelihood of evolutionary branching in multi-dimensional trait spaces. Branching possibility guarantees the existence of evolutionary pathways that lead to branching (where deterministic parts of these pathways are described with the canonical equation of adaptive dynamics theory). Branching inevitability guarantees a steady progression of evolutionary branching (where the progression degree is described with a time-increasing function related to the local Lyapunov function, the potential function in an evolutionary potential game, and Fisher’s fundamental theorem). Numerically simulated evolutions in multi-dimensional trait spaces show that points having branching possibility or inevitability can induce evolutionary branching not only under asexual reproduction but also under sexual reproduction.

## 1 Introduction

Ecological interactions can drive various evolutionary phenomena in biological communities such as coevolution, extinction, and diversification. Especially in sympatric and parapatric settings, disruptive and diversifying selections caused by ecological interactions are thought to have been playing a crucial role in speciation and adaptive radiation in various taxa (Schluter 2000; Dieckmann et al. 2004) including cichlids (Seehausen 2015) and anoles lizards (Muñoz et al. 2023). Because biological communities are generally thought to have been evolving in multi-dimensional trait spaces (Lande 1979; Lande and Arnold 1983; Blows 2007; Metz 2011), such a disruptive or diversifying selection can act not only on a single one-dimensional trait but also on multiple traits simultaneously, as exemplified by various within-species ecotypic diversifications involving multiple traits or genes (Johannesson et al. 2026), including switchgrass *Panicum virgatum* (Lowry et al. 2014), saltmarsh beetle *Pogonus chalceus* (Belleghem et al. 2018), three-spined stickleback *Gasterosteus aculeatus* (Dean et al. 2023), and bottlenose dolphin *Tursiops truncatus* (Louis et al. 2021). However, we have only limited theoretical understanding of the multi-dimensional evolutionary diversification through ecological interactions, due to the complexity of this phenomenon (Ito et al. 2009; Doebeli and Ispolatov 2010, 2017).

In adaptive dynamics theory, evolutionary diversification of a biological population into multiple distinct populations through ecological interactions is called evolutionary branching (Metz et al. 1996; Geritz et al. 1997). For a population evolving in a one-dimensional trait space, the likelihood of its evolutionary branching is well characterized by whether the space has an evolutionary branching point (Metz et al. 1996; Geritz et al. 1997). An evolutionary branching point is a convergence stable point (i.e., a point attractor for a monomorphic population through directional evolution (Eshel 1983)) that is not evolutionarily stable (i.e., the population can still be invaded by some slightly deviated mutants even after it has reached the point (Maynard Smith and Price 1973)). In a one-dimensional trait space, the existence of an evolutionary branching point guarantees the evolutionary branching of an asexual monomorphic population in its neighborhood, under rare and small mutations (Metz et al. 1996; Geritz et al. 1997).

The above direct connection in one-dimensional trait spaces between convergence stable non-ESSes and evolutionary branching is not readily extended for the higher-dimensional trait spaces, even under rare and small mutations (Doebeli 2011). A key complexity of the higher-dimensional evolutionary branching is that convergence stability has three classes: convergence stability (convergence in the *mean* trajectory of directional evolution under a *given* mutational covariance matrix), strong convergence stability (convergence in the *mean* trajectory of directional evolution under *any* regular mutational covariance matrix), and absolute convergence stability (convergence in *all possible* trajectories of directional evolution formed by an *arbitrary* set of sequential invasions by mutants having positive invasion fitnesses) (Leimar 2009). All absolutely convergence stable points are contained in strongly convergence stable points, and all strongly convergence stable points are contained in convergence stable points.

The complexity of convergence stability described above affects not only how a monomorphic population evolutionarily converges to a singular point but also how dimorphism may emerge and result in evolutionary divergence, and hence causing difficulty in the analysis of evolutionary branching. Although considerable progress has been made (e.g., Geritz et al. 2016), the conditions for existence of points ensuring the occurrence of evolutionary branching have not been established in multi-dimensional trait spaces. So far, strongly convergence stable non-ESSes have been treated as candidates for evolutionary branching points (Vukics et al. 2003; Ackermann and Doebeli 2004; Ito and Shimada 2007; Ito and Dieckmann 2012, 2014; Geritz et al. 2016; Ito and Sasaki 2016, 2020).

The present study introduces two analytically tractable criteria for evaluating the likelihood of evolutionary branching associated with convergence stable non-ESSes. The “branching possibility” ensures the existence of evolutionary paths that lead to branching, while the “branching inevitability” ensures that the evolutionary trajectory heads towards branching. The branching possibility is based on the canonical equation of adaptive dynamics theory (Dieckmann and Law 1996), while the branching inevitability is based on a time-increasing function related to the local Lyapunov function. For trait spaces of arbitrary dimensions, we show that the branching possibility is guaranteed not only for strongly convergence stable non-ESSes, but also for convergence stable non-ESSes that do not have strong convergence stability yet satisfy the conditions for “locally stable dimorphic divergence” (described later). We also show that all absolutely convergence stable non-ESSes have the branching inevitability; there exist time-increasing functions that guarantee the steady progression of evolutionary branching under deterministic population dynamics.

The remainder of this paper is structured as follows. Section 2 explains the basic framework of adaptive dynamics theory for multi-dimensional trait spaces. Section 3 describes the algorithms for numerical simulation. Section 4 presents our developed conditions for the branching possibility (Section 4.1) and branching inevitability (Section 4.2) and then shows whether convergence stable non-ESSes of each convergence class may have either or both of the branching possibility and inevitability (Section 4.3). Evolutionary branchings induced by these points are numerically shown for both asexual and sexual populations. Additionally, a simple application example of the developed branching point conditions is shown (Section 4.4). Section 5 discusses the present results in connection with relevant studies.

## 2 Analytical method

In this section, we explain the basic framework of adaptive dynamics theory for multi-dimensional trait spaces.

### 2.1 Basic assumptions of adaptive dynamics theory

We consider a trait space **s** = (*x*_1_, …, *x*_*L*_)^T^ of an arbitrary dimension denoted by *L* (≥ 2). We call phenotype (also called ecotype) as a distinct set of multiple traits and suppose that there are *M* coexisting phenotypes with trait vectors **s**_1_, …, **s**_*M*_. We write the dynamics of their population sizes *n*_1_, …, *n*_*M*_ as

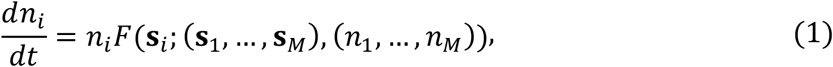

where *F*(**s**_*i*_; (**s**_1_, …, **s**_*M*_), (*n*_1_, …, *n*_*M*_)) describes the fitness of phenotype **s**_*i*_ = (*x*_*i*,1_, …, *x*_*i,L*_ )^T^. We also assume sufficiently low mutation rates and small mutational step sizes such that the population dynamics is almost at equilibrium whenever a mutant emerges. In this case, the evolutionary dynamics can be described as a sequential transitions of resident phenotypes formed by repeated mutant invasions, called a trait substitution sequence (Metz et al. 1996). When there exists only a single resident phenotype **s** = (*x*_1_, …, *x*_*L*_)^T^ with equilibrium population size 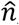, a mutant phenotype 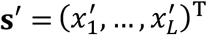 can invade if it has a positive invasion fitness, given by its initial per-capita growth rate:

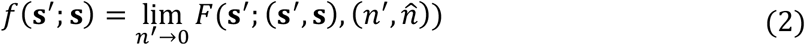

with *n*′ describing the mutant’s population size. Here, we assume that the fitness function, equation (1), satisfies the applicability conditions of adaptive dynamics theory under an arbitrary set of residents, so that the fitness landscape (e.g., fitnesses of various mutant phenotypes represented as a distribution on the trait space) changes only slightly throughout the population dynamics triggered by a mutant invasion, as long as the mutant phenotype is similar to either of the resident phenotypes (Ito et al. 2020). In this case, the invasion fitness function can be uniquely obtained from equation (1) as a smooth function of the phenotypes of a mutant and residents.

### 2.2 Directional evolution and evolutionarily singular points

The fitness gradient at resident phenotype **s** is given by

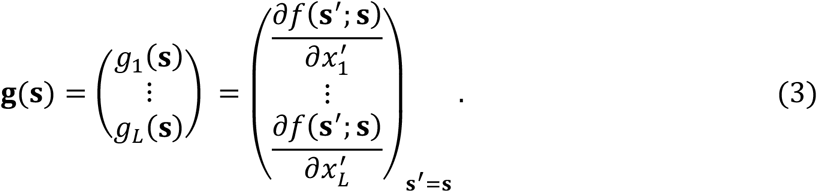

As long as this gradient is sufficiently steep so that the directional selection is dominant against the stabilizing/disruptive selection and the other selection modes (i.e., the linear term of the Taylor expansion of *f*(**s**′; **s**) is much greater than the second- and higher-order terms, allowing *f*(**s**′; **s**) ≃ **g**(**s**)^T^[**s**′ − **s**]), the invading mutant replaces the resident to become a new resident. In such a case, repeated mutant invasions induce directional evolution. The mean path among all possible directional evolutions formed by sequential mutant invasions is approximately described by an ordinary differential equation, called the canonical equation (Dieckmann and Law 1996):

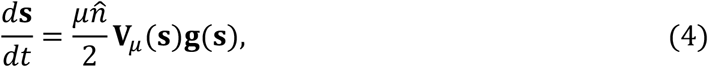

where *μ* describes the mutation rate per birth, and **V**_*μ*_(**s**) is an *L*×*L* symmetric matrix describing the mutational covariance matrix that can vary depending on **s** under the assumption of symmetric mutation with small step sizes. This directional evolution continues until the population comes close to a point **s**^∗^ at which the fitness gradient vanishes,

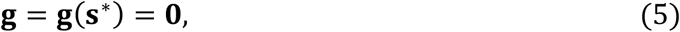

called an evolutionarily singular point.

### 2.3 Convergence stability

When the resident phenotype **s** is close to **s**^∗^ so that the fitness gradient can be linearly approximated as **g**(**s**) ≃ **C**^T^[**s** − **s**^∗^] with

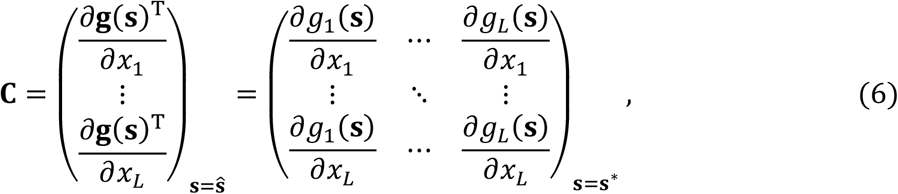

the leading order term of equation (4) is expressed as

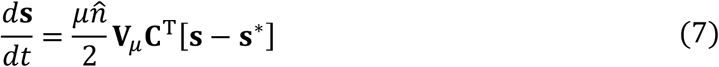

with **V**_*μ*_ = **V**_*μ*_(**s**^∗^). Based on this equation, the three classes of convergence stability are described as follows (Leimar 2009; Doebeli 2011). First, **s**^∗^ is convergence stable, when all eigenvalues of **V**_*μ*_**C**^T^ have negative real parts, expressed as

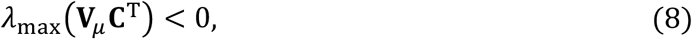

where *λ*_*max*_(**A**) gives the maximum real parts of the eigenvalues for a matrix **A**. Equation (8) guarantees the convergence of **s** toward **s**^∗^ through the mean path of directional evolution described by equation (7), under a given mutational covariance matrix, **V**_*μ*_. Hence, it is possible that equation (8) holds under some **V**_*μ*_ but does not hold under other **V**_*μ*_. Second, **s**^∗^ is *strongly* convergence stable when all eigenvalues of the symmetric part of **C** are negative (i.e., [**C** + **C**^T^]⁄2 is negative definite), expressed as

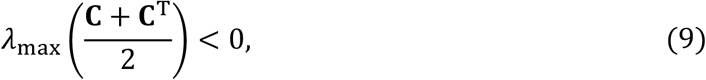

which guarantees the convergence of **s** toward **s**^∗^ under any positive definite symmetric matrix **V**_*μ*_, through the mean path of directional evolution described with equation (7). However, because equation (9) guarantees the convergence of only the mean evolutionary path among all possible evolutionary trajectories, some rare paths might not show the convergence even when equation (9) holds. Third, **s**^∗^ is *absolutely* convergence stable, when **C** is symmetric and its eigenvalues are all negative, expressed as

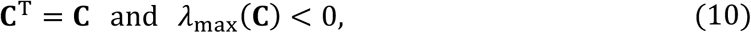

guaranteeing the convergence of **s** toward **s**^∗^ under all possible evolutionary trajectories formed by sequentially invading mutants with positive invasion fitnesses under small mutations in arbitrary directions. Note that all absolutely convergence stable points are included in the set of strongly convergence stable points, i.e., equation (10) implies equation (9), and that all strongly convergence stable points are included in convergence stable points, i.e., equation (9) implies equation (8).

### 2.2 Evolutionary stability

A singular point **s**^∗^ is (locally) evolutionarily stable, i.e., any mutant **s**′ slightly deviating from **s**^∗^ cannot invade the resident at **s**^∗^ (i.e., *f*(**s**′; **s**^∗^) < 0), if the symmetric matrix

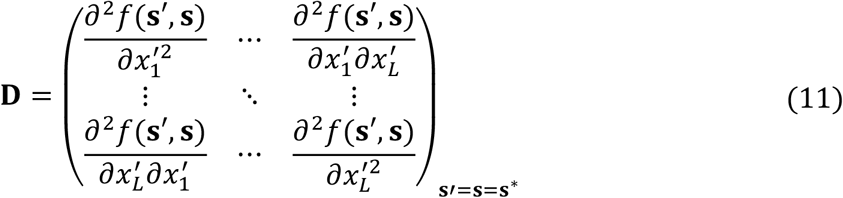

has only negative eigenvalues. Conversely, **s**^∗^ is evolutionarily unstable if the maximum eigenvalue of **D** is positive, expressed as

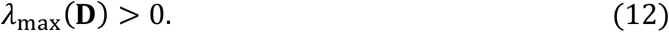

In this case, the resident placed at **s**^∗^ can be invaded by mutants deviating along the eigenvectors for the positive eigenvalues of **D**.

### 2.4 Requirements for evolutionary branching points

According to Doebeli (2011), the requirements for a point **s**^∗^ to be an evolutionary branching point are described as follows:

i. **s**^∗^ is convergence stable.
ii. **s**^∗^ is evolutionarily unstable in at least one direction of trait space.
iii. Mutual invasibility (i.e., *f*(**s**′; **s**) > 0 and *f*(**s**; **s**′) > 0) holds around **s**^∗^ in the directions of evolutionary instability given by (ii) (so that polymorphism can emerge).
iv. The different phenotypic branches constituting the protected polymorphism given by (iii) diverge evolutionarily from **s**^∗^.

In one-dimensional trait spaces, (i) and (ii) are sufficient for (iii) and (iv) (Metz et al. 1996; Geritz et al. 1997). In higher-dimensional trait spaces, however, (i) and (ii) are not sufficient for (iii) and (iv) (Doebeli 2011). Moreover, (i-iii) may not be sufficient for (iv) (Doebeli 2011; Geritz et al. 2016). So far, condition (iv) has been mathematically obtained in the framework of quantitative genetics under the assumption of weak selection such that the timescales of population dynamics and evolutionary dynamics are not separated (Geritz et al. 2016), which may correspond to the initial stage of evolutionary divergence. In the present paper, we obtain condition (iv) on a scale comparable to the process of monomorphic convergence corresponding to condition (i), by describing the evolutionary divergence in the framework of the canonical equation.

## 3 Numerical method

To examine our analytically developed conditions for evolutionary branching described in the next section, we numerically simulated evolutionary dynamics as trait substitution sequences under the assumptions of asexual reproduction and rare mutation, by using a phenotype-based algorithm called the oligomorphic stochastic model (Ito and Dieckmann 2007; Ito and Sasaki 2023) (see Appendix S1.1 for details). Additionally, we simulated evolutionary dynamics under the assumptions of sexual reproduction and non-rare mutation, by using an individual-based algorithm called the polymorphic stochastic model (Ito and Dieckmann 2007; Ito and Sasaki 2023) (see Appendix S1.2 for details). In the latter algorithm, individuals are diploid and have many additive loci that determine their trait values. To make prezygotic isolation achievable, we introduced a mating system that allows the evolution of assortative mating through “sensory drive”, which is thought to have facilitated sympatric and parapatric speciation in various animals (Dell’Aglio et al. 2024). Specifically, we introduced a scalar “mating trait” denoted by *m* affecting body colors and patterns, so that the visibility of the *k*th male individual (as a mating partner) for the *k*′ th female individual is given by

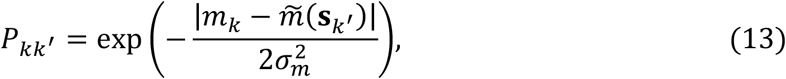

where *m*_*k*_ denotes the mating trait of the *k*th male, and 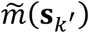 describes the potentially most outstanding or encounterable male for the *k*′th female, which is a function of the female’s ecological phenotype, **s**_*k*′_. Although trait *m* can be included in **s**, we treat *m* separately from **s** so that we can easily compare sexual and asexual populations in their simulated evolutionary branching. For the same purpose, we assume that the fitness function in equation (1) does not depend on trait *m*.

## 4 Results

This section addresses the conditions for the branching possibility and branching inevitability in relation to the requirements (i-iv) for evolutionary branching points outlined in Section 2.4. Then, we examine whether each of the three classes of convergence stable points exhibits branching possibility, inevitability or both.

### 4.1 Branching possibility

Under coexisting phenotypes **s**_1_, …, **s**_*M*_ in the neighborhood of an evolutionarily singular point **s**^∗^, we can approximate the fitness for phenotype **s**′ as a quadratic function of **s**′ (following Meszéna et al. 2005):

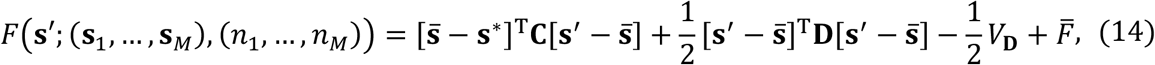

where 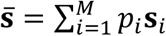 with 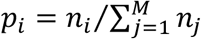 is the average phenotype among the coexisting phenotypes, 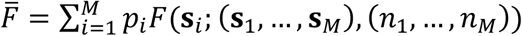 is their average fitness, and 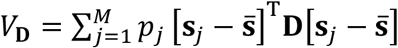 is their phenotypic variance weighted with **D** (see Appendix S2 for the derivation). Also, without loss of generality, we can approximate the mutational covariance matrix as a constant, **V**_*μ*_(**s**^∗^) = **V**_*μ*_, because equation (14) is kept identical (up to the second order terms of **s**′) even after the coordinate normalization for making **V**_*μ*_(**s**) locally constant around **s**^∗^ (Ito and Sasaki 2020). Then, the following proposition holds, as shown in Appendix S2:

If an evolutionarily singular point **s**^∗^ satisfies conditions (15) defined below, then **s**^∗^ has the *branching possibility* in the sense that there is a non-zero probability of eventual evolutionary branching for an arbitrary monomorphic resident **s** in the neighborhood of **s**^∗^, through trait substitution sequences consisting of the deterministically describable convergence of a monomorphic resident toward **s**^∗^ (monomorphic convergence), followed by the stochastic emergence of dimorphism and the initial divergence of the dimorphic phenotypes, and then by their deterministically describable divergence through directional evolution (dimorphic divergence).

Branching possibility conditions:

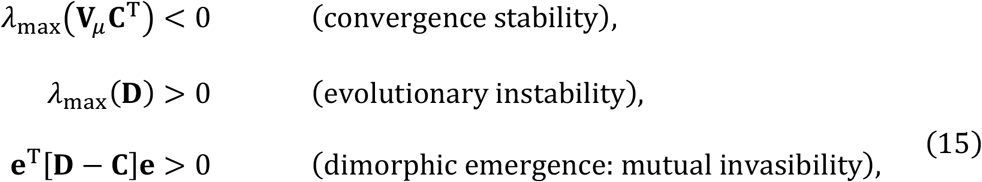

with **C**_d_ = **C** + **Cee**^T^**C**⁄{**e**^T^[**D** − **C**]**e**}, where 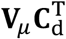 determines the convergence stability of the intermediate phenotype between the dimorphic residents (i.e., 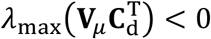 guarantees the convergence toward **s**^∗^ of the intermediate phenotype given by [**s**_1_ + **s**_2_] / 2 with **s**_1_ and **s**_2_ denoting the two resident phenotypes, as shown in equation (S2.102) in Appendix S2.7), and where **e** = **v**_*λ*max_(**V**_*μ*_**D**) is a unit vector describing the direction of dimorphic divergence, and **v**_*λ*max_(**A**) gives the eigenvector corresponding to *λ*_max_(**A**) for a matrix **A**. Note that the four inequalities in equation (15) respectively correspond to the requirements (i-iv) in Section 2.4.

The fourth inequality in equation (15), the condition for “locally stable dimorphic divergence,” guarantees the steady progression of diversifying directional evolution of two coexisting monomorphic populations described by the canonical equation, once the two populations have sufficiently diverged from each other along the direction of **e**. That is, for the directionally evolving two populations with their resident phenotypes denoted by **s**_1_ and **s**_2_, the dynamics of **u** = [**s**_1_ − **s**_2_]/|**e**^T^[**s**_1_ − **s**_2_]| − **e** and 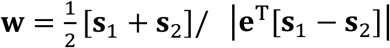 has a locally stable fixed point (**u, w**) = (**0, 0**) under equation (15), as derived in Appendix S3.3. Notably, the condition for locally stable dimorphic divergence, 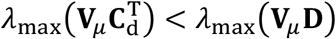, implies that the steady progression of dimorphic divergence is realized when the growth rate *λ*_max_(**V**_*μ*_**D**) of the phenotypic difference |**s**_1_ − **s**_2_| is greater than the deviation rate 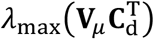 of the intermediate phenotype 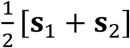 from the singular point, as shown in equation (S2.102) in Appendix S2.7.

Figure 1 shows numerically simulated evolutionary branchings for an initially monomorphic population around **s**^∗^ satisfying the branching possibility conditions. This simulation calculates trait substitution sequences (i.e., asexual reproduction with rare mutation) under the locally approximated fitness function, equation (14). Even under the assumption of sexual reproduction with non-rare mutation, as shown in figure 2, points satisfying the branching possibility conditions can induce evolutionary branching, when assortative mating can easily evolve through evolutionary emergence of strong correlation between the ecological phenotype (**s**) and the mating trait (*m*), as shown in the small panels in figure 2.

**Figure 1.**
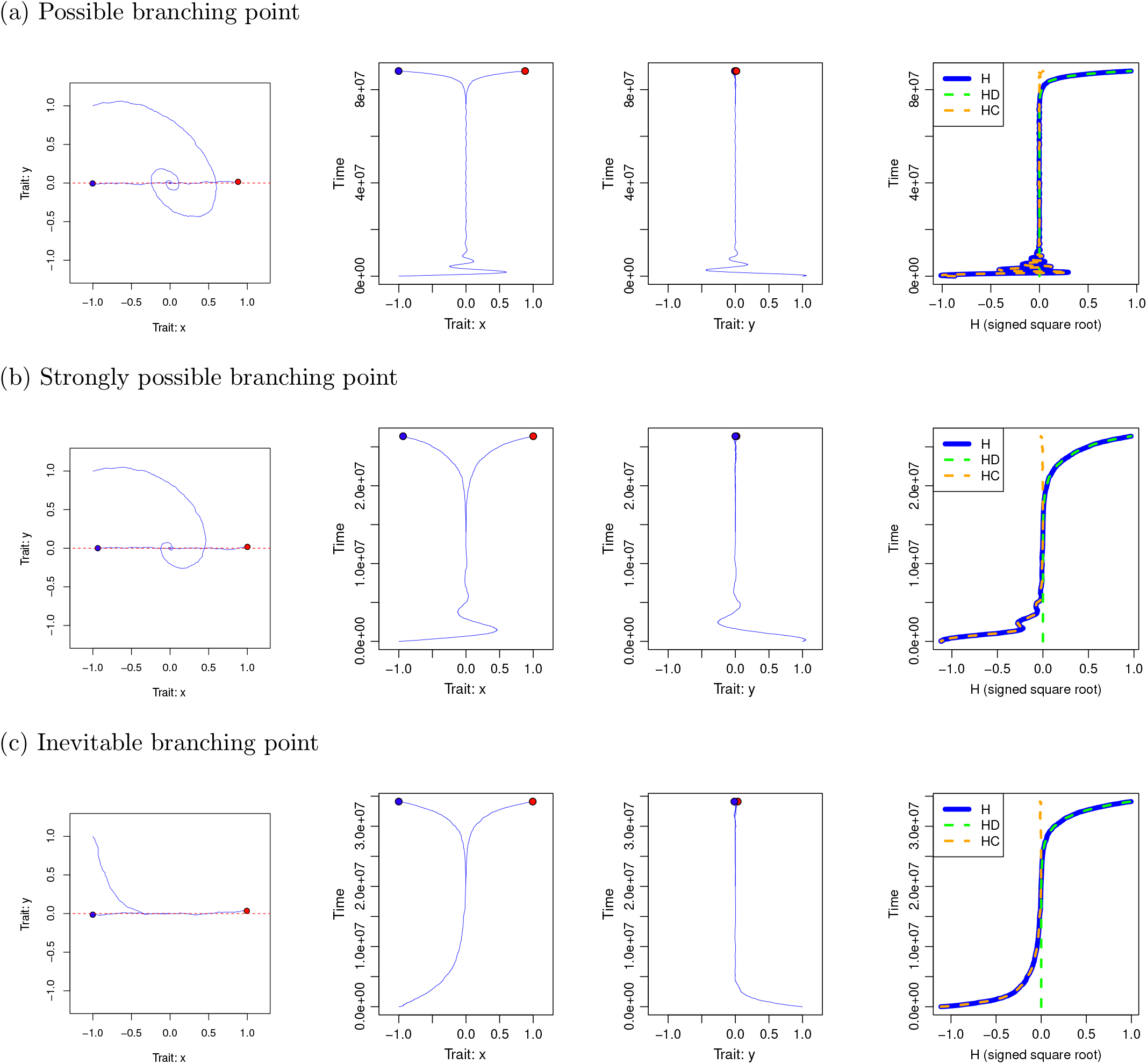
Evolutionary branching in simulated evolution under the assumption of asexual reproduction and rare mutation, in a two-dimensional trait space, **s** = (*x, y*)^T^, induced by (a) a possible branching point, (b) a strongly possible branching point, or (c) an inevitable branching point, which respectively correspond to a convergence stable, strongly convergence stable, and absolutely convergence stable non-ESS (summarized in Table 1). See Section 3 for the simulation algorithm. The initial phenotype is **s**_0_ = (™1,1)^T^. Thin blue curves indicate evolutionary trajectories. Red and blue dots indicate the dimorphic residents at the end of simulation. Red dashed lines in the left-most panels indicate the predicted direction **e** = (1,0)^T^ of dimorphic divergence, given by equation (15). The right-most panel shows the time changes of the value for the function *H* = *H*_*C*_ + *H*_*D*_ defined in equation (17). Under the existence of an inevitable branching point, as shown in panel (c), *H* increases monotonically, guaranteeing the steady progression of evolutionary branching. Parameters: 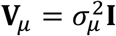 with *σ*_*μ*_ = 0.005 and 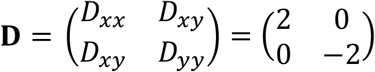 for all (a-c); 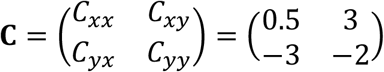 for (a), 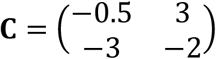 for (b), and 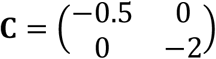 for (c).

**Figure 2.**
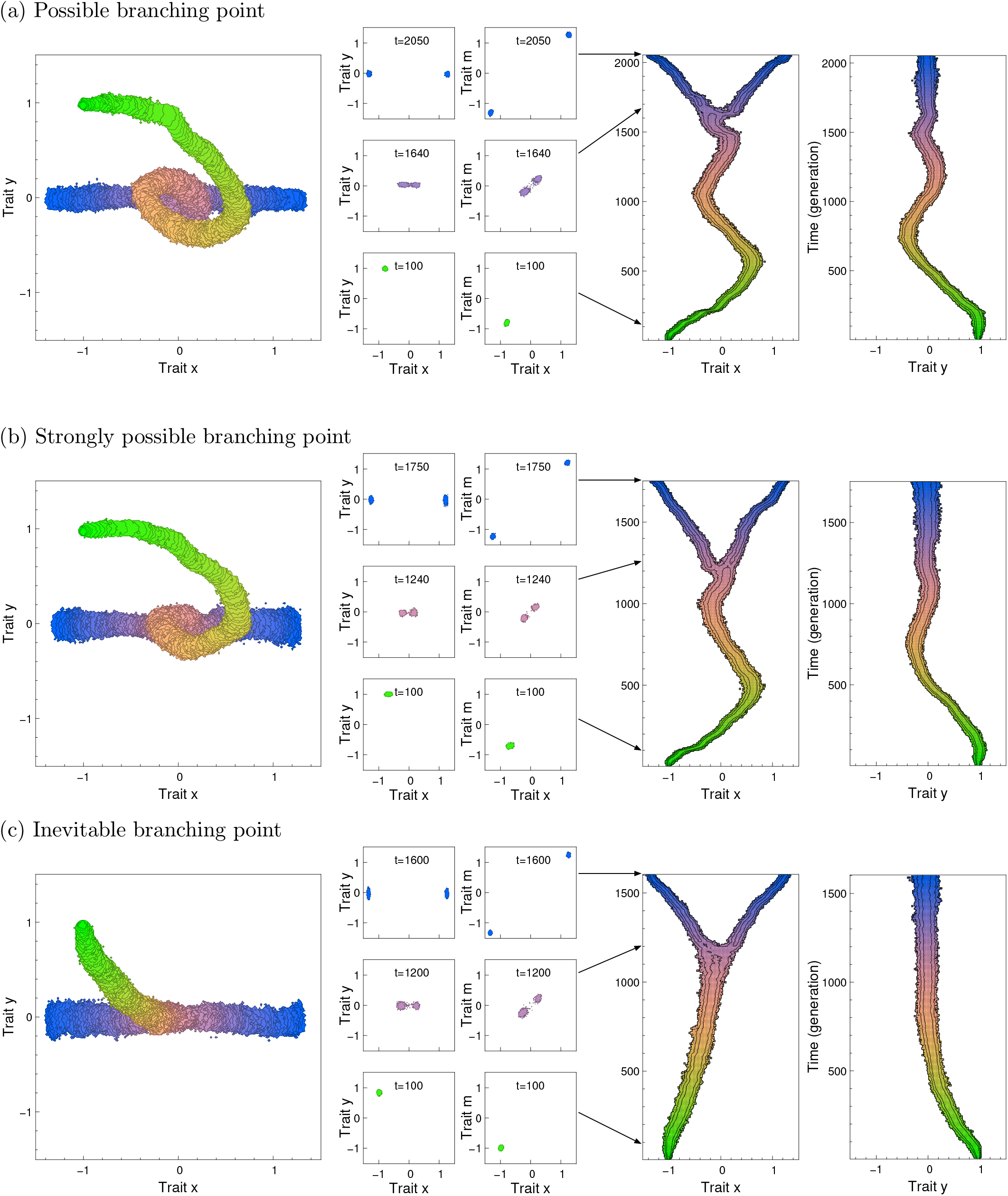
Evolutionary branching in simulated evolution under the assumption of sexual reproduction and non-rare mutation, for the same ecological settings with panels (a-c) in figure 1 (see Section 3 for the simulation algorithm). In each panel, colored areas indicate observation of at least one individual, and the color gradation indicates time (i.e., green and blue colors respectively indicate the initial and last states of the simulation). The black contours in the left two panels indicate 20 and 200 individuals. Parameters for ecological settings are the same with those used in figure1. Parameters for reproduction and mutation are as follows. Birth rate for phenotype **s**: *b*(**s**) = 1. Death rate: *d*(**s**) = 1 ™ *F*(**s**; (**s**_1_, …, **s**_*M*_), (*n*_1_, …, *n*_*M*_)/*K*_0_) with *K*_0_ = 5000 describing the expected population size sustained in the system. Number of loci for each trait (diploid): 30. Mutation rate per locus (stepwise mutation): 5 × 10^-4^. The mating trait value that is most outstanding (or encounterable) for a female with phenotype 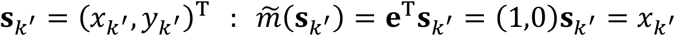, meaning that the *k*th male is the most outstanding when its mating trait *m*_*k*_ is equal to *x*_*k&*_′.

### 4.2 Branching inevitability

As shown in Appendix S4, the following proposition holds under equation (14):

**Table 1.** Three classes of convergence stable non-ESSes in an arbitrary *L*-dimensional trait space (*L* ≥ 2) and their branching possibility (equation (15)) and branching inevitability (equation (16)). The expression “Yes, provided” (followed by inequalities) indicates that the corresponding property holds provided the inequalities are satisfied, whereas “Yes” indicates that the corresponding property holds with no conditions required. *λ*_max_(**A**) denotes the largest real part of the eigenvalues of a matrix **A**, and **e** = **v**_*λ*max_(**V**_*μ*_**D**) gives the expected direction of the dimorphic divergence (equation (15)).

| Class of convergence stable non-ESS | Convergence stable non-ESS |  | Strongly convergence stable non-ESS | Absolutely convergence stable non-ESS |
| --- | --- | --- | --- | --- |
| | Evolutionary singularity | $\mathbf{g} = \mathbf{0}$ | $\mathbf{g} = \mathbf{0}$ | $\mathbf{g} = \mathbf{0}$ |
| | Convergence stability | $\lambda_{\max}(\mathbf{V}_\mu \mathbf{C}^T) < 0$ | $\lambda_{\max}\left(\frac{\mathbf{C} + \mathbf{C}^T}{2}\right) < 0$ | $\mathbf{C} = \mathbf{C}^T$,<br>$\lambda_{\max}(\mathbf{C}) < 0$ |
| | Evolutionary instability | $\lambda_{\max}(\mathbf{D}) > 0$ | $\lambda_{\max}(\mathbf{D}) > 0$ | $\lambda_{\max}(\mathbf{D}) > 0$ |
| Branching possibility | **YES, provided**<br>$\mathbf{e}^T[\mathbf{D} - \mathbf{C}]\mathbf{e} > 0$,<br>$\lambda_{\max}(\mathbf{V}_\mu \mathbf{C}_d^T) < \lambda_{\max}(\mathbf{V}_\mu \mathbf{D})$<br>(required only for $L \geq 3$) | | **YES** | **YES** |
| Branching inevitability | **Unknown** |  | **Unknown** | **YES** |
| Class of evolutionary branching point | Possible branching point |  | Strongly possible branching point | Inevitable branching point |

If an evolutionarily singular point **s**^∗^ satisfies the branching inevitability conditions defined below, then **s**^∗^ has *branching inevitability* in the sense that **s**^∗^ has a time-increasing function guaranteeing the inevitable and deterministic progression of evolutionary branching for an arbitrary monomorphic resident **s** in the neighborhood of **s**^∗^.

Branching inevitability conditions:

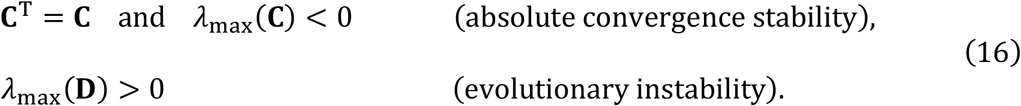

Note that all absolutely convergence stable non-ESSes have the branching inevitability. The equality **C**^T^ = **C** need not hold strictly but approximately in the sense that the effect of **C**^T^ − **C** is negligible in equation (14) (ie., **C**^T^ − **C** is of order of magnitude no greater than the typical mutational step size given by *λ*_max_ (**V**_*μ*_)^1/2^), and the same relaxation applies to the conditions for absolute convergence stability, equation (10). The time-increasing function guaranteeing evolutionary branching is defined as

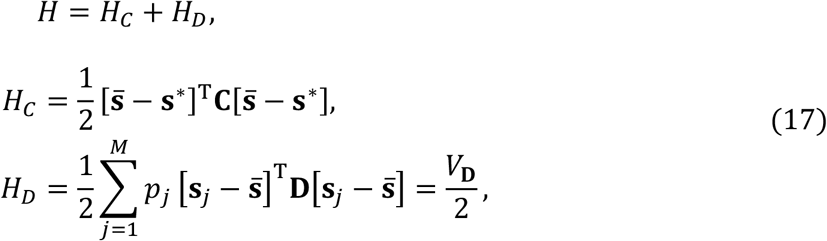

where *H*_*C*_ measures the distance from **s**^∗^ to the mean phenotype 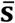 of the coexisting phenotypes (weighted by **C**), and *H*_*D*_ measures their phenotypic variance (weighted by **D**). Note that *H*_*C*_ ≤ 0 always holds under equation (16). As derived in Appendix S4.1, the time derivative of *H* satisfies

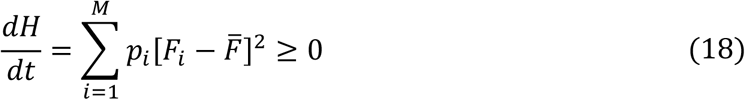

under equations (14) and (16), where *F*_*i*_ = *F*(**s**_*i*_; (**s**_1_, …, **s**_*M*_), (*n*_1_, …, *n*_*M*_)) is the fitness for the *i*th phenotype under coexisting phenotypes **s**_1_, …, **s**_*M*_, and 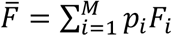 is their average fitness. Generally, *dH*⁄*dt* = 0 holds only when the population dynamics completely stops. Equation (18) guarantees monotonic increase of *H* during population dynamics triggered by each mutant invasion, until the system reaches the next equilibrium. Furthermore, under inequalities (16), the invasion of mutants is always possible for any set of resident phenotypes coexisting at their population-dynamical equilibrium (see Appendix S4.2 for the derivation). Hence, provided that changes of *H* caused directly by the emergence of mutants are negligibly small, repeated mutant invasions induce monotonic and unlimited increase of *H*, corresponding to the unlimited phenotypic divergence in the direction of positive eigenvalues of **D** (see Appendix S4.3 for the derivation). Under rare mutation, such an evolutionary diversification proceeds not as expansion of a phenotypic cloud but as evolutionary branching to new distinct phenotypes, because at most *L* + 1 phenotypes can coexist in the vicinity of an evolutionary singular point at each population-dynamical equilibrium in an *L*-dimensional trait space (Vukics et al. 2003). Equation (18) remains valid even when timescale separation between population dynamics and evolutionary dynamics is incomplete (due to non-rare mutation or weak selection), provided population dynamics of existing phenotypes are deterministically described by equations (1) and (14).

When a monomorphic population converges through directional evolution toward a point **s**^∗^ that satisfies the branching inevitability conditions, |*H*_*D*_| is kept very small, and hence the increase of *H* is realized by the monotonic increase of *H*_*C*_ from a negative value toward zero (orange dashed curve in the right panel of figure 1c). In contrast, after sufficient convergence of the population toward **s**^∗^, the increase of *H* is realized by the increase of *H*_*D*_ corresponding to the emergence of polymorphism and their divergent evolution (green dashed curve in the right panel of figure 1c).

Note also that equation (18) still applies even when **s**^∗^ is both absolutely convergence stable (**C**^T^ = **C**, *λ*_max_(**C**) < 0) and evolutionarily stable (*λ*_max_(**D**) < 0), rather than evolutionarily unstable. In this case, *H* serves as a local Lyapunov function that ensures any set of initial resident phenotypes converges to a monomorphic resident of phenotype **s**^∗^ through repeated mutant invasions (see Appendix S4.4 for details).

### 4.3 Branching possibility and inevitability for three classes of convergence stable non-ESSes

Table 1 shows whether convergence stable non-ESSes of each convergence class can have either or both of the branching possibility and inevitability (see Appendix S5 for the derivation).

Table 1 shows firstly that convergence stable non-ESSes without strong convergence can have the branching possibility, secondly that all strongly convergence stable non-ESSes have the branching possibility, and thirdly that all absolutely convergence stable non-ESSes have both the branching possibility and inevitability, as specifically described below.

First, convergence stable non-ESSes that meet two additional requirements— dimorphic emergence and locally stable dimorphic divergence—have the branching possibility, and hence they are referred to as “possible branching points.” Note that there can exist possible branching points that are not strongly convergence stable. Such possible branching points are likely to induce evolutionary branching composed of spiral monomorphic convergence followed by linear dimorphic divergence (figures 1a and 2a). This type of possible branching points can induce evolutionary branching in trait spaces of at least two to six dimensions, in simulated trait substitution sequences (figures S4-S8 in Appendix S6). Note also that in two-dimensional trait spaces the first additional condition (i.e., for dimorphic emergence) is sufficient to satisfy the second additional condition (for locally stable dimorphic divergence). Contrastingly, in higher-dimensional trait spaces, this relationship does not hold (figure 2). Hence, it is possible that the first additional condition is satisfied but not the second, in which case dimorphic divergence can start but may collapse eventually (figure 2c-d).

Second, regarding strongly convergence stable non-ESSes, they have the branching possibility under any regular positive definite **V**_*μ*_ without the additional conditions, and hence they are referred to as “strongly possible branching points” (figures1b and 2b). Note that the branching possibility conditions guarantee not a high probability but a non-zero probability of evolutionary branching for sufficiently long waiting times. Therefore, the probability of evolutionary branching under not only possible but also strongly possible branching points could be very low. Even for a strongly possible branching point, if the real parts of the eigenvalues of its **C** have much smaller absolute values than the imaginary parts, then numerically simulated trait substitution sequences can show trajectories that orbit around the point for a large number of invasions without the occurrence of evolutionary branching (see Appendix S6 for details). In simulated evolution under sexual reproduction or non-rare mutation, the likelihood of such an orbiting without evolutionary branching might be less frequent but has not been investigated extensively in the present study.

Third, regarding absolutely convergence stable non-ESSes, they have both the branching possibility and inevitability under any regular **V**_*μ*_ without the additional conditions. Hence, they are referred to as “inevitable branching points”, showing steady progression of evolutionary branching in figures 1c and 2c.

### 4.4 Application example

We show a simple application example for the conditions for possible, strongly possible, and inevitable branching points, developed in the previous section (see Appendix S7 for details of the analysis below). We consider a two-dimensional trait space **s** = (*x, y*)^T^ for describing the phenotypes of a cannibalistic population, where individuals not only consume their main resource but also prey on conspecifics. Trait *x*, describing the trait as a predator or consumer, and trait *y*, describing the trait as a prey or resource, could correspond to the foraging preference and the hiding habitat, respectively. For simplicity, we assume isotropic mutation such that 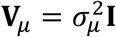 holds. For arbitrary *M* coexisting phenotypes **s**_1_, …, **s**_*M*_ with their population sizes *n*_1_, …, *n*_*M*_, we define their population dynamics as

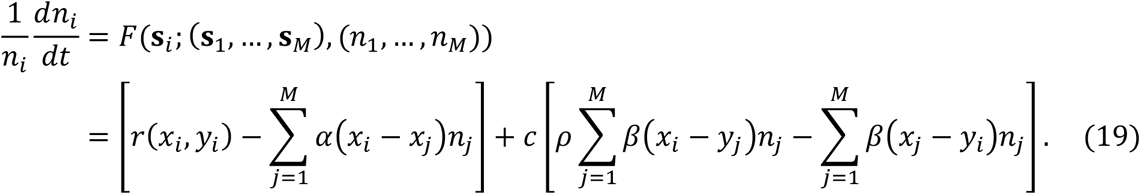

Here, *r*(*x*_*i*_, *y*_*i*_) describes the intrinsic growth rate for the *i*th phenotype sustained by the main resource in the absence of competitors. *α*(*x*_*i*_ − *x*_*j*_) describes the negative interaction between the *i*th and *j*th phenotypes through competition for the main resource. *β*(*x*_*i*_ − *y*_*j*_) describes the predation rate through cannibalistic interaction by the *i*th phenotype on the *j*th phenotype per unit population sizes for them (corresponding to the type-I functional response). A constant *c* describes the strength of cannibalistic interaction relative to resource competition for the main resource, and a constant *ρ* describes the assimilation rate for resource intake from the cannibalistic interaction, satisfying 0 ≤ *ρ* ≤ 1. We define *α*(*x*) and *β*(*x*) as

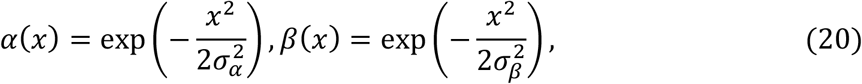

with *σ*_*α*_ and *σ*_*β*_ describing the ranges of resource competition and cannibalistic interaction, respectively. In numerically simulated evolution under equation (19) with 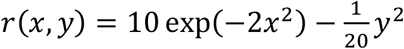, an initially monomorphic resident can repeat evolutionary branching accompanied with trophic diversification, resulting in the development of food-web structures, as shown in figure 4.

**Figure 3.**
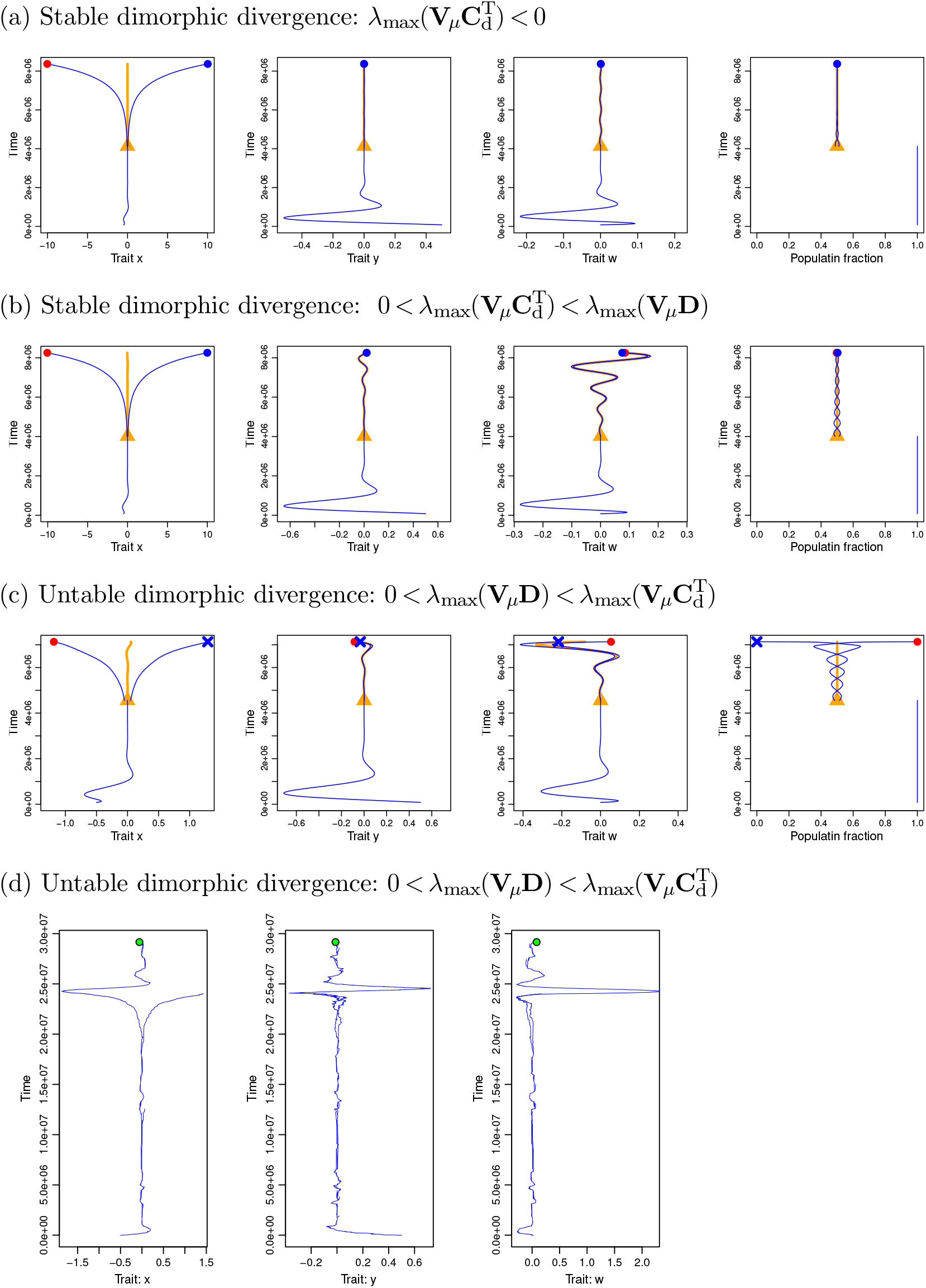
Stable and unstable dimorphic divergence in evolutionary trajectories described with the canonical equation in three-dimensional trait space **s** = (*x, y, w*)^T^. Panels (a-c) show evolutionary trajectories of monomorphic convergence and dimorphic divergence described with the canonical equation under (a) *λ*_max_(**C**_d_) < 0 (i.e., the intermediate phenotype of dimorphism converges to **s**^∗^ = (0,0,0)^T^), (b) 0 < *λ*_max_(**C**_d_) < *λ*_max_(**D**) (i.e., the intermediate phenotype deviates from **s**^∗^ but the deviation is slower than the divergence), (c) 0 < *λ*_max_(**D**) < *λ*_max_(**C**_d_) (i.e., the intermediate phenotype deviates from **s**^∗^ and the deviation is faster than the divergence). For (a) and (b) all conditions for the branching possibility (equation (15)) hold, whereas for (c) the last condition (for locally stable dimorphic divergence) does not hold although the other conditions all hold (see Appendix S5.1). The initial phenotype is **s**_0_ = (™0.5,0.5,0)^T^. In each of panels (a-c), blue curves indicate trajectories of monomorphic convergence and dimorphic divergence. The orange curve indicates the trajectory of the intermediate phenotype [**s**_1_ + **s**_2_]/2 of dimorphic residents denoted by **s**_1_ and **s**_2_ (starting from **s**_1_ = **s**^∗^ + ***δ*s** and **s**_2_ = **s**^∗^ ™ ***δ*s** with ***δ*s** = (0.05,0.0005,0)^T^ when the monomorphic resident **s** has sufficiently converged toward **s**^∗^ to satisfy |**s** ™ **s**^∗^| < 0.0001, indicated with an orange triangle). The right most panel shows dynamics of population fractions. Simulations ended when |**s**_1_ – **s**_2_| ≥ 20 has been attained (dimorphic residents are indicated with small circles) or when either resident has gone extinct (the extinct resident is indicated with cross marks). Panel (d) shows evolutionary dynamics simulated with the stochastic method used in figure 1, under the same parameter values with those for panel (c). Parameters: 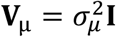 with *σ*_*μ*_ = 0.005, 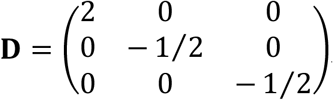, and 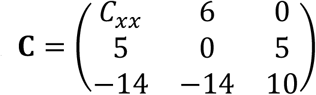 with 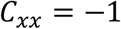 for (a), 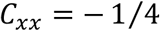 for (b), and 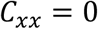 for (c) and (d).

**Figure 4.**
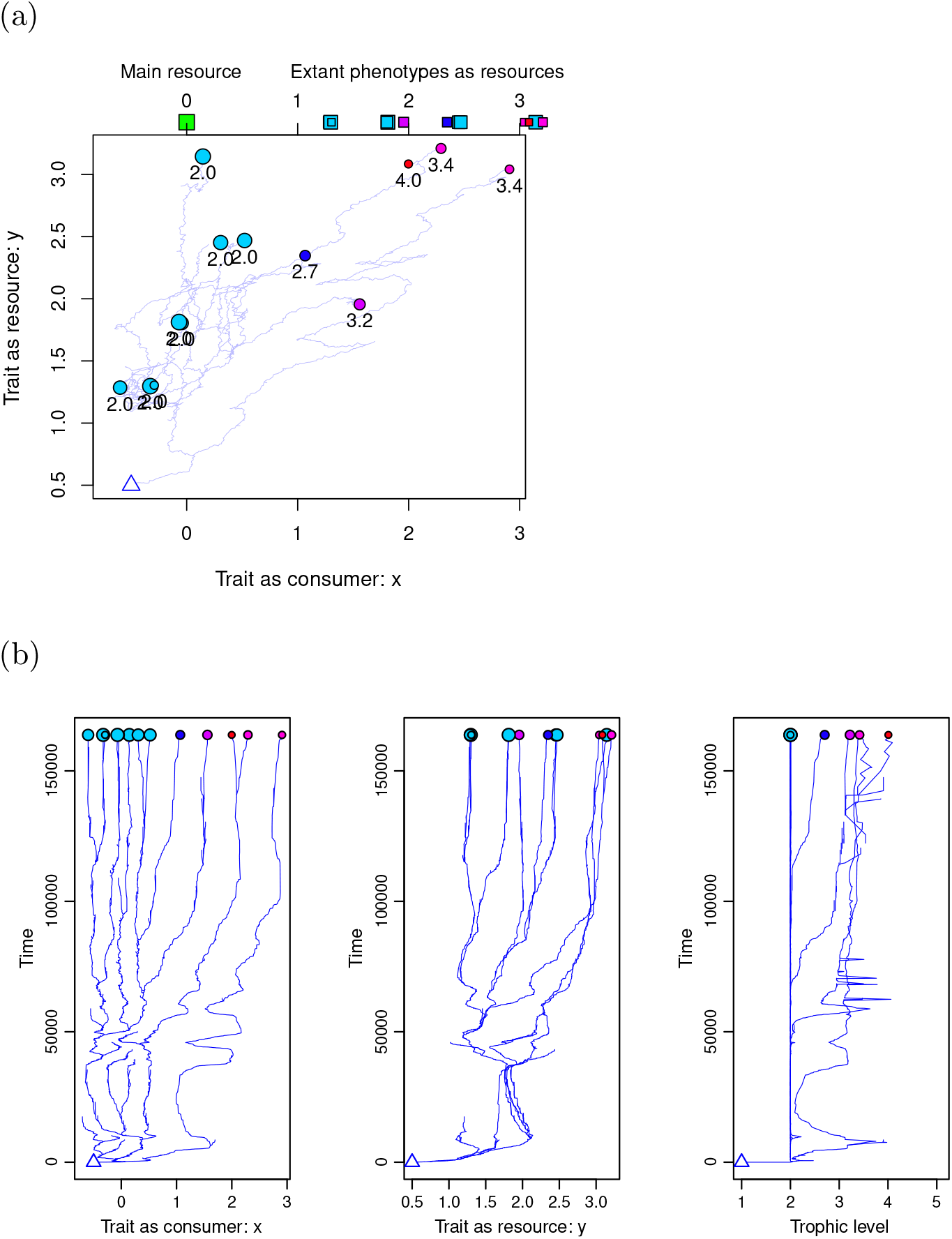
Numerically simulated evolution through resource competition and cannibalistic interaction in the model of an application example. Evolution is simulated under the assumption of asexual reproduction and rare mutation (see Section 3). Panel (a) shows evolutionary trajectories of resident phenotypes in the two-dimensional trait space ( *x, y*), where each small circle indicates existing phenotypes at the end of simulation. Each circle’s color and the number below it indicate the corresponding phenotype’s trophic level calculated by assigning zero to the trophic level of the main resource (see Appendix S7.1). Panel (b) shows time changes of residents’ trait *x*, trait *y*, and their trophic levels. Parameters: *σ*_*μ*_ = 0.005, *σ*_*α*_ = *σ*_*β*_ = 0.3, and *ρ* = *c* = 0.5.

Below, we analyze the initial evolutionary branching of a monomorphic population. For simplicity in this application example, we assume 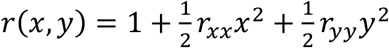 with constants *r*_*xx*_ and *r*_*yy*_. The invasion fitness for a mutant **s**′ = (*x*′, *y*′)^T^ against a resident **s** = (*x, y*)^T^ is derived as

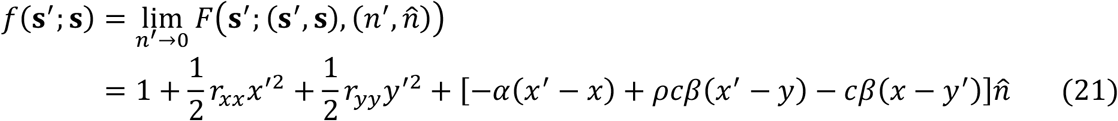

with 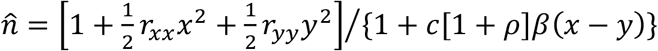 being the equilibrium population size for **s** = (*x, y*)^T^. Then, from the fitness gradient for the resident, given by

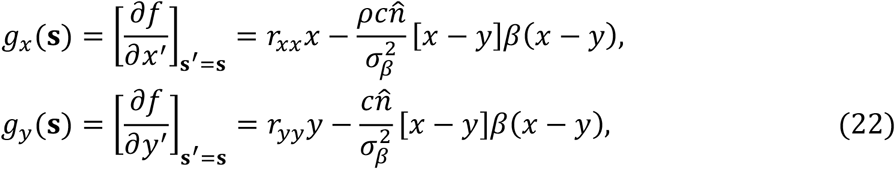

we find an evolutionarily singular point at the origin, i.e., **s**^∗^ = (0,0)^T^. By applying equations (6) and (11) for **s**^∗^ = (0,0)^T^, we obtain

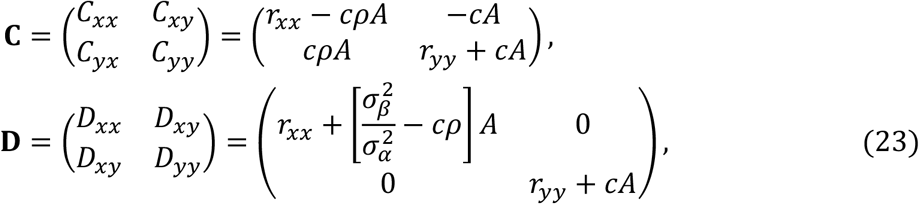

where 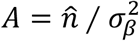 is always positive.

Then, from equations (22) and table 1, the conditions for **s**^∗^ being a possible branching point are given by

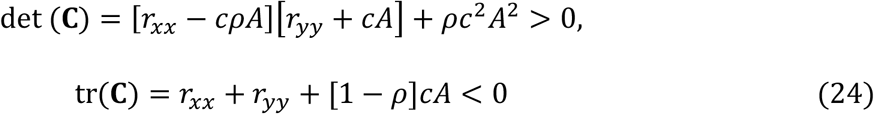

(for convergence stability), and

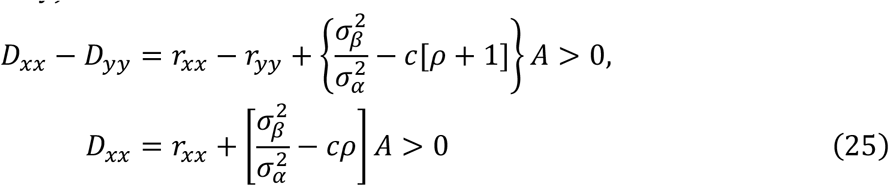

(for evolutionary instability and dimorphic emergence). The conditions for **s**^∗^ being a strongly possible branching point are given by

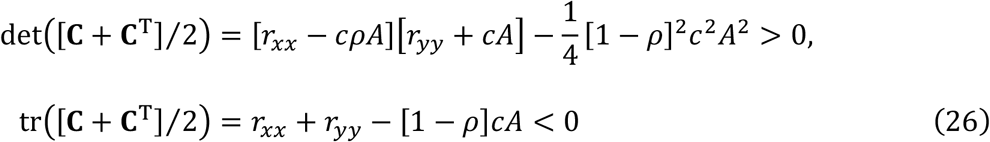

(for strong convergence stability), and equation (25). The conditions for **s**^∗^ being an inevitable branching point are given by

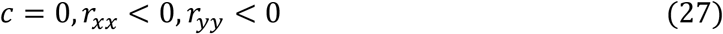

(for absolute convergence stability), and

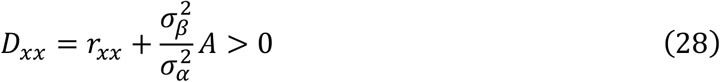

(for evolutionary instability).

The dependencies of these branching point conditions on *r*_*xx*_, *r*_*yy*_, and *c* are shown in figure 5. Numerically simulated evolutionary branchings induced by these branching points are shown in figures 6 and 7. Notably, possible branching points can induce evolutionary branching even when they lack convergence stability along the direction of dimorphic divergence given by the *x*-axis, i.e., *C*_*xx*_ = *r*_*xx*_ − *cρA* > 0 (figures 6a and 7a, which are similar to figures 1a and 2a). In this case, the coevolution of traits *x* and *y* through cannibalistic interaction is essential for the convergence stability of **s**^∗^ and hence for the occurrence of evolutionary branching.

**Figure 5.**
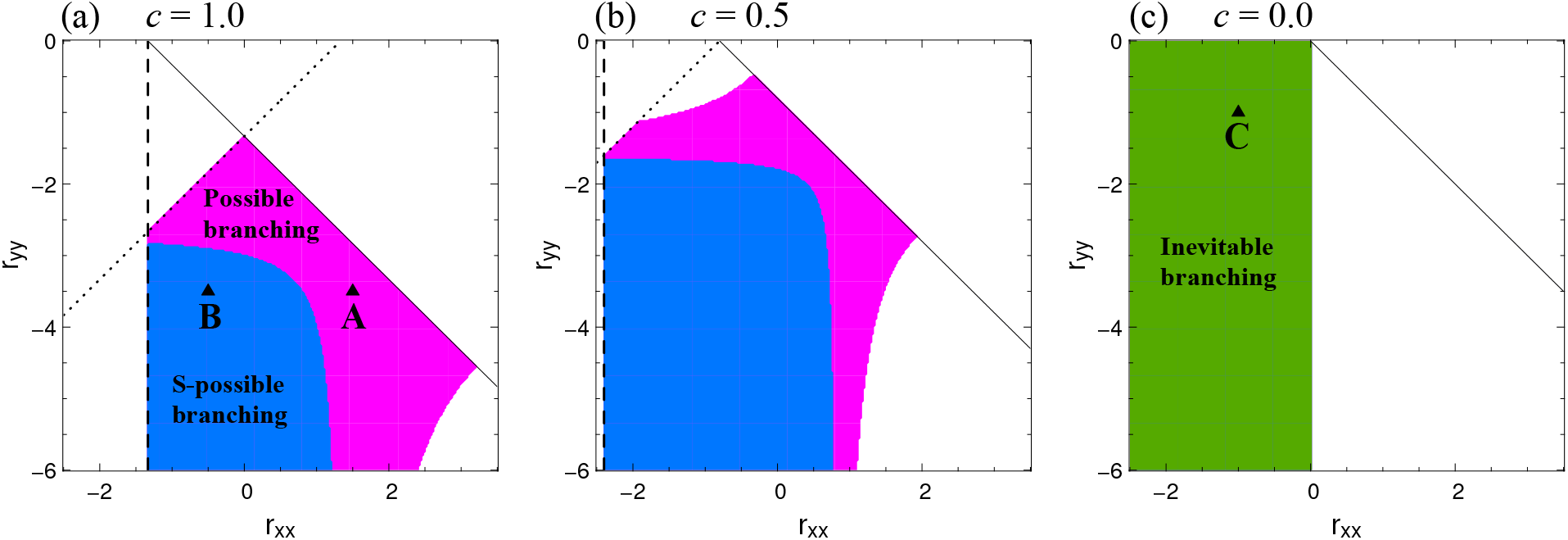
Parameter dependency of three classes of evolutionary branching points in the model of an application example (Section 4.4). Parameters: *σ*_*μ*_ = 0.005, *σ*_*α*_ = *σ*_*β*_ = 0.5, and *ρ* = 0.5 for all panels (a-c); *c* = 1, 0.5, and 0 for panels (a), (b), and (c), respectively; s *r*_*xx*_, *r*_*yy*_t = ( 1.5, ™3.5), (™0.5, ™3.5), and (™1, ™1) for “**A**”, “**B**”, and “**C**”, respectively.

**Figure 6.**
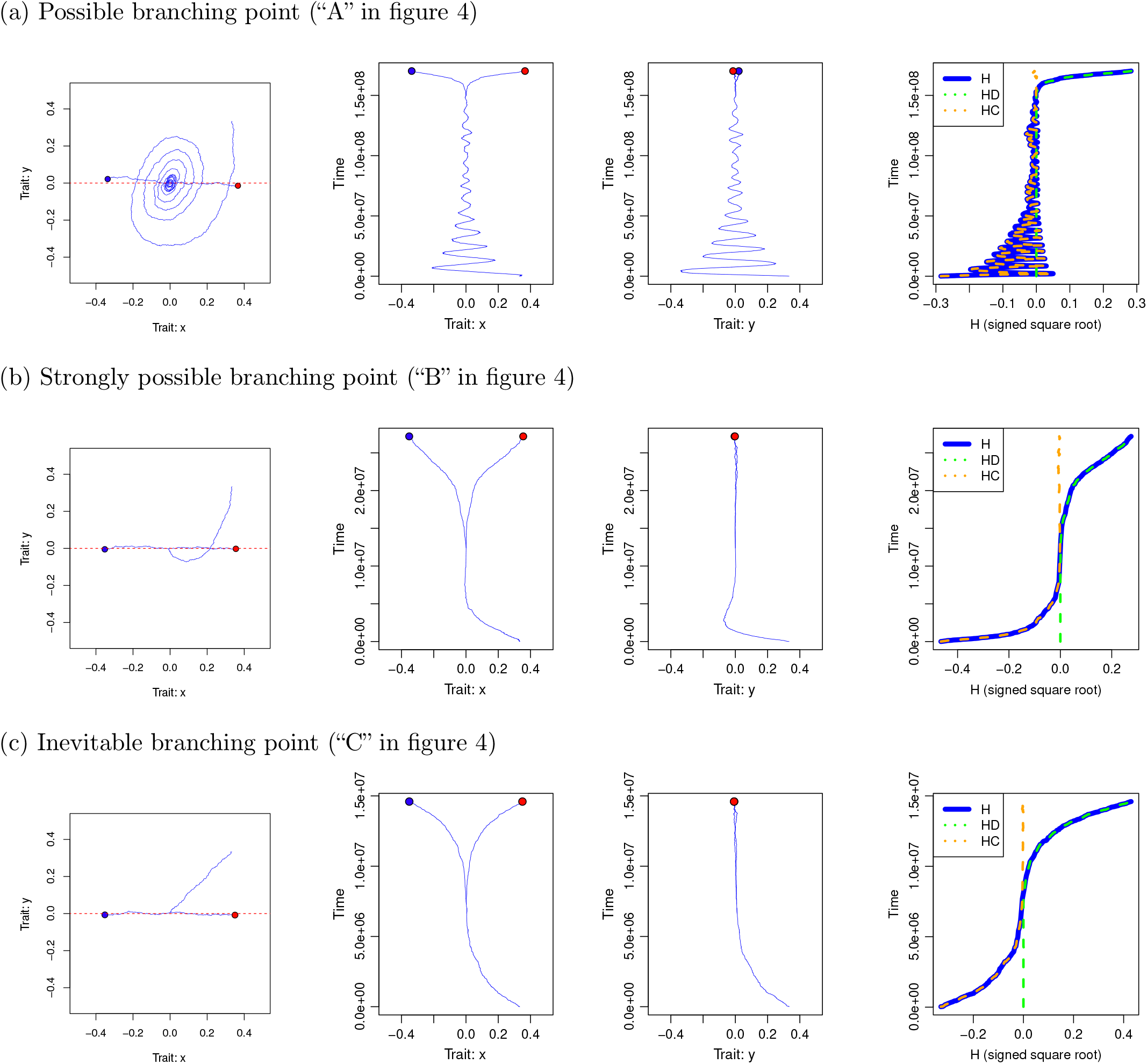
Evolutionary branching in simulated evolution under the assumption of asexual reproduction and rare mutation, induced by a possible, strongly possible, or inevitable branching point in the model of an application example (Section 4.4). Evolutionary dynamics starting from an initial phenotype **s**_0_ = (0.4,0.4)^T^ was simulated in the same manner as those shown in figure 1, except that the fitness function was given by equation (19). Parameter values for (a), (b) and (c) are the same with those for “**A**”, “**B**”, and “**C**” in figure 5.

**Figure 7.**
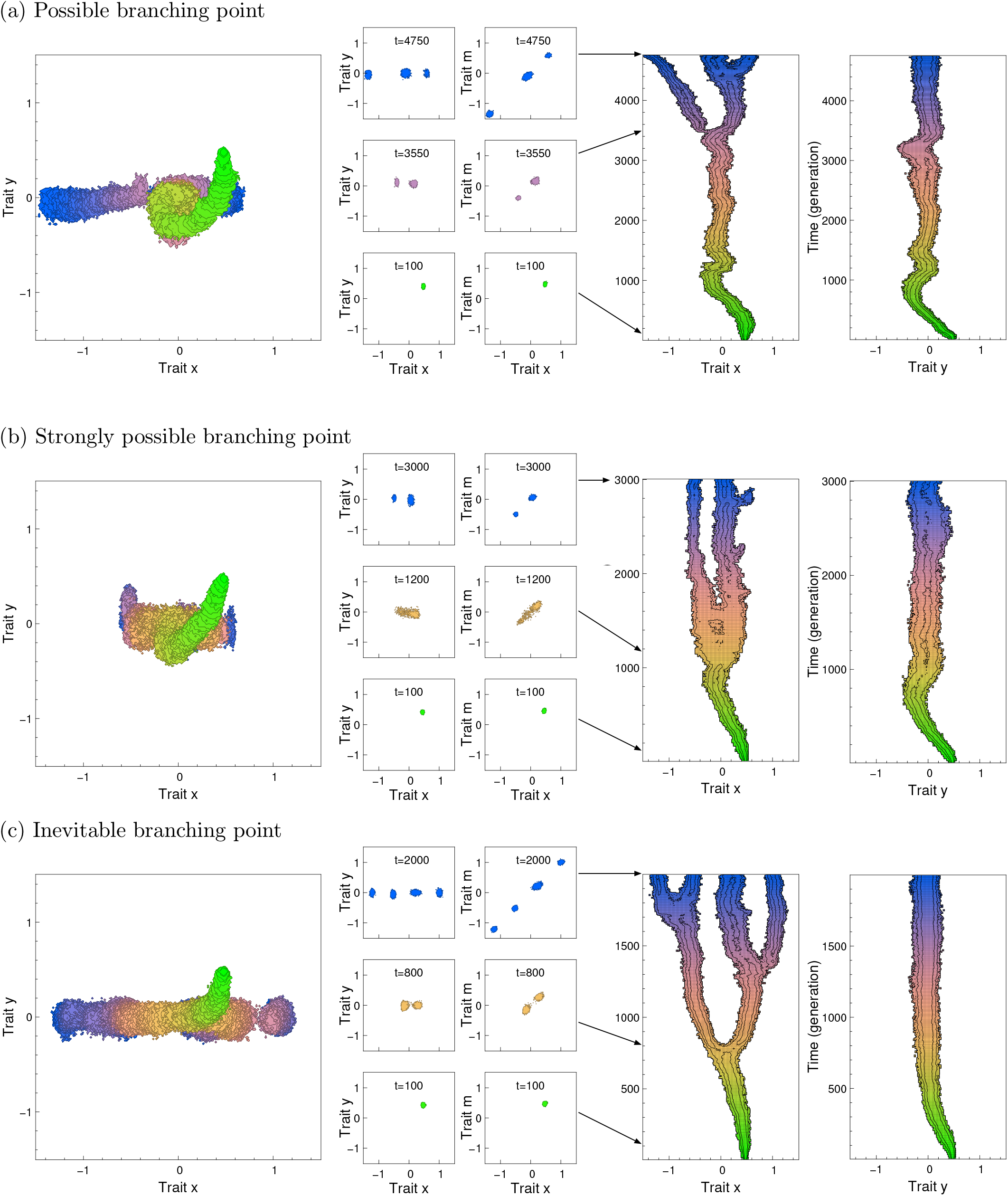
Evolutionary branching in simulated evolution under the assumption of sexual reproduction and non-rare mutation, for the same ecological settings with panels (a-c) in figure 6. Plotting formats are the same as in figure 2, except that the black contours in the left two panels indicate 10 and 100 individuals. The evolutionary dynamics was simulated in the same manner with the evolutionary dynamics shown in figure 2. Parameters for reproduction and mutation: *b*(**s**) = 1, *d*(**s**) = 1 ™ *F*(**s**; (**s**_1_, …, **s**_*M*_), (*n*_1_, …, *n*_*M*_)/*K*_0_) with *K*_0_ = 2000, number of loci (diploid): 30, mutation rate per locus (stepwise mutation): 5 × 10^-4^.

In figure 5, a large value of *c* (intense cannibalistic interaction relative to the resource competition) results in a large parameter region for possible branching points that are not strongly possible branching points (colored purple). This is because a large *c* causes a large |*C*_*xy*_ − *C*_*yx*_| through the interactions in the fitness function between mutant’s trait *x* and resident’s trait *y* and between the mutant’s trait *y* and the resident’s trait *x*. In general, this kind of relationship can be expected when two different kinds of traits interactively contribute to the fitness function so that 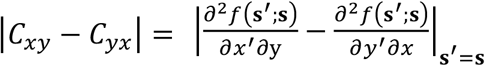 is large (e.g., the contribution of weapons and armors, or that of optical (or sonic) sensors and hiding strategies, in predator-prey interactions or in fighting for territories).

## 5 Discussion

### 5.1 Conditions for evolutionary branching points in multi-dimensional trait spaces

In one-dimensional trait spaces, the likelihood of evolutionary branching for a monomorphic population is well characterized by whether the spaces have convergence stable non-ESSes, called evolutionary branching points (Metz et al. 1996; Geritz et al. 1997). In trait spaces with arbitrary higher dimensions, the conditions for the existence of points that guarantee the entire process of evolutionary branching (consisting of monomorphic convergence, dimorphic emergence, and dimorphic divergence (Ito and Dieckmann 2014)) have not yet been established, although considerable progress has been made (e.g., Geritz et al. 2016). So far, strongly convergence stable non-ESSes have been treated as candidates for evolutionary branching points (Ackermann and Doebeli 2004; Ito and Shimada 2007; Ito and Dieckmann 2012, 2014; Geritz et al. 2016; Ito and Sasaki 2016, 2020).

In the present study, we analyzed in multi-dimensional trait spaces whether convergence stable non-ESSes can induce evolutionary branching, by developing conditions for branching possibility (based on the canonical equation) and branching inevitability (based on a time-increasing function related to the local Lyapunov function). Consequently, we showed that in trait spaces of arbitrary dimensions the branching possibility is warranted not only for strongly convergence-stable non-ESSes, but also for convergence-stable non-ESSes that are not strongly convergence stable yet satisfy the conditions for the emergence of dimorphism and their locally stable divergence (Table 1). We also showed that all absolutely convergence stable non-ESSes satisfy the branching inevitability conditions; they have time-increasing functions that guarantee the steady progression of evolutionary branching. These analytical results showed good agreement with likelihoods of evolutionary branching in numerically simulated evolution.

### 5.2 Branching possibility conditions

The branching possibility conditions were developed by describing evolutionary dynamics as trait substitution sequences, under the basic assumption of adaptive dynamics theory that mutation is sufficiently rare so that the timescales of population dynamics and evolutionary dynamics are completely separated. This description enabled decomposition of evolutionary branching into the deterministically describable processes of monomorphic convergence and dimorphic divergence, interconnected by the stochastic process of dimorphic emergence and their initial divergence. The two deterministically describable processes, i.e., monomorphic convergence and dimorphic divergence, were analyzed by applying the canonical equation with respect to their monotonicity. The stochastic process connecting them was analyzed directly as trait substitution sequences with respect to whether such a connection has non-zero probability. In contrast to this stochastic treatment of the initial divergence, Geritz et al. (2016) deterministically described the initial divergent evolution of polymorphic residents with the coupled Lande’s equations (Lande 1979), by assuming that the timescales of population dynamics and evolutionary dynamics are not separated. They derived a set of conditions guaranteeing that the initial divergent evolution of slightly differentiated morphs in a trait space of an arbitrary dimension result in dimorphic divergence without collapse into a single morph. These conditions are satisfied by strongly convergence stable non-ESSes in two-dimensional trait spaces, yet the higher-dimensional cases remain to be analyzed further (Geritz et al. 2016). It is notable that the conditions derived by Geritz et al. (2016) (i.e., equation (29) in their paper) are different from but structurally similar to our conditions for locally stable dimorphic divergence, i.e., the fourth formula of equation (15). Notably, both kinds of conditions imply that dimorphic divergence proceeds without collapse when the divergence is faster than the deviation of the intermediate phenotype of dimorphic residents from the singular point.

### 5.3 Branching inevitability conditions

Regarding the branching inevitability conditions, the existence of a time-increasing function guaranteeing evolutionary branching, shown in equation (18), implies that the dynamics of evolutionary branching can be considerably deterministic. Because the branching inevitability does not depend on the dimensionality of the trait space, the conditions are applicable not only to multi-dimensional trait spaces but also to one-dimensional ones.

Note that equation (18) gives a locally approximated representation of Fisher’s fundamental theorem of natural selection (Baez 2021), as shown in Appendix S4.1. Note also that the population dynamics given by equation (14) under **C**^T^ = **C** corresponds to a class of evolutionary games called potential games (Hofbauer and Sigmund 1998), which are known as to have potential functions that monotonically increase through replicator dynamics, corresponding to equation (18) in the present paper (see Appendix S4.5 for details).

Although absolutely convergence stability required for the branching inevitability is restrictive, this stability might be more common among eco-evolutionary models than previously thought (Leimar 2009), because convergence stable points are always absolutely convergence stable in an arbitrary trait space consisting of a one-dimensional trait under frequency-dependent selection and of one or more traits under frequency-independent selection (as in the case of *c* = 0 in Section 4.4).

### 5.4 Limitations and future directions

Note that the conditions for branching possibility and inevitability both assume asexual populations, although their usefulness for sexual populations is shown by numerical simulation (figures 2 and 7). Note also that the derivation of branching possibility conditions assumes timescale separation between population dynamics and evolutionary dynamics (i.e., rare mutation such that evolutionary dynamics can be treated as trait substitution sequences), whereas the derivation of the branching inevitability conditions does not. Hence, the branching inevitability conditions are applicable even under incomplete timescale separation between population dynamics and evolutionary dynamics (due to non-rare mutation or weak selection), as long as the population dynamics of existing phenotypes are deterministically described with the approximate fitness function, equation (14). However, the branching inevitability conditions require the approximate symmetry of matrix **C** (i.e, absolute convergence stability), which would be considerably restrictive, especially in high-dimensional trait spaces. This requirement for **C** might be relaxed in a future study, by analyzing the dynamics of the long-term average of function *H* instead of *H* itself in equation (18). Contrastingly, the branching possibility conditions may be satisfied by all classes of convergence stable points. As a future step, the branching point conditions might be extended to make a quantitative prediction of branching likelihood, by treating the stochastic process connecting two deterministically describable processes (monomorphic convergence and dimorphic divergence) as the dynamics of a probability distribution for existing phenotypes.

