## Appendices for "Evolutionary branching points in multi-dimensional trait spaces"

Throughout the text, we use italics for denoting scalars, bold lower case for column vectors, and bold upper case for matrices.

### S1. Simulation algorithm

#### S1.1. Trait substitution sequence

Evolutionary dynamics were simulated mainly as trait substitution sequences with an algorithm called the oligomorphic stochastic model (OSM) (Ito and Dieckmann, 2007; Ito and Dieckmann, 2014; Ito and Sasaki, 2020; Ito and Sasaki, 2023) that assumes asexual reproduction and rare mutation. The OSM in Ito and Sasaki (2020) and Ito and Sasaki (2023) is a slightly simplified version of the original OSM by Ito and Dieckmann (2007). The simulation in our present analysis followed Ito and Sasaki (2020), as described below. The probability density for a mutant emergence with phenotype $\mathbf{s}'$ is proportional to $\mu \sum_{j=1}^{N} P_\mu(\mathbf{s}' - \mathbf{s}_i)\ \hat{n}_j$, where $P_\mu(\mathbf{s}' - \mathbf{s}_j)$ is a mutation distribution describing the probability density distribution for the emergence of a mutant $\mathbf{s}'$ from its parental phenotype $\mathbf{s}_j$ when a mutant emerges, defined in equation (S2.81) in Section S2.7. The waiting time for a mutant emergence follows the exponential distribution with its expected value $1/[\mu \sum_{j=1}^{N} \hat{n}_j]$. The probability of successful invasion is given by the invasion fitness of the mutant (Dieckmann and Law, 1996). When a mutant has successfully invaded into the system, the next population-dynamical equilibrium is calculated by the time integration of the equation for population dynamics, with the initial population size for the mutant being set to a positive constant $\varepsilon_{\text{new}}$. During the population dynamics, a resident phenotype (or the mutant) is removed when its population size becomes smaller than an extinction threshold $\varepsilon_{\text{ext}}$ (see Ito and Sasaki [2020] for details). $\varepsilon_{\text{ext}} = 1.0 \times 10^{-6}$ and $\varepsilon_{\text{new}} = 10\,\varepsilon_{\text{ext}}$ were used for all simulations with the OSM shown in this paper.

#### S1.2. Individual-based stochastic simulation

We also simulated the evolutionary dynamics under sexual reproduction and small but nonrare mutation (Figs. 2 and 6) following Ito and Dieckmann (2007) (see also Ito and Sasaki, 2023). This algorithm considers male and female individuals, diploid inheritance, and quantitative traits based on multiloci. Specifically, $L_x$ and $L_y$ loci are considered for traits $x$ and $y$, respectively, with integer allelic values and with the value of $x$ and $y$ given by the averages of allelic values across loci. For recombination, $C_x$ and $C_y$ linkage clusters are considered for $x$ and $y$, respectively, with no recombination within a cluster and with free recombination between clusters. Individual birth and death rates are defined so that the expected population dynamics is described with equation (1) in the main text. Specifically, the $k$th female individual produces offspring individuals at rate $b_k = 2[F_k + 1]$ with $F_k$ describing the $k$th female's fitness, and dies at rate $d_k = 1$, where $N_I$ describes the total number of individuals in the system. The male individuals do not produce offsprings, and the $l$th male individual dies at rate $d_l = 1$. For offspring production, a female individual $k$ chooses a male partner $l$ for mating with probability

$$\tilde{P}_{kl} = \frac{P_{kl}\, b_l}{\sum_{l'} P_{kl'}\, b_{l'}}, \tag{S1.1}$$

where $b_l = F_l + 1$ denotes the potential of male individual $l$ for contribution to offspring production. $P_{kl}$ describes the visibility of the male $l$ for the female $k$, depending on a scalar mating trait $m_l$ in male $l$ and on trait $x$ in female $k$. $m_l$ has $L_m$ loci each of which contains an integer value, and those loci are organized into $C_m$ linkage clusters. The mating visibility function $P_{kl}$ is then given by

$$P_{kl} = \exp\left( -\frac{|m_l - \tilde{m}(\mathbf{s}_k)|^2}{2\,\sigma_{\text{mating}}^2} \right), \tag{S1.2}$$

with $\tilde{m}(\mathbf{s}_k)$ describing the most outstanding mating trait for a female of phenotype $\mathbf{s}_k$, which is defined as $\tilde{m}(\mathbf{s}_k) = x_k$ for simplicity. Allelic mutations that increase or decrease allelic values by 1 (which happens with equal probability) occur with per locus probabilities of $\tilde{\mu}_x$, $\tilde{\mu}_y$, and $\tilde{\mu}_m$ for each birth of an individual with its sex assigned randomly. For the simulation, the above algorithm was implemented in C++. s

#### S1.3. Evolutionary branching in individual-based stochastic simulation under asexual reproduction

The algorithm described in Section S3.2 can also be used for asexual populations, by assuming that individuals are all females reproducing clonally (i.e., $b_k = F_k + 1$ and $d_k = 1$). Figures S1 and

S2 show the simulated evolutionary branching of asexual populations, corresponding to figures 2f and 6, respectively, in the main text.

(a) Possible branching point

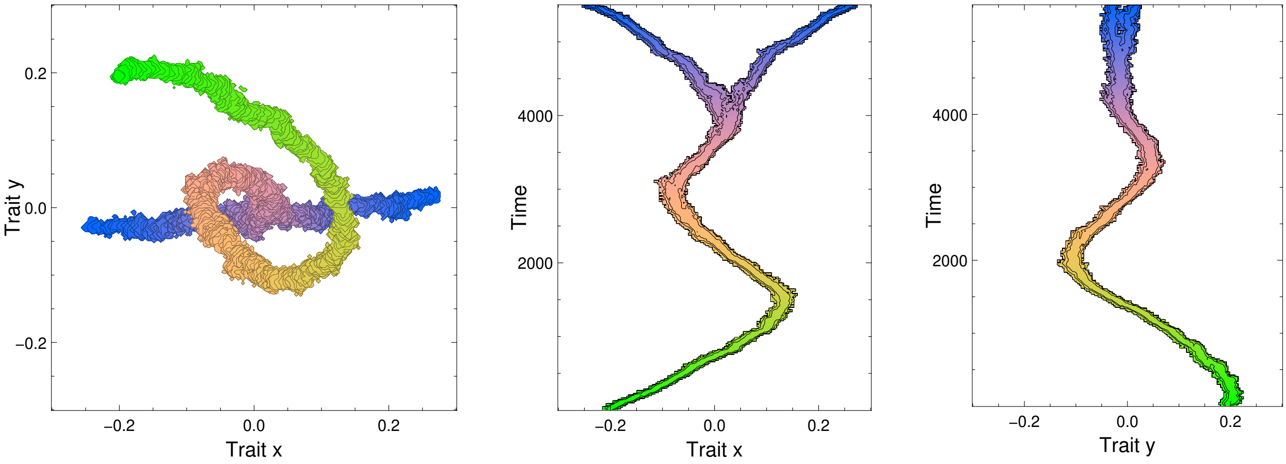

(b) Strongly possible branching point

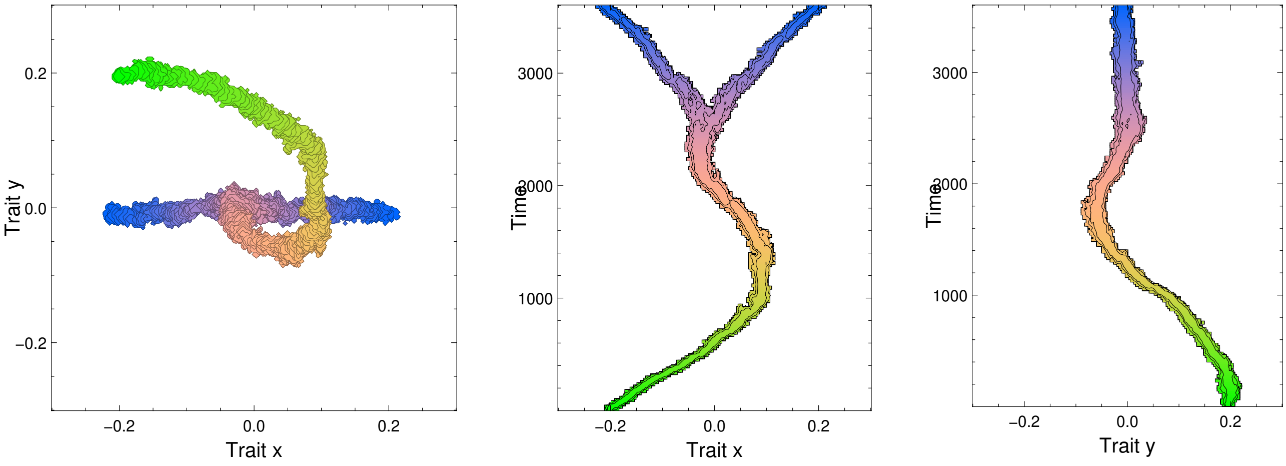

(c) Inevitable branching point

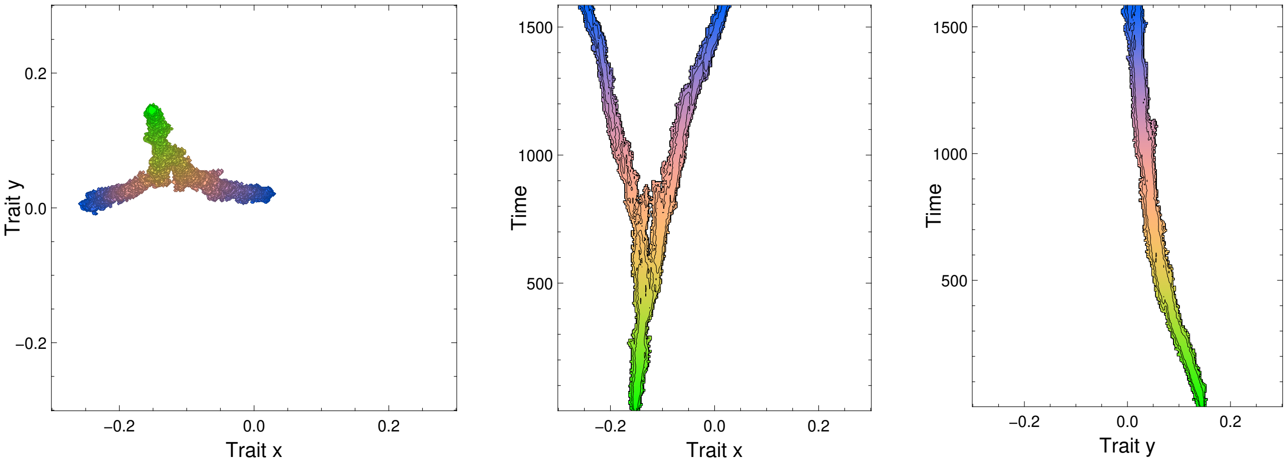

**Figure S1.** Evolutionary branching simulated under assumptions of asexual reproduction and nonrare mutation, under the same ecological parameter sets with those for figure 2 in the main text.

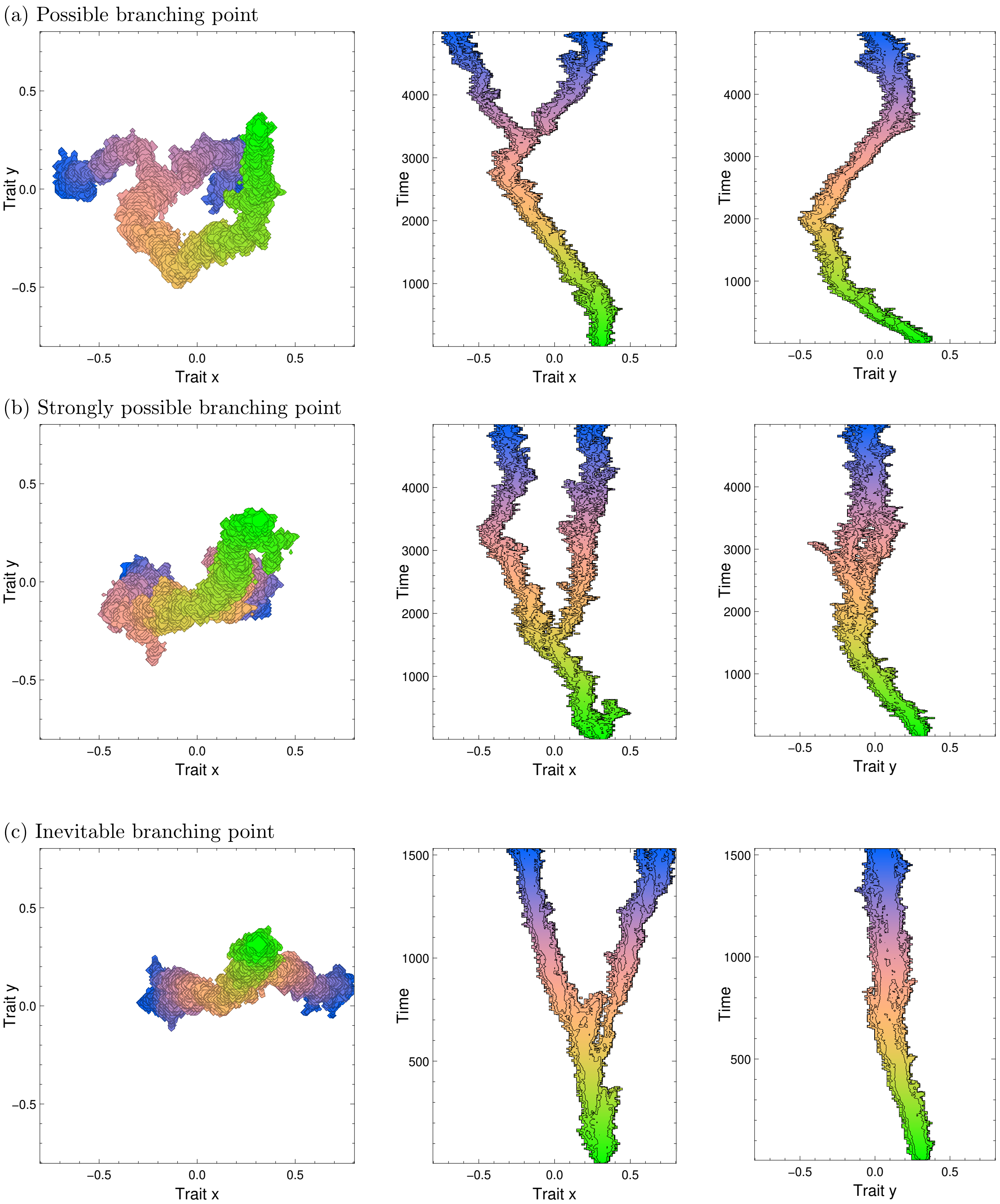

**Figure S2.** Evolutionary branching simulated under assumptions of asexual reproduction and nonrare mutation, under the same ecological parameter sets with those for figure 6 in the main text.

### S2. Derivation of equation (14): Approximate fitness function

According to Meszena et al. (2005), if $M$ coexisting phenotypes, denoted by $\mathbf{s}_1, ..., \mathbf{s}_M$, are similar to each other and are close to a point $\mathbf{s}_0$ so that $|\mathbf{s}_i-\mathbf{s}_0|<\varepsilon$ holds for all $i=1,...,M$ (where $\varepsilon$ is a positive and sufficiently small constant), then their fraction dynamics fueled by mutant invasions satisfy

$$\begin{aligned}\frac{\mathrm{dln}\,p_i}{\mathrm{dt}}-\frac{\mathrm{dln}\,p_j}{\mathrm{dt}} &= \mathbf{f}_{\mathbf{s}'}^T[\mathbf{s}_i-\mathbf{s}_j]+\frac{1}{2}[\mathbf{s}_i-\mathbf{s}_0]^T\,\mathbf{F}_{\mathbf{s}'\mathbf{s}'}[\mathbf{s}_i-\mathbf{s}_0]-\frac{1}{2}[\mathbf{s}_j-\mathbf{s}_0]^T\,\mathbf{F}_{\mathbf{s}'\mathbf{s}'}[\mathbf{s}_j-\mathbf{s}_0]\\ &\quad+[\bar{\mathbf{s}}-\mathbf{s}_0]^T\,\mathbf{F}_{\mathbf{s}\mathbf{s}'}[\mathbf{s}_i-\mathbf{s}_j]+O(\varepsilon^3)\end{aligned} \tag{S2.1}$$

for all combinations of $i,j=1,...,M$, where $\mathbf{f}_{\mathbf{s}'}$, $\mathbf{F}_{\mathbf{s}'\mathbf{s}'}$, and $\mathbf{F}_{\mathbf{s}\mathbf{s}'}$ are derivatives of the invasion fitness $f(\mathbf{s}';\mathbf{s})$ for $\mathbf{s}'$ against monomorphic resident $\mathbf{s}$:

$$\begin{aligned}\mathbf{f}_{\mathbf{s}'} &= \left[\frac{\partial f(\mathbf{s}';\mathbf{s})}{\partial\mathbf{s}'}\right]_{\mathbf{s}'=\mathbf{s}=\mathbf{s}_0}=\begin{pmatrix}\frac{\partial f(\mathbf{s}';\mathbf{s})}{\partial x_1'}\\ \vdots\\ \frac{\partial f(\mathbf{s}';\mathbf{s})}{\partial x_L'}\end{pmatrix}_{\mathbf{s}'=\mathbf{s}=\mathbf{s}_0},\\ \mathbf{F}_{\mathbf{s}'\mathbf{s}'} &= \begin{pmatrix}\frac{\partial^2 f(\mathbf{s}';\mathbf{s})}{\partial x_1'^2} & \cdots & \frac{\partial^2 f(\mathbf{s}';\mathbf{s})}{\partial x_1'\partial x_L'}\\ \vdots & \ddots & \vdots\\ \frac{\partial^2 f(\mathbf{s}';\mathbf{s})}{\partial x_1'\partial x_L'} & \cdots & \frac{\partial^2 f(\mathbf{s}';\mathbf{s})}{\partial x_L'\partial x_L'}\end{pmatrix}_{\mathbf{s}'=\mathbf{s}=\mathbf{s}_0},\\ \mathbf{F}_{\mathbf{s}\mathbf{s}'} &= \begin{pmatrix}\frac{\partial^2 f(\mathbf{s}';\mathbf{s})}{\partial x'\partial x} & \cdots & \frac{\partial^2 f(\mathbf{s}';\mathbf{s})}{\partial x_1'\partial x_L}\\ \vdots & \ddots & \vdots\\ \frac{\partial^2 f(\mathbf{s}';\mathbf{s})}{\partial x_L'\partial x_1} & \cdots & \frac{\partial^2 f(\mathbf{s}';\mathbf{s})}{\partial x_L'\partial x_L}\end{pmatrix}_{\mathbf{s}'=\mathbf{s}=\mathbf{s}_0}.\end{aligned} \tag{S2.2}$$

Without loss of generality, we assume $\mathbf{s}_0=\mathbf{0}$ and expand $f(\mathbf{s}';\mathbf{s})$ at $\mathbf{s}_0=\mathbf{0}$ with respect to $\mathbf{s}'$ and $\mathbf{s}$ as

$$\begin{aligned}f(\mathbf{s}';\mathbf{s}) &= \sum_{i=1}^{L}\frac{\partial f}{\partial x_i'}x_i'+\sum_{j=1}^{L}\frac{\partial f}{\partial x_j}x_j\\ &\quad+\frac{1}{2}\left[\sum_{i=1}^{L}\frac{\partial^2 f}{\partial x_i'^2}x_i'^2+\sum_{j=1}^{L}\frac{\partial^2 f}{\partial x_j^2}x_j^2+2\sum_{i=1}^{L}\sum_{j=1}^{L}\frac{\partial^2 f}{\partial x_i'\partial x_j}x_i'x_j\right]+O(\varepsilon^3),\end{aligned} \tag{S2.3}$$

where $\frac{\partial f}{\partial x_i'}$, $\frac{\partial f}{\partial x_j}$, $\frac{\partial^2 f}{\partial x_i'^2}$, $\frac{\partial^2 f}{\partial x_j^2}$, and $\frac{\partial^2 f}{\partial x_i'\partial x_j}$ describe the first and second derivatives of $f(\mathbf{s}',\mathbf{s})$ evaluated at $\mathbf{s}'=\mathbf{s}=\mathbf{s}_0$. Because $f(\mathbf{s};\mathbf{s})=0$ must hold for any $\mathbf{s}$, we require $\frac{\partial f}{\partial x_i'}+\frac{\partial f}{\partial x_i}=0$ and $\frac{\partial^2 f}{\partial x_i'^2}+2\,\frac{\partial^2 f}{\partial x_i'\partial x_j}+\frac{\partial^2 f}{\partial x_j^2}=0$ for any $i,j=1,...,L$, which upon substitution into equation (S1.3) gives

$$\begin{aligned}f(\mathbf{s}';\mathbf{s}) &= \mathbf{f}_{\mathbf{s}'}^T[\mathbf{s}'-\mathbf{s}]+\mathbf{s}^T[\mathbf{F}_{\mathbf{s}'\mathbf{s}'}+\mathbf{F}_{\mathbf{s}\mathbf{s}'}][\mathbf{s}'-\mathbf{s}]+\frac{1}{2}[\mathbf{s}'-\mathbf{s}]^T\,\mathbf{F}_{\mathbf{s}'\mathbf{s}'}[\mathbf{s}'-\mathbf{s}]+O(\varepsilon^3)\\ &= \mathbf{g}^T[\mathbf{s}'-\mathbf{s}]+\mathbf{s}^T\,\mathbf{C}[\mathbf{s}'-\mathbf{s}]+\frac{1}{2}[\mathbf{s}'-\mathbf{s}]^T\,\mathbf{D}[\mathbf{s}'-\mathbf{s}]+O(\varepsilon^3)\end{aligned} \tag{S2.4}$$

with

$$\begin{aligned}\mathbf{g}(\mathbf{s}) &= \left[\frac{\partial f(\mathbf{s}';\mathbf{s})}{\partial\mathbf{s}'}\right]_{\mathbf{s}'=\mathbf{s}}=\mathbf{f}_{\mathbf{s}'}+[\mathbf{F}_{\mathbf{s}'\mathbf{s}'}+\mathbf{F}_{\mathbf{s}\mathbf{s}'}]^T\,\mathbf{s},\\ \mathbf{g} &= \mathbf{g}(\mathbf{0})=\mathbf{f}_{\mathbf{s}'},\\ \mathbf{C} &= \left[\frac{\partial\mathbf{g}(\mathbf{s})^T}{\partial\mathbf{s}}\right]_{\mathbf{s}=\mathbf{0}}=\mathbf{F}_{\mathbf{s}'\mathbf{s}'}+\mathbf{F}_{\mathbf{s}\mathbf{s}'},\\ \mathbf{D} &= \begin{pmatrix}\frac{\partial^2 f(\mathbf{s}';\mathbf{s})}{\partial x'\partial x} & \cdots & \frac{\partial^2 f(\mathbf{s}';\mathbf{s})}{\partial x_1'\partial x_L}\\ \vdots & \ddots & \vdots\\ \frac{\partial^2 f(\mathbf{s}';\mathbf{s})}{\partial x_L'\partial x_1} & \cdots & \frac{\partial^2 f(\mathbf{s}';\mathbf{s})}{\partial x_L'\partial x_L}\end{pmatrix}_{\mathbf{s}'=\mathbf{s}=\mathbf{0}}=\mathbf{F}_{\mathbf{s}'\mathbf{s}'}\end{aligned} \tag{S2.5}$$

(see also Ito and Sasaki (2020) for the derivation), which upon substitution into equation (S1.1) gives

$$\begin{aligned}
\frac{\mathrm{d}\ln p_i}{\mathrm{dt}} - \frac{\mathrm{d}\ln p_j}{\mathrm{dt}} &= \mathbf{g}^T[\mathbf{s}_i - \mathbf{s}_j] + \frac{1}{2}\mathbf{s}_i^T\,\mathbf{D}\,\mathbf{s}_i - \frac{1}{2}\mathbf{s}_j^T\,\mathbf{D}\,\mathbf{s}_j + \bar{\mathbf{s}}^T[\mathbf{C}-\mathbf{D}][\mathbf{s}_i - \mathbf{s}_j] + O(\varepsilon^3)\\
&= \left\{\mathbf{g}^T[\mathbf{s}_i - \bar{\mathbf{s}}] + \bar{\mathbf{s}}^T\,\mathbf{C}[\mathbf{s}_i - \bar{\mathbf{s}}] + \frac{1}{2}[\mathbf{s}_i - \bar{\mathbf{s}}]^T\,\mathbf{D}[\mathbf{s}_i - \bar{\mathbf{s}}]\right\}\\
&\quad -\left\{\mathbf{g}^T[\mathbf{s}_j - \bar{\mathbf{s}}] + \bar{\mathbf{s}}^T\,\mathbf{C}[\mathbf{s}_j - \bar{\mathbf{s}}] + \frac{1}{2}[\mathbf{s}_j - \bar{\mathbf{s}}]^T\,\mathbf{D}[\mathbf{s}_j - \bar{\mathbf{s}}]\right\}\\
&= f(\mathbf{s}_i, \bar{\mathbf{s}}) - f(\mathbf{s}_j, \bar{\mathbf{s}}) + O(\varepsilon^3).
\end{aligned} \tag{S2.6}$$

Hence, from equations (S2.5) and (S2.6) we see that

$$\begin{aligned}
\frac{\mathrm{d}\ln p_i}{\mathrm{dt}} &= \sum_{j=1}^{M} p_j \frac{\mathrm{d}\ln p_i}{\mathrm{dt}} = \sum_{j=1}^{M} p_j\left[\frac{\mathrm{d}\ln p_j}{\mathrm{dt}} + \{f(\mathbf{s}_i;\bar{\mathbf{s}}) - f(\mathbf{s}_j;\bar{\mathbf{s}}) + h.o.t.\}\right]\\
&= \sum_{j=1}^{M} \frac{dp_j}{\mathrm{dt}} + f(\mathbf{s}_i;\bar{\mathbf{s}}) - \sum_{j=1}^{M} p_j\, f(\mathbf{s}_j;\bar{\mathbf{s}}) + O(\varepsilon^3)\\
&= \left\{\mathbf{g}^T[\mathbf{s}_i - \bar{\mathbf{s}}] + \bar{\mathbf{s}}^T\,\mathbf{C}[\mathbf{s}_i - \bar{\mathbf{s}}] + \frac{1}{2}[\mathbf{s}_i - \bar{\mathbf{s}}]^T\,\mathbf{D}[\mathbf{s}_i - \bar{\mathbf{s}}]\right\}\\
&\quad -\sum_{j=1}^{M} p_j\left\{\mathbf{g}^T[\mathbf{s}_j - \bar{\mathbf{s}}] + \bar{\mathbf{s}}^T\,\mathbf{C}[\mathbf{s}_j - \bar{\mathbf{s}}] + \frac{1}{2}[\mathbf{s}_j - \bar{\mathbf{s}}]^T\,\mathbf{D}[\mathbf{s}_j - \bar{\mathbf{s}}]\right\} + O(\varepsilon^3)\\
&= \mathbf{g}^T[\mathbf{s}_i - \bar{\mathbf{s}}] + \bar{\mathbf{s}}^T\,\mathbf{C}[\mathbf{s}_i - \bar{\mathbf{s}}] + \frac{1}{2}[\mathbf{s}_i - \bar{\mathbf{s}}]^T\,\mathbf{D}[\mathbf{s}_i - \bar{\mathbf{s}}] - \frac{1}{2}\sum_{j=1}^{M} p_j[\mathbf{s}_j - \bar{\mathbf{s}}]^T\,\mathbf{D}[\mathbf{s}_j - \bar{\mathbf{s}}] + O(\varepsilon^3)\\
&= \mathbf{g}^T[\mathbf{s}_i - \bar{\mathbf{s}}] + \bar{\mathbf{s}}^T\,\mathbf{C}[\mathbf{s}_i - \bar{\mathbf{s}}] + \frac{1}{2}[\mathbf{s}_i - \bar{\mathbf{s}}]^T\,\mathbf{D}[\mathbf{s}_i - \bar{\mathbf{s}}] - \frac{1}{2}V_{\mathbf{D}} + O(\varepsilon^3)
\end{aligned} \tag{S2.7}$$

with $V_{\mathbf{D}} = \sum_{j=1}^{M} p_j[\mathbf{s}_j - \bar{\mathbf{s}}]^T\,\mathbf{D}[\mathbf{s}_j - \bar{\mathbf{s}}]$. From equation (S2.7) we get

$$\begin{aligned}
F(\mathbf{s}_i;(\mathbf{s}_1,\ldots,\mathbf{s}_M),(n_1,\ldots,n_M)) &= \frac{\mathrm{d}\ln n_i}{\mathrm{dt}} = \frac{\mathrm{d}\ln p_i\sum_{j=1}^{M} n_j}{\mathrm{dt}} = \frac{\mathrm{d}\ln p_i}{\mathrm{dt}} + \frac{1}{\sum_{k=1}^{M} n_k}\sum_{j=1}^{M}\frac{dn_j}{\mathrm{dt}}\\
&= \mathbf{g}^T[\mathbf{s}_i - \bar{\mathbf{s}}] + \bar{\mathbf{s}}^T\,\mathbf{C}[\mathbf{s}_i - \bar{\mathbf{s}}] + \frac{1}{2}[\mathbf{s}_i - \bar{\mathbf{s}}]^T\,\mathbf{D}[\mathbf{s}_i - \bar{\mathbf{s}}] - \frac{1}{2}V_{\mathbf{D}}\\
&\quad +\sum_{j=1}^{M} p_j\left[\frac{1}{n_j}\frac{dn_j}{\mathrm{dt}}\right] + O(\varepsilon^3)\\
&= \mathbf{g}^T[\mathbf{s}_i - \bar{\mathbf{s}}] + \bar{\mathbf{s}}^T\,\mathbf{C}[\mathbf{s}_i - \bar{\mathbf{s}}] + \frac{1}{2}[\mathbf{s}_i - \bar{\mathbf{s}}]^T\,\mathbf{D}[\mathbf{s}_i - \bar{\mathbf{s}}] - \frac{1}{2}V_{\mathbf{D}}\\
&\quad +\bar{F} + O(\varepsilon^3)
\end{aligned} \tag{S2.8}$$

with $\bar{F} = \sum_{j=1}^{M} p_j\, F(\mathbf{s}_j;(\mathbf{s}_1,\ldots,\mathbf{s}_M),(n_1,\ldots,n_M))$. Finally, we obtain

$$\begin{aligned}
F(\mathbf{s}';(\mathbf{s}_1,\ldots,\mathbf{s}_M),(n_1,\ldots,n_M)) &= \lim_{n'\to 0} F(\mathbf{s}';(\mathbf{s}_1,\ldots,\mathbf{s}_M,\mathbf{s}'),(n_1,\ldots,n_M,n'))\\
&= \mathbf{g}^T[\mathbf{s}' - \bar{\mathbf{s}}] + \bar{\mathbf{s}}^T\,\mathbf{C}[\mathbf{s}' - \bar{\mathbf{s}}] + \frac{1}{2}[\mathbf{s}' - \bar{\mathbf{s}}]^T\,\mathbf{D}[\mathbf{s}' - \bar{\mathbf{s}}] - \frac{1}{2}V_{\mathbf{D}}\\
&\quad +\bar{F} + O(\varepsilon^3).
\end{aligned} \tag{S2.9}$$

Additionally, we obtain the invasion fitness for $\mathbf{s}'$ as

$$\begin{aligned}
f(\mathbf{s}';\mathbf{s}_1,\ldots,\mathbf{s}_M) &= \lim_{n'\to 0} F(\mathbf{s}';(\mathbf{s}_1,\ldots,\mathbf{s}_M,\mathbf{s}'),(\hat{n}_1,\ldots,\hat{n}_M,n'))\\
&= \mathbf{g}^T[\mathbf{s}' - \bar{\mathbf{s}}] + \bar{\mathbf{s}}^T\,\mathbf{C}[\mathbf{s}' - \bar{\mathbf{s}}] + \frac{1}{2}[\mathbf{s}' - \bar{\mathbf{s}}]^T\,\mathbf{D}[\mathbf{s}' - \bar{\mathbf{s}}] - \frac{1}{2}V_{\mathbf{D}} + O(\varepsilon^3)
\end{aligned} \tag{S2.10}$$

with $\bar{\mathbf{s}} = \sum_{j=1}^{M}\hat{p}_j\,\mathbf{s}_j$ and $V_{\mathbf{D}} = \sum_{j=1}^{M}\hat{p}_j[\mathbf{s}_j - \bar{\mathbf{s}}]^T\,\mathbf{D}[\mathbf{s}_j - \bar{\mathbf{s}}]$ with $\hat{p}_j = \hat{n}_j/\sum_{i=1}^{M}\hat{n}_i$.

When $\mathbf{s}_0 = \mathbf{0}$ is a singular point, satisfying $\mathbf{g} = \mathbf{0}$, equation (S2.9) gives equation (14) in the main text.

### S3. Derivation of equation(15): Branching possibility conditions

#### S3.1. Main derivation

##### Settings

We consider an $L$-dimensional trait space, $\mathbf{s} = (x_1, ..., x_L)^T$, having an evolutionarily singular point denoted by $\mathbf{s}^* = (x_1^*, ..., x_L^*)^T$. We assume without loss of generality that $\mathbf{s}^* = \mathbf{0}$. Also, we assume without loss of generality that the trait space is locally normalized so that the mutational covariance matrix is given by $\mathbf{V}_\mu = \sigma^2 \mathbf{I}$ (with $\mathbf{I}$ describing the identity matrix) around $\mathbf{s}^*$, i.e., the standard deviation of mutational steps is given by $\sigma$ in all directions (see also Ito and Sasaki, 2020). In this trait space, we consider trait substitution sequences (provided that mutation rates are sufficiently low so that population dynamics is almost at equilibrium whenever a mutant emerges). We denote by $\mathfrak{B}(\mathbf{s}_0) = (\mathcal{S}^0, \mathcal{S}^1, ..., \mathcal{S}^K)$ a trait substitution sequence starting from a monomorphic resident $\mathbf{s}_0$ and being formed by sequential invasions by mutants $\mathbf{s}'^0, ..., \mathbf{s}'^{K-1}$, where $\mathcal{S}^0 = \{\mathbf{s}_0\}$ describes the initial monomorphic state, and where $\mathcal{S}^k = \{\mathbf{s}_1^k, ..., \mathbf{s}_M^k\}$ for $k = 1, 2, ..., K$ describes the state after the invasion by $\mathbf{s}'^{k-1}$, represented with the set of coexisting residents $\mathbf{s}_1^k, ..., \mathbf{s}_M^k$ (or a monomorphic resident).

##### Condition for monomorphic convergence

For describing monomorphic convergence, we define a set $\mathcal{M}^c$ of monomorphic states close to a singular point $\mathbf{s}^*$ such that these monomorphic residents are inside of a hypersphere with center $\mathbf{s}^* = \mathbf{0}$ and radius $c\sigma$:

$$\mathcal{M}^c \;:=\; \{\mathcal{S} \,|\, \mathcal{S} = \{\mathbf{s}\}, |\mathbf{s}| < c\sigma\}, \tag{S3.1}$$

where $c$ is a positive constant satisfying $1 \ll c \ll 1/\sigma$. Then, we denote by $P^{\mathrm{mc}}(\mathcal{S}^0)$ the probability for a trait substitution sequence starting from $\mathcal{S}_0$ to enter $\mathcal{M}^c$ (i.e., the probability for $\mathfrak{B}(\mathbf{s}_0)$ to have at least one state belonging to $\mathcal{M}^c$). By choosing a sufficiently large $|\mathbf{s}_0|$ for $\mathcal{S}^0 = \{\mathbf{s}_0\}$ such that $|\mathbf{s}_0| \gg c\sigma$ holds, we can approximately treat a trait substitution sequence starting from $\mathcal{S}^0 \notin \mathcal{M}^c$ as directional evolution described deterministically with the canonical equation. In this case, as derived in Section S2.2, we expect that

$$P^{\mathrm{mc}}_{\min} \;=\; \min_{\mathcal{S}^0 \notin \mathcal{M}^c} (P^{\mathrm{mc}}(\mathcal{S}^0)) = 1 - \varepsilon_{\mathrm{mc}} \tag{S3.2}$$

holds with $0 < \varepsilon_{\mathrm{mc}} \ll 1$ under a sufficiently large $c$ and $K$, provided that $\mathbf{s}^*$ is convergence stable, satisfying

$$\lambda_{\max}(\mathbf{C}^T) \;<\; 0 \tag{S3.3}$$

(Leimar, 2009).

##### Condition for dimorphic divergence

We assume that $\mathbf{s}^* = \mathbf{0}$ is evolutionarily unstable, satisfying

$$\lambda_{\max}(\mathbf{D}) \;>\; 0, \tag{S3.4}$$

in which case the eigenvector $\mathbf{e}$ corresponding to $\lambda_{\max}(\mathbf{D})$ gives the direction of the potential dimorphic divergence (Ito and Dieckmann, 2014; Geritz et al., 2016). For describing dimorphic divergence, we define a set $\mathcal{D}^c$ of dimorphic states each of which consists of two residents $\mathbf{s}_1$ and $\mathbf{s}_2$, where they are differentiated from each other along the direction of $\mathbf{e}$ by more than $2\,c\sigma$ with their differentiation in other directions being kept small (i.e., their difference $\Delta\,\mathbf{s} = \mathbf{s}_1 - \mathbf{s}_2$ satisfies $|\mathbf{e}^T\,\Delta\,\mathbf{s}| > 2\,c\sigma$ and $|\Delta\,\mathbf{s} - [\mathbf{e}^T\,\Delta\,\mathbf{s}]\,\mathbf{e}| < d\sigma$ with $d \ll c$), and where they are approximately symmetric about $\mathbf{s}^*$ (i.e., satisfies $|\frac{\mathbf{s}_1+\mathbf{s}_2}{2}| < d\sigma$):

$$\mathcal{D}^c \;:=\; \left\{ \mathcal{S} \;\middle|\; \begin{array}{l} \mathcal{S} = \{\mathbf{s}_1, \mathbf{s}_2\}, \\ |\mathbf{e}^T\,\Delta\,\mathbf{s}| > 2\,c\sigma, \\ |\Delta\,\mathbf{s} - [\mathbf{e}^T\,\Delta\,\mathbf{s}]\,\mathbf{e}| < d\sigma, \\ \left|\dfrac{\mathbf{s}_1+\mathbf{s}_2}{2}\right| < d\sigma \end{array} \right\} \tag{S3.5}$$

with $\Delta\,\mathbf{s} = \mathbf{s}_1 - \mathbf{s}_2$, provided that the coexistence condition for $\mathbf{s}_1 = \mathbf{e}|\Delta\,\mathbf{s}|/2$ and $\mathbf{s}_2 = -\mathbf{e}|\Delta\,\mathbf{s}|/2$,

$$\mathbf{e}^T[\mathbf{D} - \mathbf{C}]\,\mathbf{e} \;>\; 0, \tag{S3.6}$$

is satisfied.

By choosing a sufficiently large $c$ and $d$ so that $1 \ll d \ll c$ holds, we can deterministically describe trait substitution sequences starting from $\mathcal{S} \in \mathcal{D}^c$ with the canonical equations. As derived in Section S2.3, if

$$\lambda_{\max}\!\left(\mathbf{C} + \frac{\mathbf{C}\,\mathbf{e}\,\mathbf{e}^T\,\mathbf{C}}{\mathbf{e}^T[\mathbf{D}-\mathbf{C}]\,\mathbf{e}}\right) \;<\; \lambda_{\max}(\mathbf{D}) \tag{S3.7}$$

is satisfied, then the trajectories of the canonical equations give *locally stable* dimorphic divergence in the sense that $|\mathbf{s}_2 - \mathbf{s}_1|$ monotonically grows keeping the state inside of $\mathcal{D}^c$. In this case, we expect that a trait substitution sequence starting from a state $\mathcal{S} \in \mathcal{D}^c$ is likely to enter $\mathcal{D}^\infty$ defined by

$$\mathcal{D}^\infty \;:=\; \left\{ \mathcal{S} \;\middle|\; \begin{array}{l} \mathcal{S} = \{\mathbf{s}_1, \mathbf{s}_2\}, \\ |\mathbf{s}_2 - \mathbf{s}_1| = \infty \end{array} \right\}, \tag{S3.8}$$

so that the probability $P^{\mathrm{dd}}(\mathcal{S})$ for a trait substitution sequence starting from $\mathcal{S} \in \mathcal{D}^c$ to eventually enter $\mathcal{D}^\infty$ satisfies

$$P^{\mathrm{dd}}_{\min} \;=\; \min_{\mathcal{S} \in \mathcal{D}^c} (P^{\mathrm{dd}}(\mathcal{S})) = 1 - \varepsilon_{\mathrm{dd}} \tag{S3.9}$$

with $0 < \varepsilon_{\mathrm{dd}} \ll 1$ under a sufficiently large $c$ and $K$.

##### Transition from monomorphic convergence to dimorphic divergence

As derived in Section S2.4, if the conditions for convergence stability, equation (S3.3), and for locally stable dimorphic divergence, equations (S3.4, S3.6, S3.7), are satisfied, then there is a non-zero probability for a trait substitution sequence starting from an arbitrary state in $\mathcal{M}^c$ to enter $\mathcal{D}^c$, so that the following inequality holds,

$$P^{\mathrm{mc}\to\mathrm{dd}}_{\min} \;=\; \min_{\mathcal{S} \in \mathcal{M}^c} (P^{\to\mathrm{dd}}(\mathcal{S})) > 0. \tag{S3.10}$$

##### Branching possibility conditions

We denote by $P^{\mathrm{bran}}(\mathcal{S}^0)$ the probability for a trait substitution sequence starting from the initial state $\mathcal{S}^0$ to eventually enter $\mathcal{D}^\infty$. Among all possible trait substitution sequences that start from $\mathcal{S}_0$ and eventually enter $\mathcal{D}^\infty$, there exist some sequences that enter $\mathcal{M}^c, \mathcal{D}^c$, and $\mathcal{D}^\infty$ in this order. Hence, we see from equations (S3.2, S3.9, S3.10) that the following inequality holds,

$$P^{\mathrm{bran}}(\mathbf{s}_0) \;\geqslant\; P^{\mathrm{mc}}_{\min}\,P^{\mathrm{mc}\to\mathrm{dd}}_{\min}\,P^{\mathrm{dd}}_{\min} = [1-\varepsilon_{\mathrm{mc}}][1-\varepsilon_{\mathrm{dd}}]\,P^{\mathrm{mc}\to\mathrm{dd}}_{\min} > 0. \tag{S3.11}$$

Therefore, if $\mathbf{s}^*$ satisfies the conditions for convergence stability, the emergence of dimorphism, and their locally stable divergence, i.e., equations (S3.3, S3.4, S3.6, S3.7) called the *branching possibility conditions*, then there is a non-zero probability of evolutionary branching for a monomorphic resident of an arbitrary phenotype that is sufficiently close to $\mathbf{s}^*$ so that the approximate fitness function (equation (14) in the main text) is valid.

Because $\varepsilon_{\rm mc}$, $\varepsilon_{\rm dd}$, and $P_{\rm min}^{\rm mc\to dd}$ are independent of $\sigma$ as shown in Section S2.5, the branching possibility conditions ensure that equation (S3.11) holds even under an infinitesimally small $\sigma$. Finally, by expressing the branching possibility conditions given by equations (S3.3), (S3.4), (S3.6), and (S3.7) in the trait space before the local normalization where the mutational covariance matrix is given by $\mathbf{V}_\mu$), we obtain equation (15) in the main text (see Section S2.7 for the derivation).

#### S3.2. Derivation of equation (S3.3): Convergence stability

The canonical equation for directional evolution of a monomorphic state $\{\mathbf{s}\} \notin \mathcal{M}^c$ is given by equation (7) in the main text:

$$\frac{d\mathbf{s}}{\rm dt} = \frac{\mu\,\sigma^2\,\hat{n}}{2}\mathbf{V}_\mu\,\mathbf{C}^T\,\mathbf{s} = \frac{\mu\,\sigma^2\,\hat{n}}{2}\mathbf{C}^T\,\mathbf{s}. \tag{S3.12}$$

Hence, if $\lambda_{\max}(\mathbf{C}^T) < 0$ holds, then $\mathbf{s}^* = \mathbf{0}$ is locally stable (i.e., convergence stable), in which case equation (S3.2) holds under sufficiently large $c$ and $K$.

#### S3.3. Derivation of equation (S3.6) and (S3.7): Locally stable dimorphic divergence

##### Canonical equations for dimorphic divergence

We describe the directional coevolution of dimorphic residents $\mathbf{s}_1$ and $\mathbf{s}_2$ with the canonical equations:

$$\begin{aligned}\frac{d\mathbf{s}_1}{\rm dt} &= \frac{\mu\,\sigma^2[\hat{n}_1+\hat{n}_2]}{2}\,p_1\,\mathbf{g}(\mathbf{s}_1;\mathbf{s}_1,\mathbf{s}_2),\\ \frac{d\mathbf{s}_2}{\rm dt} &= \frac{\mu\,\sigma^2[\hat{n}_1+\hat{n}_2]}{2}\,p_2\,\mathbf{g}(\mathbf{s}_2;\mathbf{s}_1,\mathbf{s}_2),\end{aligned} \tag{S3.13}$$

where $\mathbf{g}(\mathbf{s}_1;\mathbf{s}_1,\mathbf{s}_2)$ and $\mathbf{g}(\mathbf{s}_2;\mathbf{s}_1,\mathbf{s}_2)$ are the fitness gradients for $\mathbf{s}_1$ and $\mathbf{s}_2$, respectively, and where $\hat{p}_1 = \hat{n}_1/[\hat{n}_1+\hat{n}_2]$ and $\hat{p}_2 = \hat{n}_2/[\hat{n}_1+\hat{n}_2]$ are their fractions at the population dynamical equilibrium. These fitness gradients and fractions are obtained as functions of $\mathbf{s}_1$, $\mathbf{s}_2$, $\mathbf{C}$, and $\mathbf{D}$, as follows. According to equation (S2.10), the invasion fitness function $f(\mathbf{s}';\mathbf{s}_1,\mathbf{s}_2)$ is approximately given by

$$f(\mathbf{s}';\mathbf{s}_1,\mathbf{s}_2) = \bar{\mathbf{s}}^T\,\mathbf{C}[\mathbf{s}'-\bar{\mathbf{s}}] + \frac{1}{2}[\mathbf{s}'-\bar{\mathbf{s}}]^T\,\mathbf{D}[\mathbf{s}'-\bar{\mathbf{s}}] - \frac{1}{2}V_{\mathbf{D}} \tag{S3.14}$$

with $\bar{\mathbf{s}} = \sum_{j=1}^2 \hat{p}_j\,\mathbf{s}_j$ and $V_{\mathbf{D}} = \sum_{j=1}^2 \hat{p}_j[\mathbf{s}_j-\bar{\mathbf{s}}]^T\,\mathbf{D}[\mathbf{s}_j-\bar{\mathbf{s}}]$. From this equation we get the fitness gradient $\mathbf{g}(\mathbf{s};\mathbf{s}_1,\mathbf{s}_2)$ at $\mathbf{s}$ as

$$\begin{aligned}\mathbf{g}(\mathbf{s};\mathbf{s}_1,\mathbf{s}_2) &= \left[\frac{f(\mathbf{s}';\mathbf{s}_1,\mathbf{s}_2)}{\partial\mathbf{s}'}\right]_{\mathbf{s}'=\mathbf{s}} = \mathbf{C}^T\,\bar{\mathbf{s}} + \mathbf{D}[\mathbf{s}-\bar{\mathbf{s}}],\\ \bar{\mathbf{s}} &= \hat{p}_1\,\mathbf{s}_1 + \hat{p}_2\,\mathbf{s}_2,\end{aligned} \tag{S3.15}$$

For convenience we introduce $\Delta\,\mathbf{s} = \mathbf{s}_1 - \mathbf{s}_2$ and $\breve{\mathbf{s}} = [\mathbf{s}_1+\mathbf{s}_2]/2$, by which $\mathbf{s}_1$ and $\mathbf{s}_2$ are expressed as

$$\mathbf{s}_1 = \breve{\mathbf{s}} + \frac{1}{2}\Delta\,\mathbf{s}, \quad \mathbf{s}_2 = \breve{\mathbf{s}} - \frac{1}{2}\Delta\,\mathbf{s}. \tag{S3.16}$$

We also introduce a scalar $B$ such that

$$\hat{p}_1 = \frac{1}{2} + B, \quad \hat{p}_2 = \frac{1}{2} - B, \tag{S3.17}$$

by which we can express $\bar{\mathbf{s}}$, $\mathbf{s}_1 - \bar{\mathbf{s}}$, and $\mathbf{s}_2 - \bar{\mathbf{s}}$ as

$$\begin{aligned} \bar{\mathbf{s}} &= \hat{p}_1 \mathbf{s}_1 + \hat{p}_2 \mathbf{s}_2 = \breve{\mathbf{s}} + B\,\Delta\,\mathbf{s}, \\ \mathbf{s}_1 - \bar{\mathbf{s}} &= \left[\frac{1}{2} - B\right] \Delta\,\mathbf{s}, \\ \mathbf{s}_2 - \bar{\mathbf{s}} &= \left[-\frac{1}{2} - B\right] \Delta\,\mathbf{s}. \end{aligned} \tag{S3.18}$$

Since $f(\mathbf{s}_1; \mathbf{s}_1, \mathbf{s}_2) = f(\mathbf{s}_2; \mathbf{s}_1, \mathbf{s}_2) = 0$ must hold as consistency conditions, we see from equations (S3.14), (S3.16), (S3.17), and (S3.18) that

$$\begin{aligned} f(\mathbf{s}_1; \mathbf{s}_1, \mathbf{s}_2) &= \left[\frac{1}{2} - B\right] [\breve{\mathbf{s}} + B\,\Delta\,\mathbf{s}]^T \mathbf{C}\,\Delta\,\mathbf{s} + \frac{1}{2}\left[\frac{1}{2} - B\right]^2 \Delta\,\mathbf{s}^T \mathbf{D}\,\Delta\,\mathbf{s} - \frac{1}{2} V_{\mathbf{D}} = 0, \\ f(\mathbf{s}_2; \mathbf{s}_1, \mathbf{s}_2) &= \left[-\frac{1}{2} - B\right] [\breve{\mathbf{s}} + B\,\Delta\,\mathbf{s}]^T \mathbf{C}\,\Delta\,\mathbf{s} + \frac{1}{2}\left[\frac{1}{2} + B\right]^2 \Delta\,\mathbf{s}^T \mathbf{D}\,\Delta\,\mathbf{s} - \frac{1}{2} V_{\mathbf{D}} = 0, \end{aligned} \tag{S3.19}$$

yielding

$$\begin{aligned} B &= \frac{\breve{\mathbf{s}}^T \mathbf{C}\,\Delta\,\mathbf{s}}{\Delta\,\mathbf{s}^T [\mathbf{D} - \mathbf{C}]\,\Delta\,\mathbf{s}}, \\ V_{\mathbf{D}} &= \left[\frac{1}{4} + B^2\right] \Delta\,\mathbf{s}^T \mathbf{D}\,\Delta\,\mathbf{s} - 2\,B[\breve{\mathbf{s}} + B\,\Delta\,\mathbf{s}]^T \mathbf{C}\,\Delta\,\mathbf{s}. \end{aligned} \tag{S3.20}$$

Note that the coexistence of $\mathbf{s}_1$ and $\mathbf{s}_2$ require their mutual invisibility:

$$\begin{aligned} f(\mathbf{s}_1; \mathbf{s}_2) &= \mathbf{s}_2^T \mathbf{C}[\mathbf{s}_1 - \mathbf{s}_2] + \frac{1}{2}[\mathbf{s}_1 - \mathbf{s}_2]^T \mathbf{D}[\mathbf{s}_1 - \mathbf{s}_2] > 0, \\ f(\mathbf{s}_2; \mathbf{s}_1) &= \mathbf{s}_1^T \mathbf{C}[\mathbf{s}_2 - \mathbf{s}_1] + \frac{1}{2}[\mathbf{s}_2 - \mathbf{s}_1]^T \mathbf{D}[\mathbf{s}_2 - \mathbf{s}_1] > 0, \end{aligned} \tag{S3.21}$$

which are combined into

$$\Delta\,\mathbf{s}^T [\mathbf{D} - \mathbf{C}]\,\Delta\,\mathbf{s} \;>\; 2|\breve{\mathbf{s}}^T \mathbf{C}\,\Delta\,\mathbf{s}|, \tag{S3.22}$$

and which is equivalent to $|B| < 1/2$ under $\Delta\,\mathbf{s}^T[\mathbf{D} - \mathbf{C}]\,\Delta\,\mathbf{s} > 0$.

Then by using equations (S3.15-S3.22), we transform equation (S3.13) into

$$\begin{aligned} \frac{d\mathbf{s}_1}{\mathrm{dt}} &= \frac{\mu\,\sigma^2[\hat{n}_1 + \hat{n}_2]}{2}\left[\frac{1}{2} + B\right] \{\mathbf{C}^T \bar{\mathbf{s}} + \mathbf{D}[\mathbf{s}_1 - \bar{\mathbf{s}}]\} \\ &= \frac{\mu\,\sigma^2[\hat{n}_1 + \hat{n}_2]}{2}\left[\frac{1}{2} + B\right] \left\{\mathbf{C}^T \breve{\mathbf{s}} + \frac{1}{2}\mathbf{D}\,\Delta\,\mathbf{s} + B[\mathbf{C}^T - \mathbf{D}]\,\Delta\,\mathbf{s}\right\}, \\ \frac{d\mathbf{s}_2}{\mathrm{dt}} &= \frac{\mu\,\sigma^2[\hat{n}_1 + \hat{n}_2]}{2}\left[\frac{1}{2} - B\right] \{\mathbf{C}^T \bar{\mathbf{s}} + \mathbf{D}[\mathbf{s}_2 - \bar{\mathbf{s}}]\} \\ &= \frac{\mu\,\sigma^2[\hat{n}_1 + \hat{n}_2]}{2}\left[\frac{1}{2} - B\right] \left\{\mathbf{C}^T \breve{\mathbf{s}} - \frac{1}{2}\mathbf{D}\,\Delta\,\mathbf{s} + B[\mathbf{C}^T - \mathbf{D}]\,\Delta\,\mathbf{s}\right\}, \end{aligned} \tag{S3.23}$$

and which gives

$$\begin{aligned} \frac{d\Delta\,\mathbf{s}}{\mathrm{dt}} &= \frac{d\mathbf{s}_1}{\mathrm{dt}} - \frac{d\mathbf{s}_2}{\mathrm{dt}} = \alpha\{\mathbf{D}\,\Delta\,\mathbf{s} + 4\,B\,\mathbf{C}^T \breve{\mathbf{s}} + 4\,B^2[\mathbf{C}^T - \mathbf{D}]\,\Delta\,\mathbf{s}\}, \\ \frac{d\,\breve{\mathbf{s}}}{\mathrm{dt}} &= \frac{1}{2}\left[\frac{d\mathbf{s}_1}{\mathrm{dt}} + \frac{d\mathbf{s}_2}{\mathrm{dt}}\right] = \alpha\,\mathbf{C}^T[\breve{\mathbf{s}} + \Delta\,\mathbf{s}\,B] \end{aligned} \tag{S3.24}$$

with $\alpha = \frac{\mu\,\sigma^2[\hat{n}_1 + \hat{n}_2]}{4}$.

**Local stability for dimorphic divergence**

Here we derive a sufficient condition for the locally stable dimorphic divergence. For convenience we decompose $\mathbf{D}$ as

$$\begin{aligned}\mathbf{D} &= \mathbf{E}\begin{pmatrix} D_1 & \cdots & 0 \\ \vdots & \ddots & \vdots \\ 0 & \cdots & D_L \end{pmatrix}\mathbf{E}^T, \\ \mathbf{E} &= (\ \mathbf{e}_1 \ \cdots \ \mathbf{e}_L \ ),\end{aligned} \tag{S3.25}$$

satisfying $\lambda_{\max}(\mathbf{D}) = D_1 > 0$, and $D_1 > D_j$ for all $j = 2, ..., L$. By introducing $\Delta z = \mathbf{e}^T \Delta \mathbf{s}$ with $\mathbf{e} = \mathbf{e}_1$, we transform equation (S3.24) into the dynamics of $\Delta z$ and $\left(\frac{\Delta \mathbf{s}}{\Delta z}, \frac{\breve{\mathbf{s}}}{\Delta z}\right)$ as

$$\begin{aligned}\frac{d\Delta z}{\mathrm{dt}} &= \mathbf{e}^T \frac{d\Delta \mathbf{s}}{\mathrm{dt}} \\ &= \alpha\, \mathbf{e}^T\{\mathbf{D}\,\Delta \mathbf{s} + 4\,B\,\mathbf{C}^T\,\breve{\mathbf{s}} + 4\,B^2[\mathbf{C}^T - \mathbf{D}]\,\Delta \mathbf{s}\} \\ &= \alpha\{\mathbf{e}^T\,\mathbf{D}\,\Delta \mathbf{s} + 4\,B\,\mathbf{e}^T\,\mathbf{C}^T\,\breve{\mathbf{s}} + 4\,B^2\,\mathbf{e}^T[\mathbf{C}^T - \mathbf{D}]\,\Delta \mathbf{s}\} \\ &= \alpha\left\{\mathbf{e}^T\,\mathbf{D}\,\frac{\Delta \mathbf{s}}{\Delta z} + 4\,B\,\mathbf{e}^T\,\mathbf{C}^T\,\frac{\breve{\mathbf{s}}}{\Delta z} + 4\,B^2\,\mathbf{e}^T[\mathbf{C}^T - \mathbf{D}]\,\frac{\Delta \mathbf{s}}{\Delta z}\right\}\Delta z,\end{aligned}$$

$$\begin{aligned}\frac{d}{\mathrm{dt}}\left[\frac{\Delta \mathbf{s}}{\Delta z}\right] &= \frac{1}{\Delta z}\frac{d\Delta \mathbf{s}}{\mathrm{dt}} - \frac{\Delta \mathbf{s}}{\Delta z^2}\frac{d\Delta z}{\mathrm{dt}} \\ &= \frac{1}{\Delta z}\alpha\{\mathbf{D}\,\Delta \mathbf{s} + 4\,B\,\mathbf{C}^T\,\breve{\mathbf{s}} + 4\,B^2[\mathbf{C}^T - \mathbf{D}]\,\Delta \mathbf{s}\} \\ &\quad - \frac{\Delta \mathbf{s}}{\Delta z^2}\alpha\left\{\mathbf{e}^T\,\mathbf{D}\,\frac{\Delta \mathbf{s}}{\Delta z} + 4\,B\,\mathbf{e}^T\,\mathbf{C}^T\,\frac{\breve{\mathbf{s}}}{\Delta z} + 4\,B^2\,\mathbf{e}^T[\mathbf{C}^T - \mathbf{D}]\,\frac{\Delta \mathbf{s}}{\Delta z}\right\}\Delta z \\ &= \alpha\left\{\mathbf{D} - \mathbf{e}^T\,\mathbf{D}\,\frac{\Delta \mathbf{s}}{\Delta z}\right\}\frac{\Delta \mathbf{s}}{\Delta z} \\ &\quad + 4\,\alpha\,B\left\{\mathbf{C}^T\,\frac{\breve{\mathbf{s}}}{\Delta z} + B[\mathbf{C}^T - \mathbf{D}]\,\frac{\Delta \mathbf{s}}{\Delta z} - \mathbf{e}^T\,\mathbf{C}^T\,\frac{\breve{\mathbf{s}}}{\Delta z}\,\frac{\Delta \mathbf{s}}{\Delta z} - B\,\mathbf{e}^T[\mathbf{C}^T - \mathbf{D}]\,\frac{\Delta \mathbf{s}}{\Delta z}\,\frac{\Delta \mathbf{s}}{\Delta z}\right\},\end{aligned}$$

$$\begin{aligned}\frac{d}{\mathrm{dt}}\left[\frac{\breve{\mathbf{s}}}{\Delta z}\right] &= \frac{1}{\Delta z}\frac{d\breve{\mathbf{s}}}{\mathrm{dt}} - \frac{\breve{\mathbf{s}}}{\Delta z^2}\frac{d\Delta z}{\mathrm{dt}} \\ &= \frac{1}{\Delta z}\{\alpha\ \mathbf{C}^T[\breve{\mathbf{s}} + \Delta\ \mathbf{s}\ B]\} - \frac{\breve{\mathbf{s}}}{\Delta z^2}\ \alpha\left\{\mathbf{e}^T\ \mathbf{D}\ \frac{\Delta \mathbf{s}}{\Delta z} + 4\ B\ \mathbf{e}^T\ \mathbf{C}^T\ \frac{\breve{\mathbf{s}}}{\Delta z} + 4\ B^2\ \mathbf{e}^T[\mathbf{C}^T - \right. \\ &\quad \left.\mathbf{D}]\,\frac{\Delta \mathbf{s}}{\Delta z}\right\}\Delta z \\ &= \frac{1}{\Delta z}\alpha\left\{\mathbf{C}^T + \frac{\mathbf{C}^T\,\Delta \mathbf{s}\,\Delta \mathbf{s}^T\,\mathbf{C}^T}{\Delta \mathbf{s}^T[\mathbf{D} - \mathbf{C}]\,\Delta \mathbf{s}}\right\}\breve{\mathbf{s}} \\ &\quad - \frac{\breve{\mathbf{s}}}{\Delta z^2}\alpha\left\{\mathbf{e}^T\,\mathbf{D}\,\frac{\Delta \mathbf{s}}{\Delta z} + 4\,B\,\mathbf{e}^T\,\mathbf{C}^T\,\frac{\breve{\mathbf{s}}}{\Delta z} + 4\,B^2\,\mathbf{e}^T[\mathbf{C}^T - \mathbf{D}]\,\frac{\Delta \mathbf{s}}{\Delta z}\right\}\Delta z \\ &= \alpha\left[\mathbf{C}^T + \frac{\mathbf{C}^T\,\Delta \mathbf{s}\,\Delta \mathbf{s}^T\,\mathbf{C}^T}{\Delta \mathbf{s}^T[\mathbf{D} - \mathbf{C}]\,\Delta \mathbf{s}} - \mathbf{e}^T\ \mathbf{D}\ \frac{\Delta \mathbf{s}}{\Delta z} - 4\ B\ \mathbf{e}^T\ \mathbf{C}^T\ \frac{\breve{\mathbf{s}}}{\Delta z} - 4\ B^2\ \mathbf{e}^T[\mathbf{C}^T - \right. \\ &\quad \left.\mathbf{D}]\,\frac{\Delta \mathbf{s}}{\Delta z}\right]\frac{\breve{\mathbf{s}}}{\Delta z}.\end{aligned} \tag{S3.26}$$

We see that $\left(\frac{\Delta \mathbf{s}}{\Delta z}, \frac{\breve{\mathbf{s}}}{\Delta z}\right)$ has a fixed point at $\left(\frac{\Delta \mathbf{s}}{\Delta z}, \frac{\breve{\mathbf{s}}}{\Delta z}\right) = (\mathbf{e}, \mathbf{0})$ because $\frac{\breve{\mathbf{s}}}{\Delta z} = \mathbf{0}$ gives $B = 0$ at equation (S3.20). To analyze its local stability, we introduce

$$\begin{aligned}\mathbf{u} &= \frac{\Delta \mathbf{s}}{\Delta z} - \mathbf{e}, \\ \mathbf{w} &= \frac{\breve{\mathbf{s}}}{\Delta z},\end{aligned} \tag{S3.27}$$

and assume that $|\mathbf{u}| \ll 1$ and $|\mathbf{w}| \ll 1$. Substitution of equation (S3.27) into equation (S3.20) gives

$$B = \frac{\check{\mathbf{s}}^T\,\mathbf{C}\,\Delta\,\mathbf{s}}{\Delta\,\mathbf{s}^T[\mathbf{D}-\mathbf{C}]\,\Delta\,\mathbf{s}} = \frac{[\Delta\,z\,\mathbf{w}]^T\,\mathbf{C}\,\Delta\,z[\mathbf{u}+\mathbf{e}]}{\Delta\,z[\mathbf{u}+\mathbf{e}]^T[\mathbf{D}-\mathbf{C}]\,\Delta\,z[\mathbf{u}+\mathbf{e}]} = \frac{\mathbf{w}^T\,\mathbf{C}[\mathbf{u}+\mathbf{e}]}{[\mathbf{u}+\mathbf{e}]^T[\mathbf{D}-\mathbf{C}][\mathbf{u}+\mathbf{e}]} = O(|\mathbf{w}|). \tag{S3.28}$$

From equations (S3.26-S3.28) we obtain

$$\begin{aligned}\frac{d\Delta\,z}{\mathrm{dt}} &= \alpha\{\mathbf{e}^T\,\mathbf{D}[\mathbf{u}+\mathbf{e}]+4\,B\,\mathbf{e}^T\,\mathbf{C}^T\,\mathbf{w}+4\,B^2\,\mathbf{e}^T[\mathbf{C}^T-\mathbf{D}][\mathbf{u}+\mathbf{e}]\}\,\Delta\,z\\ &= \alpha[D_1+O(|\mathbf{u}|)+O(|\mathbf{w}|)]\,\Delta\,z\end{aligned} \tag{S3.29}$$

and

$$\begin{aligned}\frac{d\mathbf{u}}{\mathrm{dt}} &= \frac{d}{\mathrm{dt}}\left[\frac{\Delta\,\mathbf{s}}{\Delta\,z}\right]\\
&= \alpha\{\mathbf{D}-\mathbf{e}^T\,\mathbf{D}[\mathbf{u}+\mathbf{e}]\}[\mathbf{u}+\mathbf{e}]\\
&\quad +4\,\alpha\,B\{\mathbf{C}^T\,\mathbf{w}+B[\mathbf{C}^T-\mathbf{D}][\mathbf{u}+\mathbf{e}]-\mathbf{e}^T\,\mathbf{C}^T\,\mathbf{w}[\mathbf{u}+\mathbf{e}]-B\,\mathbf{e}^T[\mathbf{C}^T-\mathbf{D}][\mathbf{u}+\mathbf{e}][\mathbf{u}+\mathbf{e}]\}\\
&= \alpha\{\mathbf{D}[\mathbf{u}+\mathbf{e}]-\mathbf{e}^T\,\mathbf{D}[\mathbf{u}+\mathbf{e}][\mathbf{u}+\mathbf{e}]\}+O(|\mathbf{w}|^2)\\
&= \alpha\{\mathbf{D}\,\mathbf{u}+D_1\,\mathbf{e}-[D_1\,\mathbf{e}^T\,\mathbf{u}+D_1][\mathbf{u}+\mathbf{e}]\}+O(|\mathbf{w}|^2)\\
&= \alpha\{\mathbf{D}\,\mathbf{u}+D_1\,\mathbf{e}-D_1\,\mathbf{e}^T\,\mathbf{u}\,\mathbf{u}-D_1\,\mathbf{e}^T\,\mathbf{u}\,\mathbf{e}-D_1\,\mathbf{u}-D_1\,\mathbf{e}\}+O(|\mathbf{w}|^2)\\
&= \alpha\{\mathbf{D}\,\mathbf{u}-D_1\,\mathbf{e}^T\,\mathbf{u}\,\mathbf{u}-D_1\,\mathbf{e}\,\mathbf{e}^T\,\mathbf{u}-D_1\,\mathbf{u}\}+O(|\mathbf{w}|^2)\\
&= \alpha\{\mathbf{D}-D_1\,\mathbf{e}\,\mathbf{e}^T-D_1\}\,\mathbf{u}+O(|\mathbf{u}|^2)+O(|\mathbf{w}|^2)\\
&= \alpha\left\{(\,\mathbf{e}_1\;\cdots\;\mathbf{e}_L\,)\begin{pmatrix}D_1&\cdots&0\\ \vdots&\ddots&\vdots\\ 0&\cdots&D_L\end{pmatrix}\begin{pmatrix}\mathbf{e}_1^T\\ \vdots\\ \mathbf{e}_L^T\end{pmatrix}-D_1\,\mathbf{e}_1\,\mathbf{e}_1^T-D_1\,\mathbf{I}\right\}\mathbf{u}+O(|\mathbf{u}|^2)+O(|\mathbf{w}|^2)\\
&= \alpha\left\{\sum_{i=1}^{L}D_i\,\mathbf{e}_i\,\mathbf{e}_i^T-D_1\,\mathbf{e}_1\,\mathbf{e}_1^T-\sum_{i=1}^{L}D_1\,\mathbf{e}_i\,\mathbf{e}_i^T\right\}\mathbf{u}+O(|\mathbf{u}|^2)+O(|\mathbf{w}|^2)\\
&= \alpha\left\{(\,\mathbf{e}_1\;\cdots\;\mathbf{e}_L\,)\begin{pmatrix}-D_1&0&\cdots&0\\ 0&D_2-D_1&\cdots&0\\ \vdots&\vdots&\ddots&\vdots\\ 0&0&\cdots&D_L-D_1\end{pmatrix}\begin{pmatrix}\mathbf{e}_1^T\\ \vdots\\ \mathbf{e}_L^T\end{pmatrix}\right\}\mathbf{u}+O(|\mathbf{u}|^2)+O(|\mathbf{w}|^2)\\
&= \alpha\,\tilde{\mathbf{D}}\,\mathbf{u}+O(|\mathbf{u}|^2)+O(|\mathbf{w}|^2),\\
\tilde{\mathbf{D}} &= (\,\mathbf{e}_1\;\cdots\;\mathbf{e}_L\,)\begin{pmatrix}-D_1&0&\cdots&0\\ 0&D_2-D_1&\cdots&0\\ \vdots&\vdots&\ddots&\vdots\\ 0&0&\cdots&D_L-D_1\end{pmatrix}\begin{pmatrix}\mathbf{e}_1^T\\ \vdots\\ \mathbf{e}_L^T\end{pmatrix}\end{aligned} \tag{S3.30}$$

and

$$\begin{aligned}\frac{d\mathbf{w}}{\mathrm{dt}} &= \alpha\left[\mathbf{C}^T+\frac{\mathbf{C}^T[\mathbf{u}+\mathbf{e}][\mathbf{u}+\mathbf{e}]^T\,\mathbf{C}^T}{[\mathbf{u}+\mathbf{e}]^T[\mathbf{u}+\mathbf{e}]}-\mathbf{e}^T\,\mathbf{D}[\mathbf{u}+\mathbf{e}]-4\,B\,\mathbf{e}^T\,\mathbf{C}^T\,\mathbf{w}-4\,B^2\,\mathbf{e}^T[\mathbf{C}^T-\mathbf{D}]\,\mathbf{u}\right]\mathbf{w}\\
&= \alpha\left[\mathbf{C}^T+\frac{\mathbf{C}\,\mathbf{e}\,\mathbf{e}^T\,\mathbf{C}}{\mathbf{e}^T[\mathbf{D}-\mathbf{C}]\,\mathbf{e}}-\mathbf{e}^T\,\mathbf{D}\,\mathbf{e}+O(|\mathbf{u}|)+O(|\mathbf{w}|^2)\right]\mathbf{w}\\
&= \alpha[\mathbf{C}_{\mathrm{d}}^T-D_1\,\mathbf{I}]\,\mathbf{w}+O(|\mathbf{u}||\mathbf{w}|)+O(|\mathbf{w}|^3)\end{aligned} \tag{S3.31}$$

with

$$\mathbf{C}_{\mathrm{d}} = \mathbf{C}+\frac{\mathbf{C}\,\mathbf{e}\,\mathbf{e}^T\,\mathbf{C}}{\mathbf{e}^T[\mathbf{D}-\mathbf{C}]\,\mathbf{e}}. \tag{S3.32}$$

Hence, $\left(\frac{\Delta\,\mathbf{s}}{\Delta\,z},\frac{\check{\mathbf{s}}}{\Delta\,z}\right)=(\mathbf{e},\mathbf{0})$ is locally stable if

$$\begin{aligned}\lambda_{\max}(\mathbf{C}_{\mathrm{d}}^T-D_1\,\mathbf{I}) &< 0,\\ \lambda_{\max}(\tilde{\mathbf{D}}) &< 0,\end{aligned} \tag{S3.33}$$

both hold. Clearly, the second inequality is always satisfied because $\lambda_{\max}(\tilde{\mathbf{D}}) = \max\{-D_1, D_2 - D_1, ..., D_L - D_1\} < 0$ holds under the assumption of $D_1 = \lambda_{\max}(\mathbf{D}) > D_j$ for all $j = 2, ..., L$. Regarding the first inequality, because $\lambda_{\max}(\mathbf{C}_{\rm d})$ with its corresponding eigenvector $\mathbf{v}_{\max}$ satisfies $[\mathbf{C}_{\rm d} - D_1\,\mathbf{I}]\,\mathbf{v}_{\max} = [\lambda_{\max}(\mathbf{C}_{\rm d}) - D_1]\,\mathbf{v}_{\max}$, we can transform $\lambda_{\max}(\mathbf{C}_{\rm d} - D_{11}\,\mathbf{I})$ as

$$\lambda_{\max}(\mathbf{C}_{\rm d}^T - D_1\,\mathbf{I}) = \lambda_{\max}(\mathbf{C}_{\rm d}^T) - D_1 \;=\; \lambda_{\max}(\mathbf{C}_{\rm d}^T) - \lambda_{\max}(\mathbf{D}). \tag{S3.34}$$

Substituting equation (S3.35) into the first inequality of equation (S3.34), we obtain a sufficient condition for $\left(\frac{\Delta\,\mathbf{s}}{\Delta\,z}, \frac{\breve{\mathbf{s}}}{\Delta\,z}\right) = (\mathbf{e}, \mathbf{0})$ being locally stable:

$$\lambda_{\max}(\mathbf{C}_{\rm d}^T) \;<\; \lambda_{\max}(\mathbf{D}) \tag{S3.35}$$

with equation (S3.32), provided that the dimorphic residents are kept mutually invasible, i.e., equation (S3.21) with $\Delta\,\mathbf{s} = \Delta\,z\,\mathbf{e}$ and $\breve{\mathbf{s}} = \mathbf{0}$:

$$\mathbf{e}^T[\mathbf{D} - \mathbf{C}]\,\mathbf{e} \;>\; 0. \tag{S3.36}$$

Note that $\{\mathbf{s}_1, \mathbf{s}_2\}$ belonging to $\mathcal{D}^c$ with $1 \ll d \ll c$ satisfies $|\mathbf{u}| \ll 1$ and $|\mathbf{w}| \ll 1$, because we see from equations (S3.5) and (S3.16) that

$$\begin{aligned} |\Delta\,z| &= |\mathbf{e}^T\,\Delta\,\mathbf{s}| > 2\,c\sigma, \\ |\mathbf{u}| &= \left|\frac{\Delta\,\mathbf{s}}{\Delta\,z} - \mathbf{e}\right| = \frac{|\Delta\,\mathbf{s} - [\mathbf{e}^T\,\Delta\,\mathbf{s}]\,\mathbf{e}|}{|\Delta\,z|} < \frac{d\sigma}{2\,c\sigma} = \frac{d}{2\,c}, \\ |\mathbf{w}| &= \left|\frac{\breve{\mathbf{s}}}{\Delta\,z}\right| = \frac{|\mathbf{s}_1 + \mathbf{s}_2|}{2|\Delta\,z|} < \frac{d}{2\,c}. \end{aligned} \tag{S3.37}$$

Therefore, equations (S3.35) and (S3.36) are sufficient for equation (S3.9) under a sufficiently large $c$ and $K$. Note that equation (S3.24) can be transformed under $|\mathbf{u}| \ll 1$ and $|\mathbf{w}| \ll 1$ as

$$\begin{aligned} \frac{d\Delta\,\mathbf{s}}{\mathrm{dt}} &= \alpha\{\mathbf{D} + O(|\mathbf{w}|)\}\,\Delta\,\mathbf{s}, \\ \frac{d\,\breve{\mathbf{s}}}{\mathrm{dt}} &= \alpha\left\{\mathbf{C}^T + \frac{\mathbf{C}^T\,\Delta\,\mathbf{s}\,\Delta\,\mathbf{s}^T\,\mathbf{C}^T}{\Delta\,\mathbf{s}^T[\mathbf{D} - \mathbf{C}]\,\Delta\,\mathbf{s}}\right\}\breve{\mathbf{s}} \\ &= \alpha\{\mathbf{C}_{\rm d}^T + O(|\mathbf{u}|)\}\,\breve{\mathbf{s}}. \end{aligned} \tag{S3.38}$$

Hence, equation (S3.35) implies that the growth of $\Delta\,\mathbf{s}$ (i.e., the speed of evolutionary divergence) is faster than that of $\breve{\mathbf{s}}$ (i.e., the speed of deviation of the intermediate phenotype $\breve{\mathbf{s}}$ from the singular point).

#### S3.4. Derivation of equation (S3.10): Transition possibility from monomorphic convergence to dimorphic divergence

##### S3.4.1 Main derivation

We assume that $\mathbf{s}^* = \mathbf{0}$ satisfies the conditions for monomorphic convergence and for dimorphic divergence, equations (S3.3), (S3.4), (S3.6), and (S3.7). For a mutant $\mathbf{s}'$ under a monomorphic resident $\mathbf{s} = (x_1, ..., x_L)^T$ with $|\mathbf{s}' - \mathbf{s}^*|$ and $|\mathbf{s} - \mathbf{s}^*|$ being sufficiently small, the invasion fitness of $\mathbf{s}'$ is approximately given by equation (S1.4) with $\mathbf{g} = \mathbf{0}$:

$$f(\mathbf{s}'; \mathbf{s}) \;=\; \mathbf{s}^T\,\mathbf{C}[\mathbf{s}' - \mathbf{s}] + \frac{1}{2}[\mathbf{s}' - \mathbf{s}]^T\,\mathbf{D}[\mathbf{s}' - \mathbf{s}], \tag{S3.39}$$

from which the fitness gradient at $\mathbf{s}$ is derived as

$$\mathbf{g}(\mathbf{s}) \;=\; \begin{pmatrix} \frac{\partial f(\mathbf{s}';\mathbf{s})}{\partial x_1'} \\ \vdots \\ \frac{\partial f(\mathbf{s}';\mathbf{s})}{\partial x_L'} \end{pmatrix}_{\mathbf{s}'=\mathbf{s}} = \mathbf{C}^T\,\mathbf{s}. \tag{S3.40}$$

We consider an arbitrary monomorphic state $S^{\hat{K}} = \{\mathbf{s}^{\hat{K}}\} \in \mathcal{M}^c$ formed after $\hat{K}$ invasions in a trait substitution sequence $\mathfrak{B}(\mathbf{s}_0) = (S^0, S^1, ..., S^{\hat{K}}, ..., S^K)$ starting from a monomorphic resident $\mathbf{s}_0$ and formed by invading mutants $\mathbf{s}'^0, \mathbf{s}'^1, ..., \mathbf{s}'^{\hat{K}-1}, ..., \mathbf{s}'^{K-1}$. For the subsequent invading mutants, denoted by $\mathbf{s}'^k$ for $k = \hat{K}, \hat{K}+1, ..., \phi - 1$, we choose each invading mutant $\mathbf{s}'^k = \mathbf{s}'^k_g$ with

$$\mathbf{s}'^k_g \;=\; \mathbf{s}^k + \frac{\mathbf{g}(\mathbf{s}^k)}{|\mathbf{g}(\mathbf{s}^k)|}\,\rho_k\,\sigma \tag{S3.41}$$

with $0 < \rho_k \leqslant 1$. The invading mutant $\mathbf{s}'^k_g$ replaces the resident $\mathbf{s}^k$, as long as $\mathbf{s}'^k_g$ and $\mathbf{s}^k$ satisfy

$$\begin{aligned} f(\mathbf{s}'^k_g; \mathbf{s}^k) \;&=\; \rho_k\,\sigma\left[|\mathbf{g}(\mathbf{s}^k)| + \frac{1}{2}\,\frac{\mathbf{g}(\mathbf{s}^k)^T\,\mathbf{D}\,\mathbf{g}(\mathbf{s}^k)}{|\mathbf{g}(\mathbf{s}^k)|^2}\,\rho_k\,\sigma\right] > 0, \\ f(\mathbf{s}^k; \mathbf{s}'^k_g) \;&=\; \rho_k\,\sigma\left[-|\mathbf{g}(\mathbf{s}^k)| + \frac{1}{2}\,\frac{\mathbf{g}(\mathbf{s}^k)^T[\mathbf{D} - 2\,\mathbf{C}]\,\mathbf{g}(\mathbf{s}^k)}{|\mathbf{g}(\mathbf{s}^k)|^2}\,\rho_k\,\sigma\right] < 0. \end{aligned} \tag{S3.42}$$

Hence, by choosing sufficiently small $\rho_k$ for each invasion so that each invading mutant always replaces the resident, and by choosing a sufficiently large $\phi$ (i.e., large number of the subsequent invasions), we can make $|\mathbf{s}^\phi|$ satisfy

$$|\mathbf{s}^\phi| \;<\; \sigma\,\frac{\mathbf{e}^T[\mathbf{D} - \mathbf{C}]\,\mathbf{e}}{4\|\mathbf{C}\|} \tag{S3.43}$$

with $\mathbf{e}$ describing the eigenvector for $\lambda_{\max}(\mathbf{D})$ and $\|\mathbf{C}\| = \max_{\tilde{\mathbf{s}}}\ (\|\mathbf{C}\tilde{\mathbf{s}}\|\ /\ \|\tilde{\mathbf{s}}\|)$ with $\|\tilde{\mathbf{s}}\| = \sqrt{\tilde{s}_1^2 + \cdots + \tilde{s}_L^2}$.

Then, for the subsequent invading mutants after $\mathbf{s}'^k$ with $k = \phi - 1$, we assume that mutation occurs only in the direction of $\mathbf{e}$ (e.g., mutant $\mathbf{s}'$ deriving from resident $\mathbf{s}$ is described as $\mathbf{s}' = \mathbf{s} + \mathbf{e}\,\delta\,z'$ with a scalar $\delta\,z'$). Specifically, we choose each $\mathbf{s}'^k$ for $k = \phi, ..., K-1$ as a mutant deriving from the $q_k$th resident $\mathbf{s}^k_{q_k}$ in $S^k$, expressed as

$$\mathbf{s}'^k \;=\; \mathbf{s}^k_{q_k} + \mathbf{e}\,\delta\,z'^k. \tag{S3.44}$$

Consequently, in the subsequent trait substitution sequences formed by equation (S3.44), residents and invading mutants are all on the line $\mathbf{s} = \mathbf{s}_\phi + \mathbf{e}\,z$ with $z$ being a scalar parameter. Hence, the trait space can be treated as a one-dimensional space described with $z$. Accordingly, we convert equation (S3.44) into

$$z'^k \;=\; \mathbf{e}^T[\mathbf{s}'^k - \mathbf{s}_\phi] = \mathbf{e}^T[\mathbf{s}^k_{q_k} - \mathbf{s}_\phi] + \delta\,z'^k = z^k_{q_k} + \delta\,z'^k. \tag{S3.45}$$

As derived in Appendix S3.4.2, the invasion fitness for a mutant $z'$ under a monomorphic resident $z$ is given by,

$$\tilde{f}(z'; z) \;=\; \tilde{g}[z' - z] + \tilde{C}\,z[z' - z] + \frac{1}{2}\,\tilde{D}[z' - z]^2 \tag{S3.46}$$

with $\tilde{g} = \mathbf{s}^{\phi T}\,\mathbf{C}\,\mathbf{e}$, $\tilde{C} = \mathbf{e}^T\,\mathbf{C}\,\mathbf{e}$, $\tilde{D} = \mathbf{e}^T\,\mathbf{D}\,\mathbf{e}$. We assume without loss of generality that the sign of $\mathbf{e}$ is chosen so that $\tilde{g} = \mathbf{s}^{\phi T}\,\mathbf{C}\,\mathbf{e} \geqslant 0$ holds. Also, as derived in Section S2.4.2, the invasion fitness function for a mutant $z'$ under dimorphic residents $z_1$ and $z_2$ is given by

$$\tilde{f}(z'; z_1, z_2) \;=\; \frac{\tilde{D}}{2}[z' - z_1][z' - z_2] \tag{S3.47}$$

(see also Geritz et al. (1997)). Note that the coexistence of more than two resident phenotypes at population dynamical equilibrium is impossible, because the fitness function has a constant parabolic shape determined by $\tilde{D}$ (see Appendix S3.4.2).

On this basis, we choose $z'^{\phi}$ and $z'^{\phi+1}$ as

$$\begin{aligned} z'^{\phi} &= z^{\phi} + \sigma/2 = \sigma/2 \\ z'^{\phi+1} &= z'^{\phi} - \sigma = z^{\phi} - \sigma/2 = -\sigma/2, \end{aligned} \tag{S3.48}$$

where $z^{\phi} = \mathbf{e}^T[\mathbf{s}_{\phi} - \mathbf{s}_{\phi}] = 0$. As shown in Appendix S3.4.3, the residents after the invasion by $z'^{\phi+1}$ are dimorphic, given by $z_1^{\phi+2} = z'^{\phi} = \sigma/2$ and $z_2^{\phi+2} = z'^{\phi+1} = -\sigma/2$.

Next, we consider the subsequent invading mutants after $z'^{\phi+1}$, denoted by $z'^k$ for $k = \phi+2, ..., K-1$. As long as the residents are dimorphic, denoted by $z_1^k$ and $z_2^k$ with $z_2^k < z_1^k$, we choose each $z'^k$ as

$$\begin{aligned} z'^k &= \begin{cases} z_1^k + \rho_k \sigma & \text{for } |z_2^k| > |z_1^k| \\ z_2^k - \rho_k \sigma & \text{for otherwise} \end{cases}, \\ \rho_k &= \begin{cases} 1 & \text{for } \frac{|\tilde{C}|}{[\tilde{D} - \tilde{C}]} \leqslant 1 \\ \min\left\{1, \frac{\Delta z^k}{2\sigma}\left[\frac{|\tilde{C}|}{[\tilde{D} - \tilde{C}]} - \frac{1}{2}\right]^{-1}\right\} & \text{otherwise,} \end{cases} \end{aligned} \tag{S3.49}$$

where $\rho_k$ clearly satisfies $0 < \rho_k \leqslant 1$. As derived in Section S2.4, each invading mutant $z'^k$ given by equation (S3.49) (for $k = \phi+2$, ..., $K-1$) excludes its parental resident among $z_1^k$ and $z_2^k$ and coexists with the other resident (i.e., $z'^k = z_1^k + \rho_k \sigma$ gives $z_1^{k+1} = z'^k$ and $z_2^{k+1} = z_2^k$, whereas $z'^k = z_2^k - \rho_k \sigma$ gives $z_1^{k+1} = z_1^k$ and $z_2^{k+1} = z'^k$). Hence, the following relationships hold for all $k = \phi+2, ..., K-1$:

$$\begin{aligned} \Delta z^k &= \Delta z^{k-1} + \rho_{k-1}\sigma, \\ \breve{z}^k &= \begin{cases} \frac{z_1^{k-1} + z_2^{k-1} + \rho_{k-1}\sigma}{2} = \breve{z}^{k-1} + \frac{\sigma \rho_{k-1}}{2} & \text{for } \breve{z}^{k-1} < 0 \\ \frac{z_1^{k-1} + z_2^{k-1} - \rho_{k-1}\sigma}{2} = \breve{z}^{k-1} - \frac{\sigma \rho_{k-1}}{2} & \text{for } \breve{z}^{k-1} > 0, \end{cases} \\ 2|\breve{z}^k| &\leqslant \max\{2|\breve{z}^{k-1}|, \rho_{k-1}\sigma\}. \end{aligned} \tag{S3.50}$$

Consequently, equations (S3.49) and (S3.50) give a monotonic increase of $\Delta z^k$ through the repeated invasions, satisfying

$$\begin{aligned} \Delta z^k &= \Delta z^{k-1} + \rho_{k-1}\sigma, \\ &= \Delta z^{k-2} + \rho_{k-2}\sigma + \rho_{k-1}\sigma = \cdots = \Delta z^{\phi+2} + \rho_{\phi+2}\sigma + \ldots + \rho_{k-2}\sigma + \rho_{k-1}\sigma \\ &= \left[1 + \sum_{j=\phi+2}^{k-1} \rho_j\right]\sigma, \\ 2|\breve{z}^k| &\leqslant \max\{2|\breve{z}^{k-1}|, \rho_{k-1}\sigma\} \\ &\leqslant \max\{2|\breve{z}^{k-2}|, \rho_{k-2}\sigma, \rho_{k-1}\sigma\} \leqslant \cdots \leqslant <\max\{2|\breve{z}^{\phi+2}|, \rho_{\phi+2}\sigma, ..., \rho_{k-1}\sigma\} \\ &= \max\{0, \rho_{\phi+2}\sigma, ..., \rho_{k-1}\sigma\} \leqslant \sigma. \end{aligned} \tag{S3.51}$$

Equations (S3.51) and (S3.43) give

$$\begin{aligned} |\mathbf{e}^T \Delta \mathbf{s}^k| &= \Delta z^k = \left[1 + \sum_{j=\phi+2}^{k-1} \rho_j\right]\sigma, \\ |\Delta \mathbf{s}^k - [\mathbf{e}^T \Delta \mathbf{s}^k]\,\mathbf{e}| &= |\mathbf{e}\,\Delta z^k - [\mathbf{e}^T \mathbf{e}\,\Delta z^k]\,\mathbf{e}| = 0, \\ \left|\frac{\mathbf{s}_1^k + \mathbf{s}_2^k}{2}\right| &= \left|\frac{\mathbf{s}^{\phi} + \mathbf{e}\,z_1^k + \mathbf{s}^{\phi} + \mathbf{e}\,z_2^k}{2}\right| \leqslant |\mathbf{s}^{\phi}| + |\breve{z}^k| \\ &< \sigma\frac{\mathbf{e}^T[\mathbf{D} - \mathbf{C}]\,\mathbf{e}}{4\|\mathbf{C}\|} + \sigma = \sigma\left\{\frac{\mathbf{e}^T[\mathbf{D} - \mathbf{C}]\,\mathbf{e}}{4\|\mathbf{C}\|} + 1\right\} \end{aligned} \tag{S3.52}$$

Hence, for an arbitrary monomorphic state $\mathcal{S}$ in $\mathcal{M}^c$, we can find a trait substitution sequence that starts from $\mathcal{S}$ and enters $\mathcal{D}^c$ (equation (S2.5)), by choosing a sufficiently large $K$ so that we can find values for $c$ and $d$ satisfying $\left|\frac{\mathbf{s}_1^k+\mathbf{s}_2^k}{2}\right| < d \ll c < |\mathbf{e}^T\,\Delta\,\mathbf{s}^k|$. Therefore, there is a non-zero probability for a trait substitution sequence starting from an arbitrary state in $\mathcal{M}^c$ to enter $\mathcal{D}^c$, and hence equation (S2.10) holds.

##### S3.4.2 Derivation of equations (S3.46) and (S3.47)

Substitution of $\mathbf{s}'=\mathbf{s}_\phi+z'\,\mathbf{e}$, $\mathbf{s}_i=\mathbf{s}_\phi+z_i\,\mathbf{e}$ into equation (S1.9) with $\mathbf{g}=\mathbf{0}$ gives

$$\begin{aligned}\tilde{F}(z';(z_1,...,z_M),(n_1,...,n_M)) &= F(\mathbf{s}^\phi+\mathbf{e}\,z';(\mathbf{s}^\phi+\mathbf{e}\,z_i,...,\mathbf{s}^\phi+\mathbf{e}\,z_M),(n_1,...,n_M))\\ &= [\mathbf{s}^\phi+\mathbf{e}\,\bar{z}]^T\,\mathbf{C}\,\mathbf{e}[z'-\bar{z}]+\frac{1}{2}\mathbf{e}^T\,\mathbf{D}\,\mathbf{e}[z'-\bar{z}]^2-\frac{1}{2}\mathbf{e}^T\,\mathbf{D}\,\mathbf{e}\,V_z+\bar{F}\\ &= \tilde{g}[z'-\bar{z}]+\tilde{C}\,z[z'-\bar{z}]+\frac{1}{2}\tilde{D}[z'-\bar{z}]^2-\frac{1}{2}\tilde{D}\,V_z+\bar{F} \qquad \text{(S3.53)}\end{aligned}$$

with $\tilde{g}=\mathbf{s}^{\phi T}\,\mathbf{C}\,\mathbf{e}$, $\tilde{C}=\mathbf{e}^T\,\mathbf{C}\,\mathbf{e}$, $\tilde{D}=\mathbf{e}^T\,\mathbf{D}\,\mathbf{e}$, $\bar{z}=\sum_{i=1}^M p_i\,z_i$ and $V_z=\sum_{i=1}^M p_i[z_i-\bar{z}]^2$, where the higher order terms denoted by $O(\varepsilon^3)$ are neglected. Note that the coexistence of more than two resident phenotypes at population dynamical equilibrium is impossible, because equation (S3.53) is a parabolic function of $z'$ with a constant shape determined by $\tilde{D}$.

From equation (S3.53) we derive the invasion fitness for a mutant $z'$ under a monomorphic resident $z$ as

$$\begin{aligned}\tilde{f}(z';z) &= \tilde{F}(z';z,\hat{n})\\ &= \tilde{g}[z'-z]+\tilde{C}\,z[z'-z]+\frac{1}{2}\tilde{D}[z'-z]^2. \qquad \text{(S3.54)}\end{aligned}$$

Also from equations (S3.53), (S3.17), and (S3.20), we see that the invasion fitness for a mutant $z'$ under dimorphic residents $z_1$ and $z_2$ can be expressed as

$$\begin{aligned}\tilde{f}(z';z_1,z_2) &= \tilde{F}(z';(z_1,z_2),(\hat{n}_1,\hat{n}_2))\\ &= [\tilde{g}+\tilde{C}\,\bar{z}][z'-\bar{z}]+\frac{\tilde{D}}{2}\{[z'-\bar{z}]^2-\hat{p}_1[z_1-\bar{z}]^2-\hat{p}_2[z_2-\bar{z}]^2\},\\ \hat{p}_1 &= \frac{1}{2}+B,\\ \hat{p}_2 &= \frac{1}{2}-B,\\ B &= \frac{\breve{\mathbf{s}}^T\,\mathbf{C}\,\Delta\,\mathbf{s}}{\Delta\,\mathbf{s}^T[\mathbf{D}-\mathbf{C}]\,\Delta\,\mathbf{s}}=\frac{\left[\frac{\mathbf{s}_1+\mathbf{s}_2}{2}\right]^T\mathbf{C}[\mathbf{s}_1-\mathbf{s}_2]}{[\mathbf{s}_1-\mathbf{s}_2]^T[\mathbf{D}-\mathbf{C}][\mathbf{s}_1-\mathbf{s}_2]}\\ &= \frac{\left[\frac{\mathbf{s}^\phi+\mathbf{e}\,z_1+\mathbf{s}^\phi+\mathbf{e}\,z_2}{2}\right]^T\mathbf{C}\,\mathbf{e}[z_1-z_2]}{[z_1-z_2]\,\mathbf{e}^T[\mathbf{D}-\mathbf{C}]\,\mathbf{e}[z_1-z_2]}=\frac{\left[\mathbf{s}^\phi+\mathbf{e}\,\frac{z_1+z_2}{2}\right]^T\mathbf{C}\,\mathbf{e}}{[z_1-z_2]\,\mathbf{e}^T[\mathbf{D}-\mathbf{C}]\,\mathbf{e}}=\frac{\mathbf{s}^{\phi T}\,\mathbf{C}\,\mathbf{e}+\breve{z}\,\mathbf{e}^T\,\mathbf{C}\,\mathbf{e}}{\Delta\,z\,\mathbf{e}^T[\mathbf{D}-\mathbf{C}]\,\mathbf{e}}\\ &= \frac{\tilde{g}+\tilde{C}\,\breve{z}}{\Delta\,z[\tilde{D}-\tilde{C}]},\\ \bar{z} &= \hat{p}_1\,z_1+\hat{p}_2\,z_2=\breve{z}+B\,\Delta\,z=\breve{z}+\frac{\tilde{g}+\tilde{C}\,\breve{z}}{\tilde{D}-\tilde{C}}=\frac{\tilde{g}+\tilde{D}\,\breve{z}}{\tilde{D}-\tilde{C}}. \qquad \text{(S3.55)}\end{aligned}$$

We transform equation (S3.55) as

$$\begin{aligned}\tilde{f}(z';z_1,z_2) &= [\tilde{g}+\tilde{C}\,\bar{z}][z'-\bar{z}]+\frac{\tilde{D}}{2}\{z'^2-2\,z'\,\bar{z}-\hat{p}_1\,z_1^2-\hat{p}_2\,z_2^2+2\,\bar{z}^2\}\\ &= \frac{\tilde{D}}{2}z'^2+\{\tilde{g}+[\tilde{C}-\tilde{D}]\,\bar{z}\}\,z'-\{\tilde{g}+[\tilde{C}-\tilde{D}]\,\bar{z}\}\,\bar{z}-\frac{\tilde{D}}{2}\{\hat{p}_1\,z_1^2+\hat{p}_2\,z_2^2\}\\ &= \frac{\tilde{D}}{2}z'^2+\{\tilde{g}-[\tilde{g}+\tilde{D}\,\breve{z}]\}\,z'-\{\tilde{g}-[\tilde{g}+\tilde{D}\,\breve{z}]\}\,\bar{z}-\frac{\tilde{D}}{2}\left\{\hat{p}_1\left[\breve{z}+\frac{\Delta\,z}{2}\right]^2+\hat{p}_2\left[\breve{z}-\frac{\Delta\,z}{2}\right]^2\right\}\end{aligned}$$

$$\begin{aligned}
&= \frac{\tilde{D}}{2}z'^2-\tilde{D}\,\breve{z}\,z'+\tilde{D}\,\breve{z}\,\bar{z}-\frac{\tilde{D}}{2}\left\{\breve{z}^2+\frac{\Delta z^2}{4}+[\hat{p}_1-\hat{p}_2]\,\breve{z}\,\Delta z\right\}\\
&= \frac{\tilde{D}}{2}z'^2-\tilde{D}\,\breve{z}\,z'+\tilde{D}\,\breve{z}[\breve{z}+B\,\Delta z]-\frac{\tilde{D}}{2}\left\{\breve{z}^2+\frac{\Delta z^2}{4}+2\,B\,\breve{z}\,\Delta z\right\}\\
&= \frac{\tilde{D}}{2}z'^2-\tilde{D}\,\breve{z}\,z'+\frac{\tilde{D}}{2}\left\{\breve{z}^2-\frac{\Delta z^2}{4}\right\}\\
&= \frac{\tilde{D}}{2}\{z'^2-[z_1+z_2]\,z'+z_1\,z_2\}=\frac{\tilde{D}}{2}[z'-z_1][z'-z_2].
\end{aligned}\tag{S3.56}$$

##### S3.4.3 Residents after invasion by $z'^k$ for $k=\phi+1$

First, if $z'^{\phi}=\sigma\,/\,2$ excludes its parental resident, $z^{\phi}=0$, then the next invading mutant, $z'^{\phi+1}=-\sigma/2$, coexists with $z'^{\phi}$, because equations (S2.43) and (S2.46) give

$$\begin{aligned}
\tilde{f}(z'^{\phi+1};z'^{\phi}) &= -\tilde{g}\,\sigma+\frac{1}{2}[\tilde{D}-\tilde{C}]\,\sigma^2=-\mathbf{s}^{\phi T}\,\mathbf{C}\,\mathbf{e}\,\sigma+\frac{1}{2}[\tilde{D}-\tilde{C}]\,\sigma^2\\
&\geqslant \|\mathbf{C}\|\,\sigma\left\{-|\mathbf{s}^{\phi}|+\frac{\sigma[\tilde{D}-\tilde{C}]}{2\|\mathbf{C}\|}\right\}\\
&\geqslant \|\mathbf{C}\|\,\sigma\left\{-\frac{\sigma[\tilde{D}-\tilde{C}]}{4\|\mathbf{C}\|}+\frac{\sigma[\tilde{D}-\tilde{C}]}{2\|\mathbf{C}\|}\right\}=\frac{\sigma^2[\tilde{D}-\tilde{C}]}{4}>0,\\
\tilde{f}(z'^{\phi};z'^{\phi+1}) &= \tilde{g}\,\sigma+\frac{1}{2}[\tilde{D}-\tilde{C}]\,\sigma^2\\
&\geqslant \|\mathbf{C}\|\,\sigma\left\{-|\mathbf{s}^{\phi}|+\frac{\sigma[\tilde{D}-\tilde{C}]}{2\|\mathbf{C}\|}\right\}\geqslant\frac{\sigma^2[\tilde{D}-\tilde{C}]}{4}>0.
\end{aligned}\tag{S3.57}$$

Second, if $z'^{\phi}=\sigma\,/\,2$ coexists with its parental resident, $z^{\phi}=0$, then the next invading mutant, $z'^{\phi+1}=-\sigma/2$ , excludes $z^{\phi}$ and coexists with $z'^{\phi}$, because we see from equation (S3.47) that

$$\begin{aligned}
\tilde{f}(z'^{\phi+1};z^{\phi},z'^{\phi}) &= \frac{\tilde{D}}{2}\Big[-\frac{\sigma}{2}-0\Big]\Big[-\frac{\sigma}{2}-\frac{\sigma}{2}\Big]=\frac{\tilde{D}}{4}\sigma^2>0,\\
\tilde{f}(z^{\phi};z'^{\phi},z'^{\phi+1}) &= \frac{\tilde{D}}{2}\Big[0-\frac{\sigma}{2}\Big]\Big[0+\frac{\sigma}{2}\Big]=-\frac{\tilde{D}}{8}\sigma^2<0
\end{aligned}\tag{S3.58}$$

and equation (S3.57) hold.

Therefore, the residents after the invasion by $z'^{\phi+1}=-\sigma/2$ are given by $z_1^{\phi+2}=z'^{\phi}=\sigma/2$ and $z_2^{\phi+2}=z'^{\phi+1}=-\sigma/2$.

##### S3.4.4 Residents after invasion by $z'^k$ for $k=\phi+2,...,K-1$

From equations (S3.47) and (S3.49) we derive the following relationships for an arbitrary $k\geqslant\phi+2$ under $z_1^k-z_2^k>0$:

$$\begin{aligned}
\tilde{f}(z'^k;z_1^k,z_2^k) &= \frac{\tilde{D}}{2}[z'^k-z_1^k][z'^k-z_2^k]\\
&= \begin{cases}\frac{\tilde{D}}{2}[z_1^k+\rho_k\,\sigma-z_1^k][z_1^k+\rho_k\,\sigma-z_2^k] & \text{for } |z_1^k|\leqslant|z_2^k|\\ \frac{\tilde{D}}{2}[z_2^k-\rho_k\,\sigma-z_1^k][z_2^k-\rho_k\,\sigma-z_2^k] & \text{for otherwise}\end{cases}\\
&= \frac{\tilde{D}}{2}\,\rho_k\,\sigma[z_1^k-z_2^k+\rho_k\,\sigma]>0,
\end{aligned}$$

$$\begin{aligned}
\tilde f(z_1^k; z_2^k, z'^k) &= \frac{\tilde D}{2}[z_1^k - z_2^k][z_1^k - z'^k] \\
&= \begin{cases} \frac{\tilde D}{2}[z_1^k - z_2^k][z_1^k - z_1^k - \rho_k\,\sigma] & \text{for } |z_1^k| \leqslant |z_2^k| \\ \frac{\tilde D}{2}[z_1^k - z_2^k][z_1^k - z_2^k + \rho_k\,\sigma] & \text{for otherwise} \end{cases} \\
&= \begin{cases} -\frac{\tilde D}{2}[z_1^k - z_2^k]\,\rho_k\,\sigma < 0 & \text{for } |z_1^k| \leqslant |z_2^k| \\ \frac{\tilde D}{2}[z_1^k - z_2^k][z_1^k - z_2^k + \rho_k\,\sigma] > 0 & \text{for otherwise,} \end{cases} \\
\tilde f(z_2^k; z_1^k, z'^k) &= \frac{\tilde D}{2}[z_2^k - z_1^k][z_2^k - z'^k] \\
&= \begin{cases} \frac{\tilde D}{2}[z_1^k - z_2^k][z_1^k - z_2^k + \rho_k\,\sigma] > 0 & \text{for } |z_1^k| \leqslant |z_2^k| \\ -\frac{\tilde D}{2}[z_1^k - z_2^k]\,\rho_k\,\sigma < 0 & \text{for otherwise.} \end{cases}
\end{aligned} \tag{S3.59}$$

Hence, if $z'^k$ derives from $z_1^k$ (i.e., $z'^k = z_1^k + \rho_k\,\sigma$), then $z'^k$ excludes $z_1^k$ and coexists with $z_2^k$, establishing the next dimorphic residents $z_1^{k+1} = z'^k$ and $z_2^{k+1} = z_2^k$. Conversely, if $z'^k$ derives from $z_2^k$ (i.e., $z'^k = z_2^k - \rho_k\,\sigma$), then $z'^k$ excludes $z_2^k$ and coexists with $z_1^k$, establishing the next dimorphic residents $z_1^{k+1} = z_1^k$ and $z_2^{k+1} = z'^k$. The coexistence of $z_1^{k+1}$ and $z_2^{k+1}$ is warranted, because their fractions are given by $\hat p_1^{k+1} = \frac{1}{2} + B_{k+1}$ and $\hat p_2^{k+1} = \frac{1}{2} - B_{k+1}$ with

$$|B_{k+1}| = \frac{[\mathbf{s}^\phi + \mathbf{e}\,\breve z^{k+1}]^T\,\mathbf{C}\,\mathbf{e}\,\Delta\,z^{k+1}}{\mathbf{e}^T[\mathbf{D} - \mathbf{C}]\,\mathbf{e}\,[\Delta\,z^{k+1}]^2} < \frac{1}{2}, \tag{S3.60}$$

for an arbitrary $k \geqslant \phi + 2$ (derived in Section S2.4.5).

Therefore, each invading mutant $z'^k$ for $k = \phi + 2, \ldots, K - 1$ excludes its parental resident among $z_1^k$ and $z_2^k$ and coexists with the other resident (i.e., $z'^k = z_1^k + \rho_k\,\sigma$ gives $z_1^{k+1} = z'^k$ and $z_2^{k+1} = z_2^k$, whereas $z'^k = z_2^k - \rho_k\,\sigma$ gives $z_1^{k+1} = z_1^k$ and $z_2^{k+1} = z'^k$).

##### S3.4.5 Derivation of equation (S3.60)

First, if $|\tilde C|/[\tilde D - \tilde C] \leqslant 1$ holds, i.e., equation (S2.49) gives $\rho_k = 1$ for $k \geqslant \phi + 2$, then we see from equations (S2.43), (S2.50), and (S2.55) that for $k \geqslant \phi + 2$

$$\begin{aligned}
|B_{k+1}| &\leqslant \frac{|\tilde g| + |\tilde C||\breve z^{k+1}|}{\Delta\,z[\tilde D - \tilde C]} = \frac{|\mathbf{s}^{\phi T}\,\mathbf{C}\,\mathbf{e}|}{[\tilde D - \tilde C]\,\Delta\,z^{k+1}} + \frac{|\tilde C||\breve z^{k+1}|}{[\tilde D - \tilde C]\,\Delta\,z^{k+1}} \\
&< \frac{|\mathbf{s}^\phi|\|\mathbf{C}\|}{[\tilde D - \tilde C]\,\sigma} + \frac{|\tilde C|\frac{\sigma}{2}}{[\tilde D - \tilde C][\Delta\,z^k + \rho_k\,\sigma]} \\
&< \frac{1}{4} + \frac{|\tilde C|\frac{\sigma}{2}}{[\tilde D - \tilde C]\,2\,\sigma} < \frac{1}{4} + \frac{|\tilde C|}{4[\tilde D - \tilde C]} < \frac{1}{2}.
\end{aligned} \tag{S3.61}$$

Second, if $|\tilde C|/[\tilde D - \tilde C] > 1$ and $\frac{|\tilde C|}{[\tilde D - \tilde C]} \leqslant \frac{\Delta\,z^k + \sigma}{2\,\sigma}$ hold, (i.e., equation (S3.49) for $\frac{\Delta\,z^k}{2\,\sigma}\Big[\frac{|\tilde C|}{[\tilde D - \tilde C]} - \frac{1}{2}\Big]^{-1} \geqslant 1$ gives $\rho_k = 1$), then we see from equations (S3.43), (S3.50), and (S3.55) that for $k \geqslant \phi + 2$

$$\begin{aligned}
|B_{k+1}| &< \frac{1}{4} + \frac{|\tilde C|\,\sigma\,\rho_k}{2[\tilde D - \tilde C][\Delta\,z^k + \rho_k\,\sigma]} = \frac{1}{4} + \frac{|\tilde C|\,\sigma}{2[\tilde D - \tilde C][\Delta\,z^k + \sigma]} \\
&\leqslant \frac{1}{4} + \frac{1}{2}\Big[\frac{\Delta\,z^k + \sigma}{2\,\sigma}\Big]\frac{\sigma}{[\Delta\,z^k + \sigma]} = \frac{1}{4} + \frac{1}{4} = \frac{1}{2}.
\end{aligned} \tag{S3.62}$$

Third, if $|\tilde{C}|/[\tilde{D}-\tilde{C}]>1$ and $\frac{|\tilde{C}|}{[\tilde{D}-\tilde{C}]}>\frac{\Delta z^k+\sigma}{2\sigma}$ holds (i.e., equation (S3.49) for $\frac{\Delta z^k}{2\sigma}\Big[\frac{|\tilde{C}|}{[\tilde{D}-\tilde{C}]}-\frac{1}{2}\Big]^{-1}<1$ gives $\rho_k=\frac{\Delta z^k}{2\sigma}\Big[\frac{|\tilde{C}|}{[\tilde{D}-\tilde{C}]}-\frac{1}{2}\Big]^{-1}$), then we see from equations (S3.43), (S3.50), and (S3.55) that for $k\geqslant\phi+2$

$$\begin{aligned}|B_{k+1}| &< \frac{1}{4}+\frac{|\tilde{C}|\,\rho_k\,\sigma}{2[\tilde{D}-\tilde{C}][\Delta z^k+\rho_k\,\sigma]}\\ &= \frac{1}{4}+\frac{|\tilde{C}|\frac{\Delta z^k}{2\sigma}\Big[\frac{|\tilde{C}|}{[\tilde{D}-\tilde{C}]}-\frac{1}{2}\Big]^{-1}\sigma}{2[\tilde{D}-\tilde{C}]\Big[\Delta z^k+\frac{\Delta z^k}{2\sigma}\Big[\frac{|\tilde{C}|}{[\tilde{D}-\tilde{C}]}-\frac{1}{2}\Big]^{-1}\sigma\Big]}\\ &= \frac{1}{4}+\frac{1}{4}=\frac{1}{2}. \end{aligned}\qquad\text{(S3.63)}$$

#### S3.5. Invariance of fraction dynamics against scaling of trait space

We see from equation (S2.9) that the fraction dynamics for mutant $\mathbf{s}'$ under existence of $\mathbf{s}_1,\ldots,\mathbf{s}_M$ with their population sizes $n_1,\ldots,n_M$ is given by

$$\begin{aligned}\frac{d\log p'}{dt} &= \frac{d\log(n'/N)}{dt}=\frac{1}{n'}\frac{dn'}{dt}-\frac{1}{N}\frac{dN}{dt}=\frac{1}{n'}\frac{dn'}{dt}-\sum_{j=1}^{M}\frac{n_j}{N}\Big[\frac{1}{n_j}\frac{dn_j}{dt}\Big]\\ &= F(\mathbf{s}';(\mathbf{s}_1,\ldots,\mathbf{s}_M),(n_1,\ldots,n_M))-\sum_{j=1}^{N}p_j\,F(\mathbf{s}';(\mathbf{s}_1,\ldots,\mathbf{s}_M),(n_1,\ldots,n_M))\\ &= \Big\{\bar{\mathbf{s}}^T\mathbf{C}[\mathbf{s}'-\bar{\mathbf{s}}]+\frac{1}{2}[\mathbf{s}'-\bar{\mathbf{s}}]^T\mathbf{D}[\mathbf{s}'-\bar{\mathbf{s}}]-\frac{1}{2}V_{\mathbf{D}}\Big\}\\ &\quad -\sum_{j=1}^{M}p_j\Big\{\bar{\mathbf{s}}^T\mathbf{C}[\mathbf{s}'-\bar{\mathbf{s}}]+\frac{1}{2}[\mathbf{s}'-\bar{\mathbf{s}}]^T\mathbf{D}[\mathbf{s}'-\bar{\mathbf{s}}]-\frac{1}{2}V_{\mathbf{D}}\Big\}\\ &= \bar{\mathbf{s}}^T\mathbf{C}[\mathbf{s}'-\bar{\mathbf{s}}]+\frac{1}{2}[\mathbf{s}'-\bar{\mathbf{s}}]^T\mathbf{D}[\mathbf{s}'-\bar{\mathbf{s}}]-\frac{1}{2}V_{\mathbf{D}}. \end{aligned}\qquad\text{(S3.64)}$$

When a mutational covariance matrix is given by $\mathbf{V}_\mu=\tilde{\sigma}^2\mathbf{I}$ with $\tilde{\sigma}$ being different from $\sigma$, we can rescale the trait space by introducing a new coordinate system, denoted by $\tilde{\mathbf{s}}=\rho\mathbf{s}$ with $\rho=\sigma/\tilde{\sigma}$, so that the mutation size in coordinate system $\tilde{\mathbf{s}}$ becomes equal to $\sigma$. In this case, by introducing a rescaled time $\tilde{t}=\rho^{-2}t$, we can make the shape of the fitness function kept unchanged in the sense that the fraction dynamics is kept identical to equation (S2.64), satisfying

$$\begin{aligned}\frac{d\log p'}{d\tilde{t}} &= \frac{dt}{d\tilde{t}}\frac{d\log p'}{dt}=\rho^2\frac{d\log(n'/N)}{dt}=\rho^2\Big\{\frac{1}{n'}\frac{dn'}{dt}-\frac{1}{N}\frac{dN}{dt}\Big\}\\ &= \rho^2F(\tilde{\mathbf{s}}';(\tilde{\mathbf{s}}_1,\ldots,\tilde{\mathbf{s}}_M),(n_1,\ldots,n_M))-\rho^2\sum_{j=1}^{N}p_j\,F(\tilde{\mathbf{s}}_j;(\tilde{\mathbf{s}}_1,\ldots,\tilde{\mathbf{s}}_M),(n_1,\ldots,n_M))\\ &= \rho^2\Big\{\bar{\tilde{\mathbf{s}}}^T\mathbf{C}[\tilde{\mathbf{s}}'-\bar{\tilde{\mathbf{s}}}]+\frac{1}{2}[\tilde{\mathbf{s}}'-\bar{\tilde{\mathbf{s}}}]^T\mathbf{D}[\tilde{\mathbf{s}}'-\bar{\tilde{\mathbf{s}}}]-\frac{1}{2}\tilde{V}_{\mathbf{D}}+\bar{\tilde{F}}\Big\}\\ &\quad -\rho^2\sum_{j=1}^{M}p_j\Big\{\bar{\tilde{\mathbf{s}}}^T\mathbf{C}[\tilde{\mathbf{s}}_j-\bar{\tilde{\mathbf{s}}}]+\frac{1}{2}[\tilde{\mathbf{s}}_j-\bar{\tilde{\mathbf{s}}}]^T\mathbf{D}[\tilde{\mathbf{s}}_j-\bar{\tilde{\mathbf{s}}}]-\frac{1}{2}\tilde{V}_{\mathbf{D}}+\bar{\tilde{F}}\Big\}\\ &= \rho^2\Big\{\bar{\tilde{\mathbf{s}}}^T\mathbf{C}[\tilde{\mathbf{s}}'-\bar{\tilde{\mathbf{s}}}]+\frac{1}{2}[\tilde{\mathbf{s}}'-\bar{\tilde{\mathbf{s}}}]^T\mathbf{D}[\tilde{\mathbf{s}}'-\bar{\tilde{\mathbf{s}}}]-\frac{1}{2}\tilde{V}_{\mathbf{D}}\Big\} \end{aligned}\qquad\text{(S3.65)}$$

$$\begin{aligned} &= \rho^2\Bigg\{\frac{\bar{\mathbf{s}}^T}{\rho}\mathbf{C}\Big[\frac{\mathbf{s}'}{\rho}-\frac{\bar{\mathbf{s}}}{\rho}\Big]+\frac{1}{2}\Big[\frac{\mathbf{s}'}{\rho}-\frac{\bar{\mathbf{s}}}{\rho}\Big]^T\mathbf{D}\Big[\frac{\mathbf{s}'}{\rho}-\frac{\bar{\mathbf{s}}}{\rho}\Big]-\frac{1}{2}\sum_{j=1}^{M}\Big[\frac{\mathbf{s}_j}{\rho}-\frac{\bar{\mathbf{s}}}{\rho}\Big]^T\mathbf{D}\Big[\frac{\mathbf{s}_j}{\rho}-\frac{\bar{\mathbf{s}}}{\rho}\Big]\Bigg\}\\ &= \bar{\mathbf{s}}^T\mathbf{C}[\mathbf{s}'-\bar{\mathbf{s}}]+\frac{1}{2}[\mathbf{s}'-\bar{\mathbf{s}}]^T\mathbf{D}[\mathbf{s}'-\bar{\mathbf{s}}]-\frac{1}{2}V_{\mathbf{D}} \end{aligned}\qquad\text{(S3.66)}$$

with

$$
\begin{aligned}
\bar{\bar{\mathbf{s}}} &= \sum_{j=1}^{M} p_j\,\tilde{\mathbf{s}}_j = \rho^{-1}\,\bar{\mathbf{s}}, \\
\tilde{V}_{\mathbf{D}} &= \sum_{j=1}^{M} p_j[\tilde{\mathbf{s}}_j - \bar{\bar{\mathbf{s}}}]^T\,\mathbf{D}[\tilde{\mathbf{s}}_j - \bar{\bar{\mathbf{s}}}] = \rho^{-2}\,V_{\mathbf{D}}, \\
\bar{\bar{F}} &= \sum_{j=1}^{M} p_j\,F(\tilde{\mathbf{s}}'; (\tilde{\mathbf{s}}_1, ..., \tilde{\mathbf{s}}_M), (n_1, ..., n_M)).
\end{aligned}
\tag{S3.67}
$$

Hence, replacing $\sigma$ with $\tilde{\sigma}$ in the definition of $\mathcal{M}^c$ and $\mathcal{D}^c$ in equations (S3.1) and (S3.5) keeps the situation identical to that under the original mutation size, $\sigma$, which gives

$$
\begin{aligned}
P_{\min}^{\mathrm{mc}}(\tilde{\sigma}) &= P_{\min}^{\mathrm{mc}}(\sigma) = 1 - \varepsilon_{\mathrm{mc}}, \\
P_{\min}^{\mathrm{mc}\to\mathrm{dd}}(\tilde{\sigma}) &= P_{\min}^{\mathrm{mc}\to\mathrm{dd}}(\sigma) \\
P_{\min}^{\mathrm{dd}}(\tilde{\sigma}) &= P_{\min}^{\mathrm{dd}}(\sigma) = 1 - \varepsilon_{\mathrm{dd}}.
\end{aligned}
\tag{S3.68}
$$

Therefore, $\varepsilon_{\mathrm{mc}}$ and $\varepsilon_{\mathrm{dd}}$ must be independent of the mutation size, and hence are determined only by $c$, $d$, $\mathbf{C}$, and $\mathbf{D}$.

#### S3.6. Evaluation of waiting time for evolutionary branching in numerically simulated evolution

For quantitative evaluation of the waiting time for evolutionary branching, we consider a trait substitution sequence $\mathfrak{B}(\mathbf{s}_0) = (\mathcal{S}^0, ..., \mathcal{S}^K)$ that starts from a monomorphic resident $\mathcal{S}^0 = \{\mathbf{s}_0\}$ and ends when the system comes to have dimorphic residents $\mathcal{S}^K = \{\mathbf{s}_1^K, \mathbf{s}_2^K\}$ with their phenotypic distance attaining $\Delta\, s$ (i.e., $|\mathbf{s}_1 - \mathbf{s}_2| \geqslant \Delta\, s$). We evaluate the waiting time for evolutionary branching by a “relative path length” of $\mathfrak{B}(\mathbf{s}_0)$, which is the product of the mutational step size and the number of invasions, $\sigma\, K$, divided by the path length estimated from the canonical equation denoted by $l_{\mathrm{CE}}(\mathbf{s}_0, \Delta\, s)$:

$$
\tilde{l}\,(\mathbf{s}_0, \Delta\, s; \sigma) = \frac{l_{\mathfrak{B}}^{\mathrm{mc}} + l_{\mathfrak{B}}^{\mathrm{mc}\to\mathrm{dd}} + l_{\mathfrak{B}}^{\mathrm{dd}}}{l_{\mathrm{CE}}(\mathbf{s}_0, \Delta\, s)} = \frac{\sigma\, K}{l_{\mathrm{CE}}(\mathbf{s}_0, \Delta\, s)},
\tag{S3.69}
$$

where $l_{\mathfrak{B}}^{\mathrm{mc}} = \sigma\, K^{\mathrm{mc}}$, $l_{\mathfrak{B}}^{\mathrm{dd}} = \sigma[K - K^{\mathrm{dd}}]$, and $l_{\mathfrak{B}}^{\mathrm{mc}\to\mathrm{dd}} = \sigma[K^{\mathrm{dd}} - K^{\mathrm{mc}}]$ with $l_{\mathfrak{B}}^{\mathrm{mc}} + l_{\mathfrak{B}}^{\mathrm{mc}\to\mathrm{dd}} + l_{\mathfrak{B}}^{\mathrm{mc}} = \sigma\; K$ describe the path lengths for m.c. (i.e, monomorphic convergence), d.d. (i.e., dimorphic divergence), and transition from m.c. to d.d., respectively, defined in Section S2.1 (satisfying $K^{\mathrm{mc}} = \min_{\mathcal{S}^K \in \mathcal{M}^c}(K)$ and $K^{\mathrm{dd}} = \min_{\mathcal{S}^K \in \mathcal{D}^c}(K)$). $l_{\mathrm{CE}}(\mathbf{s}_0, \Delta\, s)$ describes the path length of the deterministic trajectory in the trait space, formed by connecting m.c. and d.d. described with the canonical equation, given by

$$
\begin{aligned}
l_{\mathrm{CE}}(\mathbf{s}_0, \Delta\, s) &= l_{\mathrm{CE}}^{\mathrm{mc}}(\mathbf{s}_0) + l_{\mathrm{CE}}^{\mathrm{dd}}(\Delta\, s), \\
l_{\mathrm{CE}}^{\mathrm{mc}}(\mathbf{s}_0) &= \int_0^{\infty} \frac{d\mathbf{s}}{\mathrm{dt}}\,\mathrm{dt}, \\
l_{\mathrm{CE}}^{\mathrm{dd}}(\Delta\, s) &= \int_0^{T} \left[\frac{d\mathbf{s}_1}{\mathrm{dt}} - \frac{d\mathbf{s}_2}{\mathrm{dt}}\right] \mathrm{dt}
\end{aligned}
\tag{S3.70}
$$

with $\mathbf{s}(0) = \mathbf{s}_0$ and $|\mathbf{s}_1(T) - \mathbf{s}_2(T)| = \Delta\, s$. Note that

$$
\begin{aligned}
\lim_{\sigma\to 0} E\,(l_{\mathfrak{B}}^{\mathrm{mc}}) &= l_{\mathrm{CE}}^{\mathrm{mc}}(\mathbf{s}_0), \\
\lim_{\sigma\to 0} E\,(l_{\mathfrak{B}}^{\mathrm{dd}}) &= l_{\mathrm{CE}}^{\mathrm{dd}}(\Delta\, s)
\end{aligned}
\tag{S3.71}
$$

hold.

If the branching possibility conditions ensure that we can find a positive constant, denoted by $c_{\mathrm{dd}\rightarrow\mathrm{mc}}$, satisfying

$$E\left(K^{\mathrm{dd}}-K^{\mathrm{mc}};\sigma\right) \;=\; c_{\mathrm{dd}\rightarrow\mathrm{mc}}, \tag{S3.72}$$

then the scaling invariance shown in Section S2.5 gives

$$\lim_{\sigma\rightarrow 0} E\left(K^{\mathrm{dd}}-K^{\mathrm{mc}};\sigma\right) \;=\; c_{\mathrm{dd}\rightarrow\mathrm{mc}}, \tag{S3.73}$$

resulting in

$$\lim_{\sigma\rightarrow 0} l_{\mathfrak{B}}^{\mathrm{mc}\rightarrow\mathrm{dd}} \;=\; \lim_{\sigma\rightarrow 0}\left[K^{\mathrm{dd}}-K^{\mathrm{mc}}\right]\sigma = \lim_{\sigma\rightarrow 0} c_{\mathrm{dd}\rightarrow\mathrm{mc}}\,\sigma = 0. \tag{S3.74}$$

Hence, we see from equations (S2.69), (S2.71) and (S2.74) that

$$\begin{aligned}\lim_{\sigma\rightarrow 0} E(\tilde{l}(\mathbf{s}_0,\Delta s;\sigma)) &= \lim_{\sigma\rightarrow 0} E\left(\frac{l_{\mathfrak{B}}^{\mathrm{mc}}+l_{\mathfrak{B}}^{\mathrm{mc}\rightarrow\mathrm{dd}}+l_{\mathfrak{B}}^{\mathrm{mc}}}{l_{\mathrm{CE}}(\mathbf{s}_0,\Delta s)}\right)\\ &= \frac{\lim_{\sigma\rightarrow 0} E\left(l_{\mathfrak{B}}^{\mathrm{mc}}\right)+\lim_{\sigma\rightarrow 0} l_{\mathfrak{B}}^{\mathrm{mc}\rightarrow\mathrm{dd}}+\lim_{\sigma\rightarrow 0} E\left(l_{\mathfrak{B}}^{\mathrm{dd}}\right)}{l_{\mathrm{CE}}(\mathbf{s}_0,\Delta s)}\\ &= \frac{l_{\mathrm{CE}}^{\mathrm{mc}}(\mathbf{s}_0)+l_{\mathrm{CE}}^{\mathrm{dd}}(\Delta s)}{l_{\mathrm{CE}}^{\mathrm{mc}}(\mathbf{s}_0)+l_{\mathrm{CE}}^{\mathrm{dd}}(\Delta s)} = 1. \end{aligned}\tag{S3.75}$$

However, we do not know whether the branching possibility conditions ensure equation (S.3.72). Still, we see from Section S2.4 that

$$\begin{aligned}\Pr\left(K^{\mathrm{dd}}-K^{\mathrm{mc}}\leqslant\eta\, c_{\mathrm{dd}\rightarrow\mathrm{mc}};\sigma\right) &> 0,\\ \Pr\left(l_{\mathfrak{B}}^{\mathrm{mc}}\leqslant\eta\, L_{\mathrm{CE}}^{\mathrm{mc}}(\mathbf{s}_0);\sigma\right) &> 0,\\ \Pr\left(l_{\mathfrak{B}}^{\mathrm{dd}}\leqslant\eta\, L_{\mathrm{CE}}^{\mathrm{dd}}(\Delta s);\sigma\right) &> 0\end{aligned}\tag{S3.76}$$

with a constant $\eta$ satisfying $\eta\geqslant 1$, resulting in

$$\begin{aligned}\Pr(\tilde{l}(\mathbf{s}_0,\Delta s;\sigma)\leqslant\eta) &= \Pr\left(\frac{l_{\mathfrak{B}}^{\mathrm{mc}}+l_{\mathfrak{B}}^{\mathrm{mc}\rightarrow\mathrm{dd}}+l_{\mathfrak{B}}^{\mathrm{dd}}}{l_{\mathrm{CE}}(\mathbf{s}_0,\Delta s)}\leqslant\eta\right)\\ &= \Pr\left(\frac{l_{\mathfrak{B}}^{\mathrm{mc}}+\left[K^{\mathrm{dd}}-K^{\mathrm{mc}}\right]\sigma+l_{\mathfrak{B}}^{\mathrm{dd}}}{l_{\mathrm{CE}}^{\mathrm{mc}}(\mathbf{s}_0)+l_{\mathrm{CE}}^{\mathrm{dd}}(\Delta s)}\leqslant\eta\right)>0,\end{aligned}\tag{S3.77}$$

meaning that a non-zero fraction of all possible trait substitution sequences has evolutionary branching before their relative path lengths $\tilde{l}(\mathbf{s}_0,\Delta s;\sigma)$ exceed $\eta$.

In figures S4-S8 in Section S6, the numerically calculated $\tilde{l}(\mathbf{s}_0,\Delta s;\sigma)$ is close to 1 as expected by equation (S2.75), under the existence of a possible branching point with a sufficiently small $\lambda_{\max}(\mathbf{C})$ (panel (b)) or a strongly possible branching point with a sufficiently small $\lambda_{\max}([\mathbf{C}+\mathbf{C}^T]/2)$ (panel (a)).

#### S3.7. Branching possibility conditions for non-normalized trait spaces

We assume a non-normalized trait space $\mathbf{s}$, where the mutational covariance matrix, $\mathbf{V}_\mu$, is not proportional to the identity. Following equation (S2.10), we approximate the invasion fitness for a mutant $\mathbf{s}'$ under a monomorphic resident $\mathbf{s}$ close to an evolutionarily singular point $\mathbf{s}^*=\mathbf{0}$ as

$$f(\mathbf{s}';\mathbf{s}) \;=\; \mathbf{s}^T\mathbf{C}[\mathbf{s}'-\mathbf{s}]+\frac{1}{2}[\mathbf{s}'-\mathbf{s}]^T\mathbf{D}[\mathbf{s}'-\mathbf{s}]. \tag{S3.78}$$

Following Ito and Sasaki (2020), we introduce a locally normalized coordinate system $\mathbf{u}=(w,z)^T$ (these variables are unrelated to those denoted by the same letters in Section 2.3),

$$\mathbf{s} \;=\; \mathbf{W}\,\mathbf{u}, \tag{S3.79}$$

such that the mutational covariance matrix in coordinate system $\mathbf{u}$ is given by $\sigma_\mu^2\,\mathbf{I}$ (with $\mathbf{I}$ describing the identity matrix and $\sigma_\mu^2 = \lambda_{\max}(\mathbf{V}_\mu)$), satisfying

$$\mathbf{s}^T\,\mathbf{V}_\mu^{-1}\,\mathbf{s} \;=\; \mathbf{u}^T\,\mathbf{W}^T\,\mathbf{V}_\mu^{-1}\,\mathbf{W}\,\mathbf{u} = \sigma_\mu^{-2}\,\mathbf{u}^T\,\mathbf{u}, \tag{S3.80}$$

in which case a mutation probability distribution defined with a multivariate Gaussian function,

$$P_\mu(\mathbf{s}'-\mathbf{s}_i) \;=\; \frac{1}{\sqrt{[2\,\pi]^L|\mathbf{V}_\mu|}}\exp\!\left(-\frac{1}{2}[\mathbf{s}'-\mathbf{s}_i]^T\,\mathbf{V}_\mu^{-1}[\mathbf{s}'-\mathbf{s}_i]\right), \tag{S3.81}$$

is transformed into a isotropic Gaussian function,

$$\tilde{P}_\mu(\mathbf{u}'-\mathbf{u}_i) \;=\; \frac{1}{2\,\pi\,\sigma_\mu^2}\exp\!\left(-\frac{1}{2\,\sigma_\mu^2}|\mathbf{u}'-\mathbf{u}_i|^2\right). \tag{S3.82}$$

Such a $\mathbf{W}$ is obtained by diagonalizing $\mathbf{V}_\mu$ as

$$\begin{aligned}
\mathbf{V}_\mu &= \mathbf{E}\,\mathrm{diag}(\lambda_{\mu 1},...,\lambda_{\mu L})\,\mathbf{E}^T \\
&= [\sigma_\mu\,\mathrm{diag}(\rho_1,...,\rho_L)\,\mathbf{E}^T]^T[\sigma_\mu\,\mathrm{diag}(\rho_1,...,\rho_L)\,\mathbf{E}^T] \\
&= \sqrt{\mathbf{V}_\mu}^T\,\sqrt{\mathbf{V}_\mu} = \sigma_\mu^2\,\mathbf{W}\,\mathbf{W}^T \\
\sqrt{\mathbf{V}_\mu} &= \sigma_\mu\,\mathrm{diag}(\rho_1,...,\rho_L)\,\mathbf{E}^T \\
\mathbf{W} &= \sigma_\mu^{-1}\,\sqrt{\mathbf{V}_\mu}^T,
\end{aligned} \tag{S3.83}$$

where $\mathrm{diag}(\lambda_{\mu 1},\,...,\,\lambda_{\mu L})$ is an $L\times L$ diagonal matrix with its diagonal entries given by the eigenvalues $\lambda_{\mu 1}$, ..., $\lambda_{\mu L}$ of $\mathbf{V}_\mu$, and where $\sigma_\mu = \sqrt{\lambda_{\max}(\mathbf{V}_\mu)}$ and $(\rho_1,\,...,\,\rho_L) = \big(\sqrt{\lambda_{\mu 1}}\,/\,\sigma_\mu,\,...,\,\sqrt{\lambda_{\mu L}}\,/\,\sigma_\mu\big)$. We can easily confirm that $\mathbf{W}^T\,\mathbf{V}_\mu^{-1}\,\mathbf{W} = \sigma_\mu^{-2}\,\mathbf{W}^T\,\mathbf{W}^{T-1}\,\mathbf{W}^{-1}\,\mathbf{W} = \sigma_\mu^{-2}\,\mathbf{I}$ gives

$$\begin{aligned}
\tilde{P}_\mu(\mathbf{u}'-\mathbf{u}_i) &= P_\mu(\mathbf{W}\,\mathbf{u}'-\mathbf{W}\,\mathbf{u}_i)\left|\frac{d\mathbf{s}}{d\mathbf{u}}\right| \\
&= \frac{|\mathbf{W}|}{\sqrt{[2\,\pi]^L|\mathbf{V}_\mu|}}\exp\!\left(-\frac{1}{2}[\mathbf{u}'-\mathbf{u}_i]\,\mathbf{W}^T\,\mathbf{V}_\mu^{-1}\,\mathbf{W}[\mathbf{u}'-\mathbf{u}_i]\right) \\
&= \frac{|\mathbf{W}|}{\sqrt{[2\,\pi]^L|\sigma_\mu^2\,\mathbf{W}\,\mathbf{W}^T|}}\exp\!\left(-\frac{1}{2}[\mathbf{u}'-\mathbf{u}_i]\,\sigma_\mu^{-2}\,\mathbf{I}[\mathbf{u}'-\mathbf{u}_i]\right) \\
&= \frac{1}{\sqrt{[2\,\pi]^L\,\sigma_\mu^2}}\exp\!\left(-\frac{1}{2\,\sigma_\mu^2}|\mathbf{u}'-\mathbf{u}_i|^2\right).
\end{aligned} \tag{S3.84}$$

Then substitution of equation (S3.79) into equation (S3.78) gives the invasion fitness function in the normalized coordinates $\mathbf{u}$,

$$\begin{aligned}
\tilde{f}(\mathbf{u}';\mathbf{u}) &= f(\mathbf{W}\,\mathbf{u}';\mathbf{W}\,\mathbf{u}) = \mathbf{u}^T\,\mathbf{W}^T\,\mathbf{C}\,\mathbf{W}[\mathbf{u}'-\mathbf{u}] + \frac{1}{2}[\mathbf{u}'-\mathbf{u}]^T\,\mathbf{W}^T\,\mathbf{D}\,\mathbf{W}[\mathbf{u}'-\mathbf{u}] \\
&= \mathbf{u}^T\,\tilde{\mathbf{C}}[\mathbf{u}'-\mathbf{u}] + \frac{1}{2}[\mathbf{u}'-\mathbf{u}]^T\,\tilde{\mathbf{D}}[\mathbf{u}'-\mathbf{u}]
\end{aligned} \tag{S3.85}$$

with

$$\begin{aligned}
\tilde{\mathbf{D}} &= \mathbf{W}^T\,\mathbf{D}\,\mathbf{W} = \big[\sigma_\mu^{-1}\,\sqrt{\mathbf{V}_\mu}^T\big]^T\,\mathbf{D}\big[\sigma_\mu^{-1}\,\sqrt{\mathbf{V}_\mu}^T\big] = \sigma_\mu^{-2}\,\sqrt{\mathbf{V}_\mu}\,\mathbf{D}\,\sqrt{\mathbf{V}_\mu}^T \\
\tilde{\mathbf{C}} &= \mathbf{W}^T\,\mathbf{C}\,\mathbf{W} = \sigma_\mu^{-2}\,\sqrt{\mathbf{V}_\mu}\,\mathbf{C}\,\sqrt{\mathbf{V}_\mu}^T,
\end{aligned} \tag{S3.86}$$

where the direction of the dimorphic divergence in coordinates $\mathbf{u}$ is given by

$$\tilde{\mathbf{e}} \;=\; \mathbf{v}_{\max}(\tilde{\mathbf{D}}) = \mathbf{v}_{\max}\big(\sqrt{\mathbf{V}_\mu}\,\mathbf{D}\,\sqrt{\mathbf{V}_\mu}^T\big) \tag{S3.87}$$

with $\mathbf{v}_{\max}\big(\sqrt{\mathbf{V}_\mu}\,\mathbf{D}\,\sqrt{\mathbf{V}_\mu}^T\big)$ satisfying

$$\big[\sqrt{\mathbf{V}_\mu}\,\mathbf{D}\,\sqrt{\mathbf{V}_\mu}^T\big]\,\mathbf{v}_{\max}\big(\sqrt{\mathbf{V}_\mu}\,\mathbf{D}\,\sqrt{\mathbf{V}_\mu}^T\big) \;=\; \lambda_{\max}\big(\sqrt{\mathbf{V}_\mu}\,\mathbf{D}\,\sqrt{\mathbf{V}_\mu}^T\big)\,\mathbf{v}_{\max}\big(\sqrt{\mathbf{V}_\mu}\,\mathbf{D}\,\sqrt{\mathbf{V}_\mu}^T\big). \tag{S3.88}$$

By introducing a vector $\mathbf{b}=\sqrt{\mathbf{V}_\mu}^T\,\mathbf{v}_{\max}\big(\sqrt{\mathbf{V}_\mu}\,\mathbf{D}\,\sqrt{\mathbf{V}_\mu}^T\big)$, we can transform equation (S3.88) into

$$\mathbf{V}_\mu\,\mathbf{D}\,\mathbf{b} \;=\; \lambda_{\max}\big(\sqrt{\mathbf{V}_\mu}\,\mathbf{D}\,\sqrt{\mathbf{V}_\mu}^T\big)\mathbf{b}, \tag{S3.89}$$

which gives

$$\lambda_{\max}(\mathbf{V}_\mu\,\mathbf{D}) \;=\; \lambda_{\max}\big(\sqrt{\mathbf{V}_\mu}\,\mathbf{D}\,\sqrt{\mathbf{V}_\mu}^T\big), \tag{S3.90}$$

$$\mathbf{v}_{\max}(\mathbf{V}_\mu\,\mathbf{D}) \;=\; \frac{\mathbf{b}}{|\mathbf{b}|}=\frac{\sqrt{\mathbf{V}_\mu}^T\,\mathbf{v}_{\max}\big(\sqrt{\mathbf{V}_\mu}\,\mathbf{D}\,\sqrt{\mathbf{V}_\mu}^T\big)}{\big|\sqrt{\mathbf{V}_\mu}^T\,\mathbf{v}_{\max}\big(\sqrt{\mathbf{V}_\mu}\,\mathbf{D}\,\sqrt{\mathbf{V}_\mu}^T\big)\big|}. \tag{S3.91}$$

Hence, we can express the direction of the dimorphic divergence in coordinates $\mathbf{s}$ as

$$\mathbf{e} \;=\; \frac{\mathbf{W}\,\tilde{\mathbf{e}}}{|\mathbf{W}\,\tilde{\mathbf{e}}|}=\frac{\sqrt{\mathbf{V}_\mu}^T\,\mathbf{v}_{\max}\big(\sqrt{\mathbf{V}_\mu}\,\mathbf{D}\,\sqrt{\mathbf{V}_\mu}^T\big)}{\big|\sqrt{\mathbf{V}_\mu}^T\,\mathbf{v}_{\max}\big(\sqrt{\mathbf{V}_\mu}\,\mathbf{D}\,\sqrt{\mathbf{V}_\mu}^T\big)\big|}=\mathbf{v}_{\max}(\mathbf{V}_\mu\,\mathbf{D}). \tag{S3.92}$$

From $\tilde{\mathbf{e}}=\mathbf{W}^{-1}\,\mathbf{e}/|\mathbf{W}^{-1}\,\mathbf{e}|$ and equations (S3.86), we get

$$\begin{aligned}
\lambda_{\max}(\tilde{\mathbf{C}}^T) &= \sigma_\mu^{-2}\,\lambda_{\max}\big(\sqrt{\mathbf{V}_\mu}\,\mathbf{C}^T\,\sqrt{\mathbf{V}_\mu}^T\big)\\
&= \sigma_\mu^{-2}\,\lambda_{\max}\big(\sqrt{\mathbf{V}_\mu}^T\,\sqrt{\mathbf{V}_\mu}\,\mathbf{C}^T\,\sqrt{\mathbf{V}_\mu}^T\,\sqrt{\mathbf{V}_\mu}^{T-1}\big)\\
&= \sigma_\mu^{-2}\,\lambda_{\max}(\mathbf{V}_\mu\,\mathbf{C}^T),\\
\lambda_{\max}(\tilde{\mathbf{D}}) &= \lambda_{\max}\big(\sigma_\mu^{-2}\,\sqrt{\mathbf{V}_\mu}\,\mathbf{D}\,\sqrt{\mathbf{V}_\mu}^T\big)=\sigma_\mu^{-2}\,\lambda_{\max}\big(\sqrt{\mathbf{V}_\mu}^T\,\sqrt{\mathbf{V}_\mu}\,\mathbf{D}\,\sqrt{\mathbf{V}_\mu}^T\,\sqrt{\mathbf{V}_\mu}^{T-1}\big)\\
&= \sigma_\mu^{-2}\,\lambda_{\max}(\mathbf{V}_\mu\,\mathbf{D}),
\end{aligned}$$

$$\tilde{\mathbf{e}}^T[\tilde{\mathbf{D}}-\tilde{\mathbf{C}}]\,\tilde{\mathbf{e}} \;=\; \frac{[\mathbf{W}^{-1}\,\mathbf{e}]^T}{|\mathbf{W}^{-1}\,\mathbf{e}|}\,\mathbf{W}^T[\mathbf{D}-\mathbf{C}]\,\mathbf{W}\,\frac{\mathbf{W}^{-1}\,\mathbf{e}}{|\mathbf{W}^{-1}\,\mathbf{e}|}=\sigma_\mu^{-2}\,\frac{\mathbf{e}^T[\mathbf{D}-\mathbf{C}]\,\mathbf{e}}{\mathbf{e}^T\,\mathbf{V}_\mu^{-1}\,\mathbf{e}}, \tag{S3.93}$$

and

$$\begin{aligned}
\lambda_{\max}(\tilde{\mathbf{C}}_{\mathrm{d}}^T) &= \lambda_{\max}\left(\left[\tilde{\mathbf{C}}+\frac{\tilde{\mathbf{C}}\,\tilde{\mathbf{e}}\,\tilde{\mathbf{e}}^T\,\tilde{\mathbf{C}}}{\tilde{\mathbf{e}}^T[\tilde{\mathbf{D}}-\tilde{\mathbf{C}}]\,\tilde{\mathbf{e}}}\right]^T\right)=\lambda_{\max}\left(\tilde{\mathbf{C}}^T+\frac{\tilde{\mathbf{C}}^T\,\tilde{\mathbf{e}}\,\tilde{\mathbf{e}}^T\,\tilde{\mathbf{C}}^T}{\tilde{\mathbf{e}}^T[\tilde{\mathbf{D}}-\tilde{\mathbf{C}}]\,\tilde{\mathbf{e}}}\right)\\
&= \lambda_{\max}\left(\mathbf{W}^T\,\mathbf{C}^T\,\mathbf{W}+\frac{\mathbf{W}^T\,\mathbf{C}^T\,\mathbf{W}[\mathbf{W}^{-1}\,\mathbf{e}][\mathbf{W}^{-1}\,\mathbf{e}]^T\,\mathbf{W}^T\,\mathbf{C}^T\,\mathbf{W}}{[\mathbf{W}^{-1}\,\mathbf{e}]^T\,\mathbf{W}^T[\mathbf{D}-\mathbf{C}]\,\mathbf{W}\,\mathbf{W}^{-1}\,\mathbf{e}}\right)\\
&= \lambda_{\max}\left(\mathbf{W}^T\left[\mathbf{C}^T+\frac{\mathbf{C}^T\,\mathbf{e}\,\mathbf{e}^T\,\mathbf{C}^T}{\mathbf{e}^T[\mathbf{D}-\mathbf{C}]\,\mathbf{e}}\right]\mathbf{W}\right)\\
&= \lambda_{\max}\left(\mathbf{W}\,\mathbf{W}^T\left[\mathbf{C}^T+\frac{\mathbf{C}^T\,\mathbf{e}\,\mathbf{e}^T\,\mathbf{C}^T}{\mathbf{e}^T[\mathbf{D}-\mathbf{C}]\,\mathbf{e}}\right]\mathbf{W}\,\mathbf{W}^{-1}\right)\\
&= \lambda_{\max}\left(\sigma_\mu^{-2}[\sigma_\mu^2\,\mathbf{W}\,\mathbf{W}^T]\left[\mathbf{C}^T+\frac{\mathbf{C}^T\,\mathbf{e}\,\mathbf{e}^T\,\mathbf{C}^T}{\mathbf{e}^T[\mathbf{D}-\mathbf{C}]\,\mathbf{e}}\right]\right)\\
&= \sigma_\mu^{-2}\,\lambda_{\max}\left(\mathbf{V}_\mu\left[\mathbf{C}+\frac{\mathbf{C}\,\mathbf{e}\,\mathbf{e}^T\,\mathbf{C}}{\mathbf{e}^T[\mathbf{D}-\mathbf{C}]\,\mathbf{e}}\right]^T\right)\\
&= \sigma_\mu^{-2}\,\lambda_{\max}(\mathbf{V}_\mu\,\mathbf{C}_{\mathrm{d}}^T).
\end{aligned} \tag{S3.94}$$

Finally, because the branching possibility conditions are given by substituting $\mathbf{C}=\tilde{\mathbf{C}}$, $\mathbf{D}=\tilde{\mathbf{D}}$, and $\mathbf{e}=\tilde{\mathbf{e}}$ into equations (S3.3), (S3.4), (S3.6), and (S3.7), as

$$\lambda_{\max}(\tilde{\mathbf{C}}) \;<\; 0 \qquad \text{(convergence stability)}, \tag{S3.95}$$

$$\lambda_{\max}(\tilde{\mathbf{D}}) \;>\; 0 \qquad \text{(evolutionary instability)}, \tag{S3.96}$$

$$\tilde{\mathbf{e}}^T[\tilde{\mathbf{D}}-\mathbf{C}]\,\tilde{\mathbf{e}} \;>\; 0 \qquad \text{(dimorphic emergence: mutual invasibility)}, \tag{S3.97}$$

$$\lambda_{\max}(\tilde{\mathbf{C}}_{\mathrm{d}}^T) \;<\; \lambda_{\max}(\tilde{\mathbf{D}}) \qquad \text{(locally stable dimorphic divergence)}, \tag{S3.98}$$

we substitute equations (S3.92)-(S3.94) into equations (S3.95-S3.98) to obtain

$$\lambda_{\max}(\mathbf{C}\,\mathbf{V}_\mu) \;<\; 0 \qquad \text{(convergence stability)}, \tag{S3.99}$$

$$\lambda_{\max}(\mathbf{D}) \;>\; 0 \qquad \text{(evolutionary instability)}, \tag{S3.100}$$

$$\mathbf{e}^T[\mathbf{D}-\mathbf{C}]\,\mathbf{e} \;>\; 0 \qquad \text{(dimorphic emergence: mutual invasibility)}, \tag{S3.101}$$

$$\lambda_{\max}(\mathbf{V}_\mu\,\mathbf{C}_{\mathrm{d}}^T) \;<\; \lambda_{\max}(\mathbf{V}_\mu\,\mathbf{D}) \qquad \text{(locally stable dimorphic divergence)}, \tag{S3.102}$$

with $\mathbf{e}=\mathbf{v}_{\max}(\mathbf{V}_\mu\,\mathbf{D})$ and $\mathbf{C}_{\rm d}=\mathbf{C}+\frac{\mathbf{C}\,\mathbf{e}\,\mathbf{e}^T\,\mathbf{C}}{\mathbf{e}^T[\mathbf{D}-\mathbf{C}]\,\mathbf{e}}$. Note that equation (S3.102) implies that the speed of evolutionary divergence is faster than the speed of deviation of the intermediate phenotype of the dimorphic residents, because substitution of equations (S3.79), (S3.86), (S3.94) into equation (S3.38) gives

$$\begin{aligned}
\frac{d\Delta\,\mathbf{s}}{\rm dt} &= \alpha\mathbf{W}\,\tilde{\mathbf{D}}\,\mathbf{W}^{-1}\,\Delta\,\mathbf{s}=\frac{\mu\sigma_\mu^2[\hat{n}_1+\hat{n}_2]}{4}\mathbf{W}\mathbf{W}^T\,\mathbf{D}\,\Delta\,\mathbf{s}\\
&= \frac{\mu[\hat{n}_1+\hat{n}_2]}{4}\mathbf{V}_\mu\mathbf{D}\,\Delta\,\mathbf{s},\\
\frac{d\breve{\mathbf{s}}}{\rm dt} &= \alpha\mathbf{W}\tilde{\mathbf{C}}_{\rm d}^T\,\mathbf{W}^{-1}\,\breve{\mathbf{s}}=\alpha\mathbf{W}\left\{\mathbf{W}^T\left[\mathbf{C}^T+\frac{\mathbf{C}^T\,\mathbf{e}\,\mathbf{e}^T\,\mathbf{C}^T}{\mathbf{e}^T[\mathbf{D}-\mathbf{C}]\,\mathbf{e}}\right]\mathbf{W}\right\}\mathbf{W}^{-1}\,\breve{\mathbf{s}}\\
&= \alpha\mathbf{W}\mathbf{W}^T\left[\mathbf{C}^T+\frac{\mathbf{C}^T\,\mathbf{e}\,\mathbf{e}^T\,\mathbf{C}^T}{\mathbf{e}^T[\mathbf{D}-\mathbf{C}]\,\mathbf{e}}\right]\breve{\mathbf{s}}\\
&= \frac{\mu[\hat{n}_1+\hat{n}_2]}{4}\mathbf{V}_\mu\mathbf{C}_{\rm d}^T\,\breve{\mathbf{s}} \qquad \text{(S3.103)}
\end{aligned}$$

with $\Delta\,\mathbf{s}=\mathbf{s}_1-\mathbf{s}_2$, $\breve{\mathbf{s}}=\frac{1}{2}[\mathbf{s}_1+\mathbf{s}_2]$, and $\alpha=\frac{\mu\sigma_\mu^2[\hat{n}_1+\hat{n}_2]}{4}$.

Hence, equation (S3.35) implies that the growth of $\Delta\,\mathbf{s}$ (i.e., the speed of evolutionary divergence) is faster than that of $\breve{\mathbf{s}}$ (i.e., the speed of deviation of the intermediate phenotype $\breve{\mathbf{s}}$ from the singular point).

### S4. Branching inevitability condition

#### S4.1. Derivation of equation (18)

Here we show that equation (18) in the main text is derived from equations (14), (16), and (17). Because $\sum_{i=1}^M p_i=1$ gives $\sum_{i=1}^M\frac{dp_i}{dt}=0$, we can transform $d\bar{F}/dt$ as

$$\frac{d\bar{F}}{dt} = \frac{d}{dt}\left\{\sum_{i=1}^M p_i\,F_i\right\}=\sum_{i=1}^M\frac{dp_i}{dt}F_i+\sum_{i=1}^M p_i\frac{dF_i}{dt} \qquad \text{(S4.1)}$$

with $F_i=\mathbf{g}^T[\mathbf{s}_i-\bar{\mathbf{s}}]+[\bar{\mathbf{s}}-\mathbf{s}^*]^T\,\mathbf{C}[\mathbf{s}_i-\bar{\mathbf{s}}]+\frac{1}{2}[\mathbf{s}_i-\bar{\mathbf{s}}]^T\,\mathbf{D}[\mathbf{s}_i-\bar{\mathbf{s}}]-\frac{V_{\mathbf{D}}}{2}+\bar{F}$ (i.e., equation (S2.8), which is equal to equation (14) under $\mathbf{g}=\mathbf{0}$). Then we transform the first and second terms on the right hand side of equation (S4.1) as

$$\begin{aligned}
\sum_{i=1}^M\frac{dp_i}{dt}F_i &= \sum_{i=1}^M\frac{dp_i}{dt}[F_i-\bar{F}]=\sum_{i=1}^M p_i\frac{d\ln p_i}{dt}[F_i-\bar{F}]=\sum_{i=1}^M p_i\left[\frac{d\ln n_i}{dt}-\frac{d\ln\sum_{j=1}^M n_j}{dt}\right][F_i-\bar{F}]\\
&= \sum_{i=1}^M p_i[F_i-\bar{F}]^2,\\
\sum_{i=1}^M p_i\frac{dF_i}{\rm dt} &= \sum_{i=1}^M p_i\frac{d}{\rm dt}\left\{\mathbf{g}^T[\mathbf{s}_i-\bar{\mathbf{s}}]+[\bar{\mathbf{s}}-\mathbf{s}^*]^T\,\mathbf{C}[\mathbf{s}_i-\bar{\mathbf{s}}]+\frac{1}{2}[\mathbf{s}_i-\bar{\mathbf{s}}]^T\,\mathbf{D}[\mathbf{s}_i-\bar{\mathbf{s}}]-\frac{V_{\mathbf{D}}}{2}+\bar{F}\right\}\\
&= -\sum_{i=1}^M p_i\,\mathbf{g}^T\frac{d\bar{\mathbf{s}}}{dt}+\sum_{i=1}^M p_i\left[\frac{d\bar{\mathbf{s}}}{dt}\right]^T\mathbf{C}[\mathbf{s}_i-\bar{\mathbf{s}}]-\sum_{i=1}^M p_i[\bar{\mathbf{s}}-\mathbf{s}^*]^T\,\mathbf{C}\frac{d\bar{\mathbf{s}}}{dt}\\
&\quad -\sum_{i=1}^M p_i[\mathbf{s}_i-\bar{\mathbf{s}}]^T\,\mathbf{D}\frac{d\bar{\mathbf{s}}}{\rm dt}-\frac{1}{2}\frac{dV_{\mathbf{D}}}{dt}+\frac{d\bar{F}}{dt}\\
&= -\mathbf{g}^T\frac{d\bar{\mathbf{s}}}{dt}-[\bar{\mathbf{s}}-\mathbf{s}^*]^T\,\mathbf{C}\frac{d\bar{\mathbf{s}}}{dt}-\frac{1}{2}\frac{dV_{\mathbf{D}}}{dt}+\frac{d\bar{F}}{dt}
\end{aligned}$$

$$
\begin{aligned}
&= -\mathbf{g}^T \frac{d\,\bar{\mathbf{s}}}{dt} - \frac{1}{2}\left\{[\bar{\mathbf{s}}-\mathbf{s}^*]^T \mathbf{C}\frac{d\,\bar{\mathbf{s}}}{dt} + \frac{d\,\bar{\mathbf{s}}^T}{dt}\mathbf{C}[\bar{\mathbf{s}}-\mathbf{s}^*]\right\} - \frac{1}{2}\left\{[\bar{\mathbf{s}}-\mathbf{s}^*]^T \mathbf{C}\frac{d\,\bar{\mathbf{s}}}{dt} - \frac{d\,\bar{\mathbf{s}}^T}{dt}\mathbf{C}[\bar{\mathbf{s}}-\mathbf{s}^*]\right\} \\
&\quad -\frac{1}{2}\frac{dV_{\mathbf{D}}}{dt} + \frac{d\bar{F}}{dt} \\
&= -\mathbf{g}^T \frac{d\,\bar{\mathbf{s}}}{dt} - \frac{1}{2}\frac{d[\bar{\mathbf{s}}-\mathbf{s}^*]^T\mathbf{C}[\bar{\mathbf{s}}-\mathbf{s}^*]}{dt} - \frac{1}{2}\left\{[\bar{\mathbf{s}}-\mathbf{s}^*]^T \mathbf{C}\frac{d\,\bar{\mathbf{s}}}{\mathrm{dt}} - [\bar{\mathbf{s}}-\mathbf{s}^*]^T \mathbf{C}^T\frac{d\,\bar{\mathbf{s}}}{\mathrm{dt}}\right\} - \frac{1}{2}\frac{dV_{\mathbf{D}}}{dt} + \\
&\quad \frac{d\bar{F}}{dt} \\
&= -\mathbf{g}^T \frac{d\,\bar{\mathbf{s}}}{dt} - \frac{d\{\frac{1}{2}[\bar{\mathbf{s}}-\mathbf{s}^*]^T\mathbf{C}[\bar{\mathbf{s}}-\mathbf{s}^*] + \frac{1}{2}V_{\mathbf{D}}\}}{dt} + \frac{1}{2}[\bar{\mathbf{s}}-\mathbf{s}^*]^T[\mathbf{C}^T-\mathbf{C}]\frac{d\,\bar{\mathbf{s}}}{dt} + \frac{d\bar{F}}{dt},
\end{aligned}
\tag{S4.2}
$$

which upon substitution into equation (S4.1) gives

$$
\frac{dH}{dt} = \sum_{i=1}^{M} p_i[F_i-\bar{F}]^2 - \mathbf{g}^T\frac{d\,\bar{\mathbf{s}}}{dt} + \frac{1}{2}[\bar{\mathbf{s}}-\mathbf{s}^*]^T[\mathbf{C}^T-\mathbf{C}]\frac{d\,\bar{\mathbf{s}}}{dt}, \tag{S4.3}
$$

$$
H = \frac{1}{2}[\bar{\mathbf{s}}-\mathbf{s}^*]^T\mathbf{C}[\bar{\mathbf{s}}-\mathbf{s}^*] + \frac{1}{2}V_{\mathbf{D}}. \tag{S4.4}
$$

Hence, provided that $\mathbf{s}^*$ satisfies the branching inevitability conditions, i.e., $\mathbf{g}=\mathbf{0}$, $\mathbf{C}^T-\mathbf{C}=0$, $\lambda_{\max}(\mathbf{C})<0$, and $\lambda_{\max}(\mathbf{D})>0$, we see that $\frac{dH}{\mathrm{dt}}>0$ always holds unless the population dynamics completely stops.

Note that equations (S4.2) and (S4.3) can be combined into

$$
\sum_{i=1}^{M}\frac{dp_i}{dt}F_i = \sum_{i=1}^{M} p_i[F_i-\bar{F}]^2 \tag{S4.5}
$$

$$
= \frac{dH}{dt}. \tag{S4.6}
$$

Here, $\sum_{i=1}^{M}\frac{dp_i}{dt}F_i$ describes the increase of the mean fitness $\bar{F}$ of existing phenotypes due to selection (i.e., population dynamics), which is equal to the variance of their fitnesses. Hence, equation (S4.5) represents the Fisher's fundamental theorem of natural selection (Fisher, 1930; Baez, 2021). Therefore, equation (S4.6) means that equation (18) in the main text (i.e., $\frac{dH}{dt} = \sum_{i=1}^{M}p_i[F_i-\bar{F}]^2$) is a locally approximated representation of the fundamental theorem, in the sense that the fitness function is approximated as a parabolic function of the mutant phenotype (equation (14) in the main text) provided that all existing phenotypes are close to the singular point, $\mathbf{s}^*$.

#### S4.2. Mutant invasibility under an arbitrary set of resident phenotypes

Here we derive that there always exist mutants having positive invasion fitness under an arbitrary set of resident phenotypes denoted by $\mathbf{s}_1, ..., \mathbf{s}_M$ at population-dynamical equilibrium, provided that the branching inevitability conditions (i.e., $\mathbf{C}^T-\mathbf{C}=0$, $\lambda_{\max}(\mathbf{C})<0$, and $\lambda_{\max}(\mathbf{D})>0$) are satisfied. The invasion fitness for a mutant $\mathbf{s}'$ against the residents is derived from equation (14) in the main text as

$$
\begin{aligned}
f(\mathbf{s}';\mathbf{s}_1,...,\mathbf{s}_M) &= \lim_{n'\to 0} F(\mathbf{s}';(\mathbf{s}_1,...,\mathbf{s}_M,\mathbf{s}'),(\hat{n}_1,...,\hat{n}_M,n')) \\
&= [\bar{\mathbf{s}}-\mathbf{s}^*]^T\mathbf{C}[\mathbf{s}'-\bar{\mathbf{s}}] + \frac{1}{2}[\mathbf{s}'-\bar{\mathbf{s}}]^T\mathbf{D}[\mathbf{s}'-\bar{\mathbf{s}}] - \frac{V_{\mathbf{D}}}{2},
\end{aligned}
\tag{S4.7}
$$

where $\hat{n}_1, ..., \hat{n}_M$ describe the residents' equilibrium population sizes, and where $\bar{\mathbf{s}} = \sum_{j=1}^{M}\hat{n}_j\,\mathbf{s}_j / \sum_{j=1}^{M}\hat{n}_j$ gives their average phenotype at their population-dynamical equilibrium. Since the residents are at population-dynamical equilibrium, $\mathbf{s}_i$ for all $i=1,...M$ must satisfy

$$
f(\mathbf{s}_i;\mathbf{s}_1,...,\mathbf{s}_M) = [\bar{\mathbf{s}}-\mathbf{s}^*]^T\mathbf{C}[\mathbf{s}_i-\bar{\mathbf{s}}] + \frac{1}{2}[\mathbf{s}_i-\bar{\mathbf{s}}]^T\mathbf{D}[\mathbf{s}_i-\bar{\mathbf{s}}] - \frac{V_{\mathbf{D}}}{2} = 0. \tag{S4.8}
$$

We assume without loss of generality that $\mathbf{s}'$ derives from the $i$th resident, $\mathbf{s}_i$, and express the mutant as $\mathbf{s}'^{+} = \mathbf{s}_i + \delta\,\mathbf{s}$. Then by using equation (S4.8), we transform the invasion fitnesses of $\mathbf{s}'^{+} = \mathbf{s}_i + \delta\,\mathbf{s}$ and $\mathbf{s}'^{-} = \mathbf{s}_i - \delta\,\mathbf{s}$ as

$$\begin{aligned} f(\mathbf{s}'^{+};\mathbf{s}_1,...,\mathbf{s}_M) &= [\bar{\mathbf{s}} - \mathbf{s}^*]^T\,\mathbf{C}[\delta\,\mathbf{s} + \mathbf{s}_i - \bar{\mathbf{s}}] + \frac{1}{2}[\delta\,\mathbf{s} + \mathbf{s}_i - \bar{\mathbf{s}}]^T\,\mathbf{D}[\delta\,\mathbf{s} + \mathbf{s}_i - \bar{\mathbf{s}}] - \frac{V_{\mathbf{D}}}{2} \\ &= [\bar{\mathbf{s}} - \mathbf{s}^*]^T\,\mathbf{C}\,\delta\,\mathbf{s} + [\mathbf{s}_i - \bar{\mathbf{s}}]^T\,\mathbf{D}\,\delta\,\mathbf{s} + \frac{1}{2}\delta\,\mathbf{s}^T\,\mathbf{D}\,\delta\,\mathbf{s} \\ &\quad + [\bar{\mathbf{s}} - \mathbf{s}^*]^T\,\mathbf{C}[\mathbf{s}_i - \bar{\mathbf{s}}] + \frac{1}{2}[\mathbf{s}_i - \bar{\mathbf{s}}]^T\,\mathbf{D}[\mathbf{s}_i - \bar{\mathbf{s}}] - \frac{V_{\mathbf{D}}}{2} \\ &= \{[\bar{\mathbf{s}} - \mathbf{s}^*]^T\,\mathbf{C} + [\mathbf{s}_i - \bar{\mathbf{s}}]^T\,\mathbf{D}\}\,\delta\,\mathbf{s} + \frac{1}{2}\delta\,\mathbf{s}^T\,\mathbf{D}\,\delta\,\mathbf{s}, \\ f(\mathbf{s}'^{-};\mathbf{s}_1,...,\mathbf{s}_M) &= -\{[\bar{\mathbf{s}} - \mathbf{s}^*]^T\,\mathbf{C} + [\mathbf{s}_i - \bar{\mathbf{s}}]^T\,\mathbf{D}\}\,\delta\,\mathbf{s} + \frac{1}{2}\delta\,\mathbf{s}^T\,\mathbf{D}\,\delta\,\mathbf{s}, \end{aligned} \tag{S4.9}$$

from which we get

$$f(\mathbf{s}'^{+};\mathbf{s}_1,...,\mathbf{s}_M) + f(\mathbf{s}'^{-};\mathbf{s}_1,...,\mathbf{s}_M) = \delta\,\mathbf{s}^T\,\mathbf{D}\,\delta\,\mathbf{s}. \tag{S4.10}$$

Therefore, as long as $\lambda_{\max}(\mathbf{D}) > 0$ holds, we can find $\delta\,\mathbf{s}$ satisfying $f(\mathbf{s}'^{+};\mathbf{s}_1,...,\mathbf{s}_M) + f(\mathbf{s}'^{-};\mathbf{s}_1,..., \mathbf{s}_M) = \delta\,\mathbf{s}^T\,\mathbf{D}\,\delta\,\mathbf{s} > 0$, in which case either $\mathbf{s}'^{+} = \mathbf{s}_i + \delta\,\mathbf{s}$ or $\mathbf{s}'^{-} = \mathbf{s}_i - \delta\,\mathbf{s}$ has positive invasion fitness.

#### S4.3. Phenotypic variance guaranteed by $H$

We denote the eigenvalues of $\mathbf{D}$ by $\lambda_{\mathbf{D},1}$, ..., $\lambda_{\mathbf{D},L}$. We assume without loss of generality that $1,...,\tilde{L}$th eigenvalues are positive, satisfying $\lambda_{\mathbf{D},1} \geqslant ... \geqslant \lambda_{\mathbf{D},\tilde{L}} > 0 \geqslant \lambda_{\mathbf{D},\tilde{L}+1} \geqslant \cdots \geqslant \lambda_{\mathbf{D},L}$. Then we decompose $\mathbf{D}$ into two matrices $\mathbf{D}^+$ and $\mathbf{D}^-$ which correspond to the positive and non-positive eigenvalues of $\mathbf{D}$, as

$$\begin{aligned} \mathbf{D} &= (\,\mathbf{v}_{\mathbf{D}1}\ \cdots\ \mathbf{v}_{\mathbf{D}L}\,)\begin{pmatrix} \lambda_{\mathbf{D}1} & \cdots & 0 \\ \vdots & \ddots & \vdots \\ 0 & \cdots & \lambda_{\mathbf{D}L} \end{pmatrix}(\,\mathbf{v}_{\mathbf{D}1}\ \cdots\ \mathbf{v}_{\mathbf{D}L}\,)^T = \sum_{l=1}^{L} \lambda_{\mathbf{D}l}\,\mathbf{v}_{\mathbf{D}l}\,\mathbf{v}_{\mathbf{D}l}^T \\ &= \sum_{l=1}^{\tilde{L}} \lambda_{\mathbf{D}l}\,\mathbf{v}_{\mathbf{D}l}\,\mathbf{v}_{\mathbf{D}l}^T + \sum_{l=\tilde{L}+1}^{L} \lambda_{\mathbf{D}l}\,\mathbf{v}_{\mathbf{D}l}\,\mathbf{v}_{\mathbf{D}l}^T \\ &= \mathbf{D}^+ + \mathbf{D}^-, \\ \mathbf{D}^+ &= \sum_{l=1}^{\tilde{L}} \lambda_{\mathbf{D}l}\,\mathbf{v}_{\mathbf{D}l}\,\mathbf{v}_{\mathbf{D}l}^T, \\ \mathbf{D}^- &= \sum_{l=\tilde{L}+1}^{L} \lambda_{\mathbf{D}l}\,\mathbf{v}_{\mathbf{D}l}\,\mathbf{v}_{\mathbf{D}l}^T. \end{aligned} \tag{S4.11}$$

On this basis, we decompose $V_{\mathbf{D}}$ as

$$\begin{aligned} V_{\mathbf{D}} &= \sum_{j=1}^{M} p_j[\mathbf{s}_j - \bar{\mathbf{s}}]^T\,\mathbf{D}[\mathbf{s}_j - \bar{\mathbf{s}}] = \sum_{j=1}^{M} p_j[\mathbf{s}_j - \bar{\mathbf{s}}]^T\,[\mathbf{D}^+ + \mathbf{D}^-][\mathbf{s}_j - \bar{\mathbf{s}}] = V_{\mathbf{D}}^+ + V_{\mathbf{D}}^-, \\ V_{\mathbf{D}}^+ &= \sum_{j=1}^{M} p_j[\mathbf{s}_j - \bar{\mathbf{s}}]^T\,\mathbf{D}^+[\mathbf{s}_j - \bar{\mathbf{s}}], \\ V_{\mathbf{D}}^- &= \sum_{j=1}^{M} p_j[\mathbf{s}_j - \bar{\mathbf{s}}]^T\,\mathbf{D}^-[\mathbf{s}_j - \bar{\mathbf{s}}], \end{aligned} \tag{S4.12}$$

which upon substitution into equation (S4.4) gives

$$V_{\mathbf{D}}^+ = H - [\bar{\mathbf{s}} - \mathbf{s}^*]^T\,\mathbf{C}[\bar{\mathbf{s}} - \mathbf{s}^*] - V_{\mathbf{D}}^-. \tag{S4.13}$$

Note that $V_{\mathbf{D}}^{+}\geqslant 0$, $V_{\mathbf{D}}^{-}\leqslant 0$, and $[\bar{\mathbf{s}}-\mathbf{s}^*]^T\,\mathbf{C}[\bar{\mathbf{s}}-\mathbf{s}^*]\leqslant 0$ always hold under the branching inevitability conditions (i.e., $\mathbf{C}^T-\mathbf{C}=0$, $\lambda_{\max}(\mathbf{C})<0$, and $\lambda_{\max}(\mathbf{D})>0$). Hence we get

$$V_{\mathbf{D}}^{+} \;\geqslant\; H. \tag{S4.14}$$

Therefore, if $H$ has a positive value, then the phenotypic variance of existing phenotypes measured by means of $V_{\mathbf{D}}^{+}$ (i.e., phenotypic variance along the eigenvectors of positive eigenvalues of $\mathbf{D}$) is no smaller than $H$.

#### S4.4. Local lyapunov function for absolutely convergence stable ESS

Here we assume that $\mathbf{s}^*$ is an absolutely convergence stable ESS, satisfying $\mathbf{C}^T-\mathbf{C}=0$, $\lambda_{\max}(\mathbf{C})<0$, and $\lambda_{\max}(\mathbf{D})<0$. In this case, the following inequalities trivially hold,

$$\begin{aligned} H_C &= \frac{1}{2}[\bar{\mathbf{s}}-\mathbf{s}^*]^T\,\mathbf{C}[\bar{\mathbf{s}}-\mathbf{s}^*]\leqslant 0,\\ H_D &= \frac{1}{2}V_{\mathbf{D}}=\frac{1}{2}\sum_{j=1}^{M} p_j[\mathbf{s}_j-\bar{\mathbf{s}}]^T\,\mathbf{D}[\mathbf{s}_j-\bar{\mathbf{s}}]\leqslant 0,\\ H &= H_C+H_D\leqslant 0, \end{aligned} \tag{S4.15}$$

where $H=0$ (i.e., $H_C=H_D=0$) holds only when the system has a single phenotype equal to $\mathbf{s}^*$. Note that equation (18) in the main text,

$$\frac{dH}{dt} \;=\; \sum_{i=1}^{M} p_i[F_i-\bar{F}]^2\geqslant 0, \tag{S4.16}$$

is still satisfied, because its derivation shown in Appendix S4.1 is independent of the signs of $\lambda_{\max}(\mathbf{C})$ and $\lambda_{\max}(\mathbf{D})$.

Under a single resident phenotype $\mathbf{s}^*$ having its equilibrium population size $\hat{n}^*$, the invasion fitness for a mutant $\mathbf{s}'$ is given by

$$\begin{aligned} f(\mathbf{s}';\mathbf{s}^*) &= \lim_{n'\to 0} F(\mathbf{s}';(\mathbf{s}',\mathbf{s}^*),(n',\hat{n}^*))\\ &= \frac{1}{2}[\mathbf{s}'-\mathbf{s}^*]^T\,\mathbf{D}[\mathbf{s}'-\mathbf{s}^*]. \end{aligned} \tag{S4.17}$$

Hence $f(\mathbf{s}';\mathbf{s}^*)<0$ holds for any $\mathbf{s}'\neq\mathbf{s}^*$ under $\lambda_{\max}(\mathbf{D})<0$, in which case no mutant can invade the resident $\mathbf{s}^*$. Clearly, when there exists a resident having phenotype $\mathbf{s}^*$, no other phenotype can coexist with it. When the system has $M$ coexisting residents (with $M\geqslant 1$) that differ from $\mathbf{s}^*$, denoted by $\mathbf{s}_1,...,\mathbf{s}_M$, we see from equation (S4.9) that the invasion fitness for a mutant deriving from the $i$th resident, $\mathbf{s}_i$, can be expressed as

$$\begin{aligned} f(\mathbf{s}';\mathbf{s}_1,...,\mathbf{s}_M) &= \mathbf{g}(\mathbf{s}_i;\mathbf{s}_1,...,\mathbf{s}_M)^T[\mathbf{s}'-\mathbf{s}_i]+\frac{1}{2}[\mathbf{s}'-\mathbf{s}_i]^T\,\mathbf{D}[\mathbf{s}'-\mathbf{s}_i],\\ \mathbf{g}(\mathbf{s}_i;\mathbf{s}_1,...,\mathbf{s}_M)^T &= [\bar{\mathbf{s}}-\mathbf{s}^*]^T\,\mathbf{C}+[\mathbf{s}_i-\bar{\mathbf{s}}]^T\,\mathbf{D} \end{aligned} \tag{S4.18}$$

Then we can find mutants with positive invasion fitnesses by choosing $\mathbf{s}'=\mathbf{s}_i+\frac{\mathbf{g}(\mathbf{s}_i;\,\mathbf{s}_1,\,...,\,\mathbf{s}_M)}{|\mathbf{g}(\mathbf{s}_i;\,\mathbf{s}_1,\,...,\,\mathbf{s}_M)|}\,\sigma$ with sufficiently small and positive $\sigma$ so that

$$f(\mathbf{s}';\mathbf{s}_1,...,\mathbf{s}_M) \;=\; |\mathbf{g}(\mathbf{s}_i;\mathbf{s}_1,...,\mathbf{s}_M)|\,\sigma+\frac{1}{2}\frac{\mathbf{g}(\mathbf{s}_i;\mathbf{s}_1,...,\mathbf{s}_M)^T\,\mathbf{D}\,\mathbf{g}(\mathbf{s}_i;\mathbf{s}_1,...,\mathbf{s}_M)}{|\mathbf{g}(\mathbf{s}_i;\mathbf{s}_1,...,\mathbf{s}_M)|^2}\,\sigma^2>0$$

holds. Hence, unless the system has reached the state of a monomorphic resident phenotype equal to $\mathbf{s}^*$ (i.e., $H=0$), mutants can invade the system. Therefore, for an arbitrary set of initial resident phenotypes corresponding to $H<0$, mutant invasion is repeated so that each invasion monotonically increases $H$, until the system has reached the state of a monomorphic resident phenotype equal to $\mathbf{s}^*$ (i.e., $H=0$), provided that changes of $H$ caused directly by the emergence of mutants are negligibly small.

#### S4.5. Relationship with potential game in evolutionary game theory

We see from equation (14) in the main text that the fraction dynamics is given by

$$\frac{d\ln p_i}{dt} = \frac{d\ln n_i}{dt} - \frac{d\ln\sum_{j=1}^{M} n_j}{dt} = F_i - \bar{F} = \bar{\mathbf{s}}^T\mathbf{C}[\mathbf{s}_i - \bar{\mathbf{s}}] + \frac{1}{2}[\mathbf{s}_i - \bar{\mathbf{s}}]^T\mathbf{D}[\mathbf{s}_i - \bar{\mathbf{s}}] - \frac{1}{2}V_{\mathbf{D}}. \quad \text{(S4.19)}$$

This equation can also be expressed as

$$\begin{aligned} \frac{d\ln p_i}{dt} &= U_i - \bar{U}, \\ U_i &= \frac{\partial H}{\partial p_i}, \\ \bar{U} &= \sum_{j=1}^{M} p_j U_j \end{aligned} \quad \text{(S4.20)}$$

with $H = \frac{1}{2}\bar{\mathbf{s}}^T\mathbf{C}\,\bar{\mathbf{s}} + \frac{1}{2}V_{\mathbf{D}}$ (equation (17) in the main text), because we see that

$$\begin{aligned} U_i &= \frac{\partial H}{\partial p_i} = \frac{1}{2}\frac{\partial\bar{\mathbf{s}}^T}{\partial p_i}\mathbf{C}\,\bar{\mathbf{s}} + \frac{1}{2}\bar{\mathbf{s}}^T\mathbf{C}\frac{\partial\bar{\mathbf{s}}}{\partial p_i} + \frac{1}{2}\frac{\partial V_{\mathbf{D}}}{\partial p_i} \\ &= \bar{\mathbf{s}}^T\mathbf{C}\frac{\partial\sum_{j=1}^{M} p_j\mathbf{s}_j}{\partial p_i} + \frac{1}{2}\frac{\partial\sum_{j=1}^{M} p_j[\mathbf{s}_j - \bar{\mathbf{s}}]^T\mathbf{D}[\mathbf{s}_j - \bar{\mathbf{s}}]}{\partial p_i} \\ &= \bar{\mathbf{s}}^T\mathbf{C}\,\mathbf{s}_i + \frac{1}{2}[\mathbf{s}_i - \bar{\mathbf{s}}]^T\mathbf{D}[\mathbf{s}_i - \bar{\mathbf{s}}] + \frac{1}{2}\sum_{j=1}^{M} p_j\frac{\partial\,[\mathbf{s}_j - \bar{\mathbf{s}}]^T}{\partial p_i}\mathbf{D}[\mathbf{s}_j - \bar{\mathbf{s}}] + \frac{1}{2}\sum_{j=1}^{M} p_j[\mathbf{s}_j - \bar{\mathbf{s}}]^T\mathbf{D}\frac{\partial\,[\mathbf{s}_j - \bar{\mathbf{s}}]}{\partial p_i} \\ &= \bar{\mathbf{s}}^T\mathbf{C}\,\mathbf{s}_i + \frac{1}{2}[\mathbf{s}_i - \bar{\mathbf{s}}]^T\mathbf{D}[\mathbf{s}_i - \bar{\mathbf{s}}] - \frac{1}{2}\sum_{j=1}^{M} p_j\,\mathbf{s}_i^T\mathbf{D}[\mathbf{s}_j - \bar{\mathbf{s}}] - \frac{1}{2}\sum_{j=1}^{M} p_j[\mathbf{s}_j - \bar{\mathbf{s}}]^T\mathbf{D}\,\mathbf{s}_i \\ &= \bar{\mathbf{s}}^T\mathbf{C}\,\mathbf{s}_i + \frac{1}{2}[\mathbf{s}_i - \bar{\mathbf{s}}]^T\mathbf{D}[\mathbf{s}_i - \bar{\mathbf{s}}] \\ &= \left\{\bar{\mathbf{s}}^T\mathbf{C}[\mathbf{s}_i - \bar{\mathbf{s}}] + \frac{1}{2}[\mathbf{s}_i - \bar{\mathbf{s}}]^T\mathbf{D}[\mathbf{s}_i - \bar{\mathbf{s}}] - \frac{1}{2}V_{\mathbf{D}}\right\} + \bar{\mathbf{s}}^T\mathbf{C}\,\bar{\mathbf{s}} + \frac{1}{2}V_{\mathbf{D}} \\ &= F_i - \bar{F} + \bar{\mathbf{s}}^T\mathbf{C}\,\bar{\mathbf{s}} + \frac{1}{2}V_{\mathbf{D}}, \\ \bar{U} &= \sum_{j=1}^{M} p_j U_j = \bar{\mathbf{s}}^T\mathbf{C}\,\bar{\mathbf{s}} + \frac{1}{2}V_{\mathbf{D}}, \end{aligned} \quad \text{(S4.21)}$$

and hence that $U_i - \bar{U} = F_i - \bar{F}$ holds, where $\partial\bar{\mathbf{s}} / \partial p_i$ is calculated by treating $p_1$, ..., $p_M$ as independent variables (Sandholm, 2001).

In evolutionary game theory, the dynamics in the form of equation (S4.20) is known as the replicator dynamics under the potential game, ensuring that the time derivative of the potential, $H$, satisfies

$$\frac{dH}{dt} = \sum_{i=1}^{M} p_i[U_i - \bar{U}]^2, \quad \text{(S4.22)}$$

(equation 19.18 in Hofbauer and Sigmund, 1998), which corresponds to Fisher's fundamental theorem of natural selection (Sandholm, 2001). Because $U_i - \bar{U} = F_i - \bar{F}$ holds, equation (S4.19) is equivalent to equation (18) in the main text,

$$\frac{dH}{dt} = \sum_{j=1}^{M} p_j[F_j - \bar{F}]^2. \quad \text{(S4.23)}$$

### S5. Branching possibility for three classes of convergence stable points

In this section, for the three classes of convergence stable points, we examine the branching possibility conditions, equation (15) in the main text. Without loss of generality, we assume that the trait space has been normalized so that the mutational covariance matrix is given by $\mathbf{V}_\mu = \sigma_\mu^2\,\mathbf{I}$. Then the branching possibility conditions, equation (15), are simplified into

$$\begin{aligned}
\lambda_{\max}(\mathbf{C}) &< 0 && \text{(convergence stability)}, && \text{(S5.1)}\\
\lambda_{\max}(\mathbf{D}) &> 0 && \text{(evolutionary instability)}, && \text{(S5.2)}\\
\mathbf{e}^T[\mathbf{D}-\mathbf{C}]\,\mathbf{e} &> 0 && \text{(dimorphic emergence: mutual invasibility)}, && \text{(S5.3)}\\
\lambda_{\max}(\mathbf{C}_\mathrm{d}) &< \lambda_{\max}(\mathbf{D}) && \text{(locally stable dimorhpic divergence)}, && \text{(S5.4)}\\
\mathbf{C}_\mathrm{d} &= \mathbf{C} + \frac{\mathbf{C}\,\mathbf{e}\,\mathbf{e}^T\,\mathbf{C}}{\mathbf{e}^T[\mathbf{D}-\mathbf{C}]\,\mathbf{e}}, \\
\mathbf{e} &= \mathbf{v}_{\lambda\max}(\mathbf{D}).
\end{aligned}$$

The conditions for an evolutionarily singular point $\mathbf{s}^*$ being strong convergence stable, and absolute convergence stable, are respectively given by

$$\lambda_{\max}\left(\frac{\mathbf{C}+\mathbf{C}^T}{2}\right) < 0 \quad \text{(strong convergence)}, \qquad \text{(S5.5)}$$

$$\mathbf{C} = \mathbf{C}^T,\ \lambda_{\max}(\mathbf{C}) < 0 \quad \text{(absolute convergence)}. \qquad \text{(S5.6)}$$

#### S5.1. Convergence stable non-ESSes in two-dimensional trait spaces

Here we show that a convergence stable non-ESS having mutual invasibility (i.e., equations (S5.1), (S5.2), and (S5.3)) in two-dimensional trait spaces always satisfies the condition for locally stable dimorphic divergence (i.e., equation (S5.4)). In an arbitrary two-dimensional trait space $\mathbf{s} = (x, y)^T$, we consider a convergence stable non-ESS that also satisfies the condition for mutual invasibility (i.e., equations (S5.1), (S5.2), and (S5.3)). Without loss of generality, we assume that the trait space has been appropriately rotated so that $\mathbf{e} = (1, 0)^T$ holds. Then we transform $\mathbf{C}_\mathrm{d}$ in equation (S4.4) as

$$\begin{aligned}
\mathbf{C}_\mathrm{d} &= \mathbf{C} + \frac{\mathbf{C}\,\mathbf{e}\,\mathbf{e}^T\,\mathbf{C}}{\mathbf{e}^T[\mathbf{D}-\mathbf{C}]\,\mathbf{e}} \\
&= \begin{pmatrix} C_\mathrm{xx} & C_\mathrm{xy} \\ C_\mathrm{yx} & C_\mathrm{yy} \end{pmatrix} + \frac{\begin{pmatrix} C_\mathrm{xx} & C_\mathrm{xy} \\ C_\mathrm{yx} & C_\mathrm{yy} \end{pmatrix}\begin{pmatrix} 1 \\ 0 \end{pmatrix}\begin{pmatrix} 1 & 0 \end{pmatrix}\begin{pmatrix} C_\mathrm{xx} & C_\mathrm{xy} \\ C_\mathrm{yx} & C_\mathrm{yy} \end{pmatrix}}{\begin{pmatrix} 1 & 0 \end{pmatrix}\begin{pmatrix} C_\mathrm{xx} & C_\mathrm{xy} \\ C_\mathrm{yx} & C_\mathrm{yy} \end{pmatrix}\begin{pmatrix} 1 \\ 0 \end{pmatrix}} \\
&= \begin{pmatrix} C_\mathrm{xx} & C_\mathrm{xy} \\ C_\mathrm{yx} & C_\mathrm{yy} \end{pmatrix} + \frac{\begin{pmatrix} C_\mathrm{xx} \\ C_\mathrm{yx} \end{pmatrix}\begin{pmatrix} C_\mathrm{xx} & C_\mathrm{xy} \end{pmatrix}}{[D_\mathrm{xx} - C_\mathrm{xx}]} \\
&= \begin{pmatrix} C_\mathrm{xx} & C_\mathrm{xy} \\ C_\mathrm{yx} & C_\mathrm{yy} \end{pmatrix} + \frac{\begin{pmatrix} C_\mathrm{xx}^2 & C_\mathrm{xx}C_\mathrm{xy} \\ C_\mathrm{xx}C_\mathrm{yx} & C_\mathrm{xy}C_\mathrm{yx} \end{pmatrix}}{[D_\mathrm{xx} - C_\mathrm{xx}]} \\
&= \frac{D_\mathrm{xx}}{D_\mathrm{xx} - C_\mathrm{xx}}\begin{pmatrix} C_\mathrm{xx} & C_\mathrm{xy} \\ C_\mathrm{yx} & C_\mathrm{yy} - \frac{C_\mathrm{xx}C_\mathrm{yy} - C_\mathrm{xy}C_\mathrm{yx}}{D_\mathrm{xx}} \end{pmatrix},
\end{aligned} \qquad \text{(S5.7)}$$

where $D_\mathrm{xx} = \lambda_{\max}(\mathbf{D}) > 0$ and $D_\mathrm{xx} - C_\mathrm{xx} = \mathbf{e}^T[\mathbf{D}-\mathbf{C}]\,\mathbf{e} > 0$ hold under equations (S5.2) and (S5.3).

Because $\lambda_{\max}(\mathbf{C}) < 0$ holds, the two eigenvalues of $\mathbf{C}$, denoted by $\lambda_{C1}$ and $\lambda_{C2}$, both have negative real parts, satisfying

$$\begin{aligned} \lambda_{C1} + \lambda_{C2} &= C_{\rm xx} + C_{\rm yy} < 0, \\ \lambda_{C1}\, \lambda_{C2} &= C_{\rm xx}\, C_{\rm yy} - C_{\rm xy}\, C_{\rm yx} > 0. \end{aligned} \tag{S5.8}$$

In this case, the eigenvalues of $\mathbf{C}_{\rm d}$ in equation (S5.7), denoted by $\breve{\lambda}_{C1}$ and $\breve{\lambda}_{C2}$, have negative real parts as well, because we see that

$$\begin{aligned} \breve{\lambda}_{C1} + \breve{\lambda}_{C2} &= \frac{D_{\rm xx}}{D_{\rm xx} - C_{\rm xx}} \left[ C_{\rm xx} + C_{\rm yy} - \frac{C_{\rm xx}\, C_{\rm yy} - C_{\rm xy}\, C_{\rm yx}}{D_{\rm xx}} \right] < \frac{D_{\rm xx}[C_{\rm xx} + C_{\rm yy}]}{D_{\rm xx} - C_{\rm xx}} < 0, \\ \breve{\lambda}_{C1}\, \breve{\lambda}_{C2} &= \frac{D_{\rm xx}^2}{[D_{\rm xx} - C_{\rm xx}]^2} \left\{ C_{\rm xx} \left[ C_{\rm yy} - \frac{C_{\rm xx}\, C_{\rm yy} - C_{\rm xy}\, C_{\rm yx}}{D_{\rm xx}} \right] - C_{\rm xy}\, C_{\rm yx} \right\} \\ &= \frac{D_{\rm xx}}{[D_{\rm xx} - C_{\rm xx}]} [C_{\rm xx}\, C_{\rm yy} - C_{\rm xy}\, C_{\rm yx}] > 0. \end{aligned} \tag{S5.9}$$

Therefore, in a two-dimensional trait space, a convergence stable non-ESS having mutual invasibility (i.e., equations (S5.1), (S5.2), and (S5.3) always satisfies the condition for locally stable dimorphic divergence, equation (S5.4).

Regarding three-dimensional trait space, the condition for dimorphic emergence is not sufficient for locally stable dimorphic divergence. For example,

$$\mathbf{C} = \begin{pmatrix} 0 & 6 & 0 \\ 5 & 0 & 5 \\ -14 & -14 & -11 \end{pmatrix}, \quad \mathbf{D} = \begin{pmatrix} 2 & 0 & 0 \\ 0 & -\frac{1}{2} & 0 \\ 0 & 0 & -\frac{1}{2} \end{pmatrix} \tag{S5.10}$$

give

$$\lambda_{\max}(\mathbf{C}) = \frac{[566 - 9\sqrt{3955}]^{1/3} + [566 - 9\sqrt{3955}]^{-1/3} - 22}{6} \simeq -1.913684 < 0, \tag{S5.11}$$

$$\lambda_{\max}(\mathbf{D}) = 2 > 0, \tag{S5.12}$$

$$\mathbf{e} = \begin{pmatrix} 1 \\ 0 \\ 0 \end{pmatrix}, \tag{S5.13}$$

$$\mathbf{e}^T [\mathbf{D} - \mathbf{C}]\, \mathbf{e} = 2 > 0, \tag{S5.14}$$

and

$$\begin{aligned} \mathbf{C}_{\rm d} &= \mathbf{C} + \frac{\mathbf{C}\, \mathbf{e}\, \mathbf{e}^T\, \mathbf{C}}{\mathbf{e}^T [\mathbf{D} - \mathbf{C}]\, \mathbf{e}}, \qquad \text{(S5.15)} \\ &= \mathbf{C} + \frac{\begin{pmatrix} 0 & 6 & 0 \\ 5 & 0 & 5 \\ -14 & -14 & -11 \end{pmatrix} \begin{pmatrix} 1 \\ 0 \\ 0 \end{pmatrix} \begin{pmatrix} 1 & 0 & 0 \end{pmatrix} \begin{pmatrix} 0 & 6 & 0 \\ 5 & 0 & 5 \\ -14 & -14 & -11 \end{pmatrix}}{2} \\ &= \mathbf{C} + \frac{1}{2} \begin{pmatrix} 0 \\ 5 \\ -14 \end{pmatrix} \begin{pmatrix} 0 & 6 & 0 \end{pmatrix} = \mathbf{C} + \begin{pmatrix} 0 & 0 & 0 \\ 0 & 15 & 0 \\ 0 & -42 & 0 \end{pmatrix} \\ &= \begin{pmatrix} 0 & 6 & 0 \\ 5 & 0 & 5 \\ -14 & -14 & -11 \end{pmatrix} + \begin{pmatrix} 0 & 0 & 0 \\ 0 & 15 & 0 \\ 0 & -42 & 0 \end{pmatrix} \\ &= \begin{pmatrix} 0 & 6 & 0 \\ 5 & 15 & 5 \\ -14 & -56 & -11 \end{pmatrix}, \end{aligned}$$

$$
\begin{aligned}
|\lambda\,\mathbf{I}-\mathbf{C}_{\mathrm{d}}| &= \lambda^3-4\,\lambda^2+85\,\lambda+90=0,\\
\lambda(\mathbf{C}_{\mathrm{d}}) &= \left\{\frac{5+\sqrt{335}\,i}{2},\frac{5-\sqrt{335}\,i}{2},-1\right\}\\
\lambda_{\max}(\mathbf{C}_{\mathrm{d}})-\lambda_{\max}(\mathbf{D}) &= \frac{5}{2}-2=\frac{1}{2}>0.
\end{aligned}
\tag{S5.16}
$$

Hence, under equation (S5.10), the conditions for convergence stability ($\lambda_{\max}(\mathbf{C})<0$), evolutionary instability ($\lambda_{\max}(\mathbf{D})>0$), and dimorphic emergence ($\mathbf{e}^T[\mathbf{D}-\mathbf{C}]\,\mathbf{e}>0$) are all satisfied, but the condition for locally stable dimorphic divergence ($\lambda_{\max}(\mathbf{C}_{\mathrm{d}})<\lambda_{\max}(\mathbf{D})$) is not satisfied.

#### S5.2. Strongly convergence stable non-ESS

In an arbitrary trait space $\mathbf{s}=(x_1,...,x_L)^T$ of an arbitrary dimension $L\geqslant 2$, we consider a strongly convergence stable non-ESS denoted by $\mathbf{s}^*$, satisfying equations (S5.2) and (S5.5).

First, a strongly convergence stable non-ESS satisfies the condition for mutual invasibility along $\mathbf{e}$, because we see from $\lambda_{\max}(\mathbf{D})>0$ and $\lambda_{\max}\Big(\frac{\mathbf{C}+\mathbf{C}^T}{2}\Big)<0$ that

$$
\begin{aligned}
\mathbf{e}^T[\mathbf{D}-\mathbf{C}]\,\mathbf{e} &= \mathbf{e}^T\,\mathbf{D}\,\mathbf{e}-\mathbf{e}^T\,\mathbf{C}\,\mathbf{e}\\
&= \lambda_{\max}(\mathbf{D})-\mathbf{e}^T\left[\frac{\mathbf{C}+\mathbf{C}^T}{2}\right]\mathbf{e}\\
&\geqslant \lambda_{\max}(\mathbf{D})-\lambda_{\max}\left(\frac{\mathbf{C}+\mathbf{C}^T}{2}\right)>0.
\end{aligned}
\tag{S5.17}
$$

Second, a strongly convergence stable non-ESS satisfies the condition for locally stable dimorphic divergence, equation (S5.4), as derived below. Because a sufficient condition for $\lambda_{\max}(\mathbf{C})<0$ is given by $\lambda_{\max}\Big(\frac{\mathbf{C}^T+\mathbf{C}}{2}\Big)<0$, which is equivalent to that $\mathbf{w}^T\,\mathbf{C}\,\mathbf{w}<0$ holds for any $L$-dimensional vector $\mathbf{w}$, a sufficient condition for equation (S5.4) is also given by that

$$
\mathbf{w}^T\,\mathbf{C}_{\mathrm{d}}\,\mathbf{w} \;<\; 0 \tag{S5.18}
$$

holds for any $\mathbf{w}$. To confirm that this inequality holds good, we transform $\mathbf{w}^T\,\mathbf{C}_{\mathrm{d}}\,\mathbf{w}$ as

$$
\begin{aligned}
\mathbf{w}^T\,\mathbf{C}_{\mathrm{d}}\,\mathbf{w} &= \mathbf{w}^T\left[\mathbf{C}+\frac{\mathbf{C}\,\mathbf{e}\,\mathbf{e}^T\,\mathbf{C}}{\mathbf{e}^T[\mathbf{D}-\mathbf{C}]\,\mathbf{e}}\right]\mathbf{w}\\
&= \mathbf{w}^T\left[\frac{\mathbf{e}^T[\mathbf{D}-\mathbf{C}]\,\mathbf{e}\,\mathbf{C}+\mathbf{C}\,\mathbf{e}\,\mathbf{e}^T\,\mathbf{C}}{\mathbf{e}^T[\mathbf{D}-\mathbf{C}]\,\mathbf{e}}\right]\mathbf{w}\\
&= \mathbf{w}^T\left[\frac{\mathbf{e}^T\,\mathbf{D}\,\mathbf{e}\,\mathbf{C}-\mathbf{e}^T\,\mathbf{C}\,\mathbf{e}\,\mathbf{C}+\mathbf{C}\,\mathbf{e}\,\mathbf{e}^T\,\mathbf{C}}{\mathbf{e}^T[\mathbf{D}-\mathbf{C}]\,\mathbf{e}}\right]\mathbf{w}\\
&= \mathbf{w}^T\left[\frac{\mathbf{C}\,\mathbf{e}\,\mathbf{e}^T\,\mathbf{C}-\mathbf{e}^T\,\mathbf{C}\,\mathbf{e}\,\mathbf{C}}{\mathbf{e}^T[\mathbf{D}-\mathbf{C}]\,\mathbf{e}}+\frac{\mathbf{e}^T\,\mathbf{D}\,\mathbf{e}\,\mathbf{C}}{\mathbf{e}^T[\mathbf{D}-\mathbf{C}]\,\mathbf{e}}\right]\mathbf{w}\\
&= \frac{\mathbf{w}^T\,\mathbf{C}\,\mathbf{e}\,\mathbf{e}^T\,\mathbf{C}\,\mathbf{w}-\mathbf{e}^T\,\mathbf{C}\,\mathbf{e}\,\mathbf{w}^T\,\mathbf{C}\,\mathbf{w}}{\mathbf{e}^T[\mathbf{D}-\mathbf{C}]\,\mathbf{e}}+\frac{\lambda_{\max}(\mathbf{D})}{\mathbf{e}^T[\mathbf{D}-\mathbf{C}]\,\mathbf{e}}\mathbf{w}^T\,\mathbf{C}\,\mathbf{w}\\
&= \frac{[\mathbf{w}^T\,\mathbf{C}\,\mathbf{e}][\mathbf{w}^T\,\mathbf{C}^T\,\mathbf{e}]-[\mathbf{w}^T\,\mathbf{C}\,\mathbf{w}][\mathbf{e}^T\,\mathbf{C}\,\mathbf{e}]}{\mathbf{e}^T[\mathbf{D}-\mathbf{C}]\,\mathbf{e}}+\frac{\lambda_{\max}(\mathbf{D})\,\mathbf{w}^T\,\mathbf{C}\,\mathbf{w}}{\mathbf{e}^T[\mathbf{D}-\mathbf{C}]\,\mathbf{e}}.
\end{aligned}
\tag{S5.19}
$$

To transform equation (S5.19) further, we denote the symmetric and asymmetric parts of $\mathbf{C}$ by $\mathbf{C}^\circ$ and $\mathbf{C}^\bullet$, i.e.,

$$
\begin{aligned}
\mathbf{C} &= \mathbf{C}^\circ+\mathbf{C}^\bullet,\\
\mathbf{C}^T &= \mathbf{C}^\circ-\mathbf{C}^\bullet
\end{aligned}
\tag{S5.20}
$$

with $\lambda_{\max}(\mathbf{C}^\circ) = \lambda_{\max}\Big(\frac{\mathbf{C}+\mathbf{C}^T}{2}\Big) < 0$, where $\mathbf{e}^T\,\mathbf{C}^\bullet\,\mathbf{e} = 0$ and $\mathbf{w}^T\,\mathbf{C}^\bullet\,\mathbf{w} = 0$ hold. Substitution of equation (S5.20) into equation (S5.19) gives

$$\begin{aligned}
\mathbf{w}^T\,\mathbf{C}_\mathrm{d}\,\mathbf{w} &= \frac{\{\mathbf{w}^T[\mathbf{C}^\circ+\mathbf{C}^\bullet]\,\mathbf{e}\}\{\mathbf{w}^T[\mathbf{C}^\circ-\mathbf{C}^\bullet]\,\mathbf{e}\}}{\mathbf{e}^T[\mathbf{D}-\mathbf{C}]\,\mathbf{e}} - \frac{\mathbf{e}^T[\mathbf{C}^\circ+\mathbf{C}^\bullet]\,\mathbf{e}\,\mathbf{w}^T[\mathbf{C}^\circ+\mathbf{C}^\bullet]\,\mathbf{w}}{\mathbf{e}^T[\mathbf{D}-\mathbf{C}]\,\mathbf{e}} \\
&\quad + \frac{\lambda_{\max}(\mathbf{D})\,\mathbf{w}^T[\mathbf{C}^\circ+\mathbf{C}^\bullet]\,\mathbf{w}}{\mathbf{e}^T[\mathbf{D}-\mathbf{C}]\,\mathbf{e}} \\
&= \frac{[\mathbf{w}^T\,\mathbf{C}^\circ\,\mathbf{e}]^2-[\mathbf{w}^T\,\mathbf{C}^\bullet\,\mathbf{e}]^2}{\mathbf{e}^T[\mathbf{D}-\mathbf{C}]\,\mathbf{e}} - \frac{[\mathbf{e}^T\,\mathbf{C}^\circ\,\mathbf{e}][\mathbf{w}^T\,\mathbf{C}^\circ\,\mathbf{w}]}{\mathbf{e}^T[\mathbf{D}-\mathbf{C}]\,\mathbf{e}} + \frac{\lambda_{\max}(\mathbf{D})[\mathbf{w}^T\,\mathbf{C}^\circ\,\mathbf{w}]}{\mathbf{e}^T[\mathbf{D}-\mathbf{C}]\,\mathbf{e}} \\
&= \frac{[\mathbf{w}^T\,\mathbf{C}^\circ\,\mathbf{e}]^2-[\mathbf{w}^T\,\mathbf{C}^\circ\,\mathbf{w}][\mathbf{e}^T\,\mathbf{C}^\circ\,\mathbf{e}]}{\mathbf{e}^T[\mathbf{D}-\mathbf{C}]\,\mathbf{e}} - \frac{[\mathbf{w}^T\,\mathbf{C}^\bullet\,\mathbf{e}]^2}{\mathbf{e}^T[\mathbf{D}-\mathbf{C}]\,\mathbf{e}} + \frac{\lambda_{\max}(\mathbf{D})[\mathbf{w}^T\,\mathbf{C}^\circ\,\mathbf{w}]}{\mathbf{e}^T[\mathbf{D}-\mathbf{C}]\,\mathbf{e}} \\
&\leqslant \frac{[\mathbf{w}^T\,\mathbf{C}^\circ\,\mathbf{e}]^2-[\mathbf{w}^T\,\mathbf{C}^\circ\,\mathbf{w}][\mathbf{e}^T\,\mathbf{C}^\circ\,\mathbf{e}]}{\mathbf{e}^T[\mathbf{D}-\mathbf{C}]\,\mathbf{e}} + \frac{\lambda_{\max}(\mathbf{D})|\mathbf{w}|^2\,\lambda_{\max}(\mathbf{C}^\circ)}{\mathbf{e}^T[\mathbf{D}-\mathbf{C}]\,\mathbf{e}} \\
&< \frac{[\mathbf{w}^T\,\mathbf{C}^\circ\,\mathbf{e}]^2-[\mathbf{w}^T\,\mathbf{C}^\circ\,\mathbf{w}][\mathbf{e}^T\,\mathbf{C}^\circ\,\mathbf{e}]}{\mathbf{e}^T[\mathbf{D}-\mathbf{C}]\,\mathbf{e}} \qquad \text{(S5.21)}
\end{aligned}$$

To transform equation (S5.21) further, we decompose $\mathbf{C}^\circ$ as

$$\begin{aligned}
-\mathbf{C}^\circ &= \mathbf{E}^\circ\,\mathrm{diag}(\lambda_1^\circ,...,\lambda_L^\circ)\,\mathbf{E}^{\circ T} = \sqrt{\mathbf{C}^\circ}^T\sqrt{\mathbf{C}^\circ}, \\
\sqrt{\mathbf{C}^\circ} &= \mathrm{diag}\big(\sqrt{\lambda_1^\circ},...,\sqrt{\lambda_L^\circ}\big)^T\,\mathbf{E}^{\circ T}, \qquad \text{(S5.22)}
\end{aligned}$$

and introduce

$$\begin{aligned}
\mathbf{w}^\circ &= \sqrt{\mathbf{C}^\circ}\,\mathbf{w}, \\
\mathbf{e}^\circ &= \sqrt{\mathbf{C}^\circ}\,\mathbf{e}, \qquad \text{(S5.23)}
\end{aligned}$$

which upon substitution into equation (S5.21) gives

$$\begin{aligned}
\mathbf{w}^T\,\mathbf{C}_\mathrm{d}\,\mathbf{w} &< \frac{[-\mathbf{w}^T\sqrt{\mathbf{C}^\circ}^T\sqrt{\mathbf{C}^\circ}\,\mathbf{e}]^2-[-\mathbf{w}^T\sqrt{\mathbf{C}^\circ}^T\sqrt{\mathbf{C}^\circ}\,\mathbf{w}][-\mathbf{e}^T\sqrt{\mathbf{C}^\circ}^T\sqrt{\mathbf{C}^\circ}\,\mathbf{e}]}{\mathbf{e}^T[\mathbf{D}-\mathbf{C}]\,\mathbf{e}} \\
&= \frac{[\mathbf{w}^{\circ T}\,\mathbf{e}^\circ]^2-[\mathbf{w}^{\circ T}\,\mathbf{w}^\circ][\mathbf{e}^{\circ T}\,\mathbf{e}^\circ]}{\mathbf{e}^T[\mathbf{D}-\mathbf{C}]\,\mathbf{e}} \\
&= \frac{|\mathbf{w}^\circ|^2|\mathbf{e}^\circ|^2}{\mathbf{e}^T[\mathbf{D}-\mathbf{C}]\,\mathbf{e}}\left\{\left(\left[\frac{\mathbf{w}^\circ}{|\mathbf{w}^\circ|}\right]^T\left[\frac{\mathbf{e}^\circ}{|\mathbf{e}^\circ|}\right]\right)^2-1\right\}\leqslant 0. \qquad \text{(S5.24)}
\end{aligned}$$

Therefore, $\lambda_{\max}\Big(\frac{\mathbf{C}_\mathrm{d}+\mathbf{C}_\mathrm{d}^T}{2}\Big) < 0$ holds and hence $\lambda_{\max}(\mathbf{C}_\mathrm{d}) < 0$ holds, which satisfies the condition for locally stable dimorphic divergence, equation (S5.4).

##### S5.3. Absolutely convergence stable non-ESS

Since any absolutely convergence stable non-ESS is also a strongly convergence stable non-ESS, an absolutely convergence stable non-ESS also satisfies the branching possibility conditions.

#### S6. Quantitative index for branching likelihood

##### S6.1. Outline of this section

Because what the branching possibility conditions ensure is not a high probability but a non-zero probability for evolutionary branching (equations S3.11 in Appendix S3.1), the probability for evolutionary branching under some possible or strongly possible branching points could be very low. Indeed, when the real parts of the eigenvalues of $\mathbf{C}$ have much smaller absolute values than the imaginary parts, numerically simulated evolution tends to show monomorphic directional evolution that orbits around the possible (or strongly possible) branching point for a large number of invasions without the s occurrence of evolutionary branching. For a simple example, a point with

$$\mathbf{C} = \begin{pmatrix} -a & -1 \\ 1 & -a \end{pmatrix}, \ \mathbf{D} = \begin{pmatrix} a & 0 \\ 0 & -a \end{pmatrix}, \ \mathbf{V}_\mu = \sigma^2 \begin{pmatrix} 1 & 0 \\ 0 & 1 \end{pmatrix} \tag{S6.1}$$

with $a$ and $\sigma$ being positive constants is a strongly possible branching point (where $\sigma$ describes the average mutational step size and is assumed to satisfy $0 < \sigma \ll 1$). However, when $a$ is much smaller than 1, the monomorphic directional evolution tends to show a circular orbit around $\mathbf{s}^*$ characterized by a circle with its radius $\sigma / a$ being much larger than $\sigma$ (figure S3f). In this case, the population is kept away from $\mathbf{s}^*$ so that the emergence of dimorphism and their divergence are extremely unlikely, despite that $\mathbf{s}^*$ is a strongly possible branching point (figure S3b).

To roughly estimate the likelihood of evolutionary branching for possible branching points, a sufficient condition for the emergence of dimorphism (along $\mathbf{e}$ during directional evolution in the circular orbit around $\mathbf{s}^*$) is approximately derived in the subsequent sections as

$$Q = \frac{\mathbf{e}^T[\mathbf{D} - \mathbf{C}]\,\mathbf{e}}{\mathbf{e}^T\,\mathbf{V}_\mu^{-1}\,\mathbf{e}} \min_{i \in \{1,\ldots,L\}} \left( \frac{|\mathrm{Re}(\lambda_i(\mathbf{V}_\mu\,\mathbf{C}^T))|}{\mathrm{Re}(\lambda_i(\mathbf{V}_\mu\,\mathbf{C}^T))^2 + \mathrm{Im}(\lambda_i(\mathbf{V}_\mu\,\mathbf{C}^T))^2} \right) > 1, \tag{S6.2}$$

with $\lambda_i(\mathbf{V}_\mu\ \mathbf{C}^T)$ describing the $i$th eigenvalue of $\mathbf{V}_\mu\ \mathbf{C}^T$. On this basis, we may expect a high likelihood of evolutionary branching when $Q > 1$ holds. For example, equation (S6.1) gives $Q = 2\ a^2 / [1 + a^2]$ for $a \ll 1$. As far as numerically examined (figures S3-S8), $Q$ seems useful as a quantitative index for the likelihood of evolutionary branching in simulated evolution as trait substitution sequences.

#### S6.2. Derivation for two-dimensional trait spaces

Here we consider a monomorphic population of a phenotype in the neighborhood of a possible branching point $\mathbf{s}^* = \mathbf{0}$ in a two-dimensional trait space $\mathbf{s} = (x_1, x_2)^T$. We assume that the trait space has been normalized so that the mutational covariance matrix is given by $\mathbf{V}_\mu = \sigma^2\,\mathbf{I}$. Then the branching possibility conditions, equation (15), are simplified into

$$\begin{aligned}
\lambda_{\max}(\mathbf{C}^T) &< 0 && \text{(convergence stability)}, && \text{(S6.3)} \\
\lambda_{\max}(\mathbf{D}) &> 0 && \text{(evolutionary instability)}, && \text{(S6.4)} \\
\mathbf{e}^T[\mathbf{D} - \mathbf{C}]\,\mathbf{e} &> 0 && \text{(dimorphic emergence: mutual invasibility)}, && \text{(S6.5)} \\
\lambda_{\max}(\mathbf{C}_\mathrm{d}^T) &< \lambda_{\max}(\mathbf{D}) && \text{(locally stable dimorhpic divergence)}, && \text{(S6.6)} \\
\mathbf{C}_\mathrm{d} &= \mathbf{C} + \frac{\mathbf{C}\,\mathbf{e}\,\mathbf{e}^T\,\mathbf{C}}{\mathbf{e}^T[\mathbf{D} - \mathbf{C}]\,\mathbf{e}}.
\end{aligned}$$

Below, we approximately derive a sufficient condition for the emergence of dimorphism along $\mathbf{e}$ during the directional evolution going around $\mathbf{s}^*$. To describe the directional evolution of a monomorphic population going around $\mathbf{s}^* = \mathbf{0}$, we denote the monomorphic resident by $\mathbf{s}$ and describe a typical invading mutant as

$$\begin{aligned}
\mathbf{s}' &= \mathbf{s} + \mathbf{e}_g\,\sigma, \\
\mathbf{e}_g &= \frac{\mathbf{g}(\mathbf{s})}{|\mathbf{g}(\mathbf{s})|} = \frac{\mathbf{C}^T\,\mathbf{s}}{|\mathbf{C}^T\,\mathbf{s}|}
\end{aligned} \tag{S6.7}$$

with $\mathbf{g}(\mathbf{s}) = \mathbf{C}^T \mathbf{s}$ describing the fitness gradient for $\mathbf{s}$. For simplicity of the derivation, we assume that the population is going around $\mathbf{s}^*$ so that $|\mathbf{s}| \gg \sigma$ is kept. In this case, the mutant $\mathbf{s}' = \mathbf{s} + \mathbf{e}_g \sigma$ always replaces the resident $\mathbf{s}$, when the following inequalities both hold:

$$\begin{aligned} f(\mathbf{s}';\mathbf{s}) &= |\mathbf{s}^T \mathbf{C}|\,\sigma + \frac{1}{2}\mathbf{e}_g^T \mathbf{D}\,\mathbf{e}_g \sigma^2 \simeq |\mathbf{s}^T \mathbf{C}|\,\sigma > 0, \\ f(\mathbf{s};\mathbf{s}') &= -[\mathbf{s} + \mathbf{e}_g \sigma]^T \mathbf{C}\,\mathbf{e}_g \sigma + \frac{1}{2}\mathbf{e}_g^T \mathbf{D}\,\mathbf{e}_g \sigma^2 \simeq -|\mathbf{s}^T \mathbf{C}|\,\sigma < 0. \end{aligned} \tag{S6.8}$$

Because $\lambda_{\max}(\mathbf{C}^T) < 0$ holds, we can find a real matrix $\mathbf{P}$ so that $\mathbf{C}^T$ is decomposed as

$$\begin{aligned} \mathbf{C}^T &= \mathbf{P}\,\mathbf{A}\,\mathbf{P}^{-1} \\ \mathbf{A} &= \begin{pmatrix} \mathrm{Re}(\lambda_1) & \mathrm{Im}(\lambda_1) \\ \mathrm{Im}(\lambda_2) & \mathrm{Re}(\lambda_2) \end{pmatrix}, \end{aligned} \tag{S6.9}$$

where $\mathrm{Re}(\lambda_1) = \mathrm{Re}(\lambda_2)$ and $\mathrm{Im}(\lambda_1) = -\mathrm{Im}(\lambda_2)$ hold when $\lambda_1$ and $\lambda_2$ are complex numbers. Then we introduce

$$\mathbf{z} = \begin{pmatrix} z_1 \\ z_2 \end{pmatrix} = \mathbf{P}^{-1}\mathbf{s}, \tag{S6.10}$$

and transform $\mathbf{z}' = \mathbf{P}^{-1}\mathbf{s}'$ as

$$\begin{aligned} \mathbf{z}' &= \mathbf{P}^{-1}\mathbf{s}' = \mathbf{P}^{-1}[\mathbf{s} + \mathbf{e}_g \sigma] = \mathbf{P}^{-1}\left[\mathbf{s} + \frac{\mathbf{C}^T\mathbf{s}}{|\mathbf{C}^T\mathbf{s}|}\sigma\right] \\ &= \mathbf{z} + \frac{\mathbf{P}^{-1}\mathbf{C}^T\mathbf{P}\,\mathbf{z}}{|\mathbf{C}^T\mathbf{P}\,\mathbf{z}|}\sigma = \mathbf{z} + \frac{\mathbf{A}\,\mathbf{z}}{|\mathbf{C}^T\mathbf{P}\,\mathbf{z}|}\sigma, \end{aligned} \tag{S6.11}$$

from which we derive

$$\begin{aligned} |\mathbf{z}'|^2 - |\mathbf{z}|^2 &= \left[\mathbf{z} + \frac{\mathbf{A}\,\mathbf{z}}{|\mathbf{C}^T\mathbf{P}\,\mathbf{z}|}\sigma\right]^T \left[\mathbf{z} + \frac{\mathbf{A}\,\mathbf{z}}{|\mathbf{C}^T\mathbf{P}\,\mathbf{z}|}\sigma\right] - \mathbf{z}^T\mathbf{z} \\ &= 2\frac{\mathbf{z}^T\mathbf{A}\,\mathbf{z}}{|\mathbf{C}^T\mathbf{P}\,\mathbf{z}|}\sigma + \frac{\mathbf{z}^T\mathbf{A}^T\mathbf{A}\,\mathbf{z}}{|\mathbf{C}^T\mathbf{P}\,\mathbf{z}|^2}\sigma^2 \\ &= \frac{2\,\mathbf{z}^T\mathbf{A}\,\mathbf{z}\,\sigma}{|\mathbf{C}^T\mathbf{P}\,\mathbf{z}|^2}\left\{|\mathbf{C}^T\mathbf{P}\,\mathbf{z}| + \frac{\mathbf{z}^T\mathbf{A}^T\mathbf{A}\,\mathbf{z}}{\mathbf{z}^T\mathbf{A}\,\mathbf{z}}\frac{\sigma}{2}\right\}. \end{aligned} \tag{S6.12}$$

Note that the following relationships hold

$$\begin{aligned} \mathbf{z}^T\mathbf{A}\,\mathbf{z} &= (\,z_1, z_2\,)\begin{pmatrix} \mathrm{Re}(\lambda_1) & \mathrm{Im}(\lambda_1) \\ \mathrm{Im}(\lambda_2) & \mathrm{Re}(\lambda_2) \end{pmatrix}\begin{pmatrix} z_1 \\ z_2 \end{pmatrix} = \mathrm{Re}(\lambda_1)\,z_1^2 + \mathrm{Re}(\lambda_2)\,z_2^2 < 0, \\ \mathbf{z}^T\mathbf{A}^T\mathbf{A}\,\mathbf{z} &= (\,z_1, z_2\,)\begin{pmatrix} \mathrm{Re}(\lambda_1)^2 + \mathrm{Im}(\lambda_1)^2 & 0 \\ 0 & \mathrm{Re}(\lambda_2)^2 + \mathrm{Im}(\lambda_2)^2 \end{pmatrix}\begin{pmatrix} z_1 \\ z_2 \end{pmatrix} \\ &= [\mathrm{Re}(\lambda_1)^2 + \mathrm{Im}(\lambda_1)^2]\,z_1^2 + [\mathrm{Re}(\lambda_2)^2 + \mathrm{Im}(\lambda_2)^2]\,z_2^2 > 0. \end{aligned} \tag{S6.13}$$

As long as directional evolution continues so that the mutant always replaces the resident, we expect that the population eventually orbit $\mathbf{s}^*$, roughly satisfying $|\mathbf{z}'|^2 = |\mathbf{z}|^2$. Substitution of $|\mathbf{z}'|^2 \simeq |\mathbf{z}|^2$ and equation (S6.13) into equation (S6.12) gives a rough magnitude estimation for $|\mathbf{C}^T\mathbf{s}|$:

$$\begin{aligned} |\mathbf{C}^T\mathbf{P}\,\mathbf{z}| &= |\mathbf{C}^T\mathbf{s}| \simeq \max_{\mathbf{z}}\left(\frac{\mathbf{z}^T\mathbf{A}^T\mathbf{A}\,\mathbf{z}}{|\mathbf{z}^T\mathbf{A}\,\mathbf{z}|}\right)\frac{\sigma}{2} = W_{\max}\frac{\sigma}{2}, \\ W_{\max} &= \max_{\mathbf{z}}\left(\frac{\mathbf{z}^T\mathbf{A}^T\mathbf{A}\,\mathbf{z}}{|\mathbf{z}^T\mathbf{A}\,\mathbf{z}|}\right) \\ &= \max_{\mathbf{z}}\left(\frac{[\mathrm{Re}(\lambda_1)^2 + \mathrm{Im}(\lambda_1)^2]\,z_1^2 + [\mathrm{Re}(\lambda_2)^2 + \mathrm{Im}(\lambda_2)^2]\,z_2^2}{|\mathrm{Re}(\lambda_1)\,z_1^2 + \mathrm{Re}(\lambda_2)\,z_2^2|}\right) \\ &= \max_{i\in\{1,2\}}\left(\frac{[\mathrm{Re}(\lambda_i)^2 + \mathrm{Im}(\lambda_i)^2]}{|\mathrm{Re}(\lambda_i)|}\right). \end{aligned} \tag{S6.14}$$

Hence, we expect that $|\mathbf{C}^T\, \mathbf{s}| = W_{\max}\, \frac{\sigma}{2}$ characterizes the size and shape of the orbit. In the simple case such that $\mathbf{P} = \mathbf{I}$ holds and that $\mathbf{C}$ has complex eigenvalues (i.e., $\mathrm{Re}(\lambda_1) = \mathrm{Re}(\lambda_2)$ and $\mathrm{Im}(\lambda_1) = -\mathrm{Im}(\lambda_2)$), the dynamics of $\mathbf{s}$ is expected to be isotropic, in which case the orbit radius can be derived from equation (S6.14) as

$$|\mathbf{s}| \;\simeq\; \frac{\sqrt{\mathrm{Re}(\lambda_1)^2 + \mathrm{Im}(\lambda_1)^2}}{2|\mathrm{Re}(\lambda_1)|}\,\sigma. \tag{S6.15}$$

Next, we approximately derive a sufficient condition for a mutant $\mathbf{s}' = \mathbf{s} + \mathbf{e}_g\,\sigma$ coexisting with its resident $\mathbf{s}$ when $\mathbf{e}_g \simeq \mathbf{e}$ is realized in the orbit. We transform $f(\mathbf{s}';\mathbf{s})$ and $f(\mathbf{s};\mathbf{s}')$ as

$$\begin{aligned}
f(\mathbf{s}';\mathbf{s}) \;&=\; \mathbf{s}^T\,\mathbf{C}[\mathbf{s}'-\mathbf{s}] + \frac{1}{2}[\mathbf{s}'-\mathbf{s}]^T\,\mathbf{D}[\mathbf{s}'-\mathbf{s}] \\
&=\; \left[\frac{\mathbf{s}+\mathbf{s}'}{2}\right]^T \mathbf{C}[\mathbf{s}'-\mathbf{s}] + \frac{1}{2}[\mathbf{s}'-\mathbf{s}]^T[\mathbf{D}-\mathbf{C}][\mathbf{s}'-\mathbf{s}] \\
&=\; \left[\mathbf{s} + \frac{\mathbf{e}_g\,\sigma}{2}\right]^T \mathbf{C}\,\mathbf{e}_g\,\sigma + \frac{1}{2}\,\mathbf{e}^T[\mathbf{D}-\mathbf{C}]\,\mathbf{e}\,\sigma^2 \\
&\simeq\; \mathbf{s}^T\,\mathbf{C}\,\mathbf{e}_g\,\sigma + \frac{1}{2}\,\mathbf{e}^T[\mathbf{D}-\mathbf{C}]\,\mathbf{e}\,\sigma^2 \\
&=\; |\mathbf{s}^T\,\mathbf{C}|\,\sigma + \frac{1}{2}\,\mathbf{e}^T[\mathbf{D}-\mathbf{C}]\,\mathbf{e}\,\sigma^2 \\
&\simeq\; W_{\max}\frac{\sigma^2}{2} + \frac{1}{2}\,\mathbf{e}^T[\mathbf{D}-\mathbf{C}]\,\mathbf{e}\,\sigma^2 \\
&=\; W_{\max}\frac{\sigma^2}{2}\left\{1 + \frac{\mathbf{e}^T[\mathbf{D}-\mathbf{C}]\,\mathbf{e}}{W_{\max}}\right\}, \\
f(\mathbf{s};\mathbf{s}') \;&=\; \mathbf{s}'^T\,\mathbf{C}[\mathbf{s}-\mathbf{s}'] + \frac{1}{2}[\mathbf{s}-\mathbf{s}']^T\,\mathbf{D}[\mathbf{s}-\mathbf{s}'] \\
&\simeq\; W_{\max}\frac{\sigma^2}{2}\left\{-1 + \frac{\mathbf{e}^T[\mathbf{D}-\mathbf{C}]\,\mathbf{e}}{W_{\max}}\right\}.
\end{aligned} \tag{S6.16}$$

Hence a sufficient condition for their coexistence, i.e., $f(\mathbf{s}';\mathbf{s}) > 0$ and $f(\mathbf{s};\mathbf{s}') > 0$, is approximately given by

$$\begin{aligned}
Q \;&:=\; \frac{\mathbf{e}^T[\mathbf{D}-\mathbf{C}]\,\mathbf{e}}{W_{\max}} > 1, \\
W_{\max} \;&=\; \max_{i\in\{1,2\}}\left(\frac{[\mathrm{Re}(\lambda_i)^2 + \mathrm{Im}(\lambda_i)^2]}{|\mathrm{Re}(\lambda_i)|}\right).
\end{aligned} \tag{S6.17}$$

Therefore, when equation (S6.17) holds, we may expect that dimorphism can easily emerge along $\mathbf{e}$ during the directional evolution going around $\mathbf{s}^*$.

#### S6.3. Higher-dimensional trait space

For an $L$-dimensional trait space, in a manner analogous to the two-dimensional case in Section S5.1, a sufficient condition for the emergence of dimorphism along $\mathbf{e}$ during the directional evolution going around $\mathbf{s}^*$ is approximately given by

$$\begin{aligned}
Q \;&=\; \frac{\mathbf{e}^T[\mathbf{D}-\mathbf{C}]\,\mathbf{e}}{W_{\max}} > 1, \\
W_{\max} \;&=\; \max_{i\in\{1,\ldots,L\}}\left(\frac{\mathrm{Re}(\lambda_i)^2 + \mathrm{Im}(\lambda_i)^2}{|\mathrm{Re}(\lambda_i)|}\right),
\end{aligned} \tag{S6.18}$$

where $\lambda_i$ is the $i$th eigenvalue of $\mathbf{C}$.

#### S6.4. Higher-dimensional non-normalized trait spaces

We consider an $L$-dimensional trait space, where the mutational covariance matrix is not proportional to the identity. We describe the mutation probability distribution for emergence of $\mathbf{s}'$ from $\mathbf{s}$ as

$$P_\mu(\mathbf{s}'-\mathbf{s}_i) \;=\; \frac{1}{\sqrt{[2\,\pi]^L|\mathbf{V}_\mu|}}\exp\!\Big(-\frac{1}{2}[\mathbf{s}'-\mathbf{s}_i]^T\,\mathbf{V}_\mu^{-1}[\mathbf{s}'-\mathbf{s}_i]\Big). \tag{S6.19}$$

Following Appendix S3.7, we express the invasion fitness function in the normalized trait space $\mathbf{u}$ (with mutational covariance matrix $\mathbf{V}_\mu=\sigma^2\,\mathbf{I}$) for a mutant $\mathbf{u}'$ under a monomorphic resident $\mathbf{u}$ as

$$\tilde{f}(\mathbf{u}',\mathbf{u}) \;=\; \mathbf{u}^T\,\tilde{\mathbf{C}}[\mathbf{u}'-\mathbf{u}]+\frac{1}{2}[\mathbf{u}'-\mathbf{u}]^T\,\tilde{\mathbf{D}}[\mathbf{u}'-\mathbf{u}] \tag{S6.20}$$

with

$$\begin{aligned}
\tilde{\mathbf{C}} \;&=\; \sigma^{-2}\sqrt{\mathbf{V}_\mu}\,\mathbf{C}\,\sqrt{\mathbf{V}_\mu}^T,\\
\tilde{\mathbf{D}} \;&=\; \sigma^{-2}\sqrt{\mathbf{V}_\mu}\,\mathbf{D}\,\sqrt{\mathbf{V}_\mu}^T,\\
\lambda_i(\tilde{\mathbf{C}}^T) \;&=\; \sigma^{-2}\,\lambda_i(\mathbf{V}_\mu\,\mathbf{C}^T),\\
\lambda_{\max}(\tilde{\mathbf{C}}^T) \;&=\; \sigma^{-2}\,\lambda_{\max}(\mathbf{V}_\mu\,\mathbf{C}^T),\\
\tilde{\mathbf{e}}^T\,\tilde{\mathbf{D}}\,\tilde{\mathbf{e}}=\lambda_{\max}(\tilde{\mathbf{D}}) \;&=\; \sigma^{-2}\,\frac{\mathbf{e}^T\,\mathbf{D}\,\mathbf{e}}{\mathbf{e}^T\,\mathbf{V}_\mu^{-1}\,\mathbf{e}},\\
\tilde{\mathbf{e}}^T[\tilde{\mathbf{D}}-\tilde{\mathbf{C}}]\,\tilde{\mathbf{e}} \;&=\; \sigma^{-2}\,\frac{\mathbf{e}^T[\mathbf{D}-\mathbf{C}]\,\mathbf{e}}{\mathbf{e}^T\,\mathbf{V}_\mu^{-1}\,\mathbf{e}},\\
\tilde{\mathbf{e}} \;&=\; \frac{\tilde{\mathbf{v}}}{|\tilde{\mathbf{v}}|}=\frac{\mathbf{v}_{\max}(\tilde{\mathbf{D}})}{\mathbf{v}_{\max}(\tilde{\mathbf{D}})}=\frac{\mathbf{v}_{\max}\big(\sqrt{\mathbf{V}_\mu}\,\mathbf{D}\,\sqrt{\mathbf{V}_\mu}^T\big)}{\mathbf{v}_{\max}\big(\sqrt{\mathbf{V}_\mu}\,\mathbf{D}\,\sqrt{\mathbf{V}_\mu}^T\big)},\\
\tilde{\mathbf{e}} \;&=\; \frac{\mathbf{W}^{-1}\,\mathbf{e}}{|\mathbf{W}^{-1}\,\mathbf{e}|},
\end{aligned} \tag{S6.21}$$

where $\lambda_i(\tilde{\mathbf{C}}^T)$ is the $i$th eigenvalue of $\tilde{\mathbf{C}}^T$. In a normalized trait space, a sufficient condition for the emergence of dimorphism along $\tilde{\mathbf{e}}$ during the directional evolution going around $\mathbf{s}^*$ is approximately given by

$$\begin{aligned}
Q \;&=\; \frac{\tilde{\mathbf{e}}^T[\tilde{\mathbf{D}}-\tilde{\mathbf{C}}]\,\tilde{\mathbf{e}}}{\tilde{W}_{\max}}>1,\\
\tilde{W}_{\max} \;&=\; \max_{i\in\{1,\ldots,L\}}\left(\frac{\mathrm{Re}(\lambda_i(\tilde{\mathbf{C}}^T))^2+\mathrm{Im}(\lambda_i(\tilde{\mathbf{C}}^T))^2}{|\mathrm{Re}(\lambda_i(\tilde{\mathbf{C}}^T))|}\right),
\end{aligned} \tag{S6.22}$$

By using equation (S6.21), we transform equation (S6.18) into equation (S6.2):

$$\begin{aligned}
Q \;&=\; \sigma^{-2}\,\frac{\mathbf{e}^T[\mathbf{D}-\mathbf{C}]\,\mathbf{e}}{\mathbf{e}^T\,\mathbf{V}_\mu^{-1}\,\mathbf{e}}\min_{i\in\{1,\ldots,L\}}\left(\frac{|\mathrm{Re}(\lambda_i(\tilde{\mathbf{C}}^T))|}{\mathrm{Re}(\lambda_i(\tilde{\mathbf{C}}^T))^2+\mathrm{Im}(\lambda_i(\tilde{\mathbf{C}}^T))^2}\right)\\
&=\; \frac{\mathbf{e}^T[\mathbf{D}-\mathbf{C}]\,\mathbf{e}}{\mathbf{e}^T\,\mathbf{V}_\mu^{-1}\,\mathbf{e}}\min_{i\in\{1,\ldots,L\}}\left(\frac{|\mathrm{Re}(\lambda_i(\mathbf{V}_\mu\,\mathbf{C}^T))|}{\mathrm{Re}(\lambda_i(\mathbf{V}_\mu\,\mathbf{C}^T))^2+\mathrm{Im}(\lambda_i(\mathbf{V}_\mu\,\mathbf{C}^T))^2}\right)>1.
\end{aligned} \tag{S6.23}$$

#### S6.5. Numerical examination

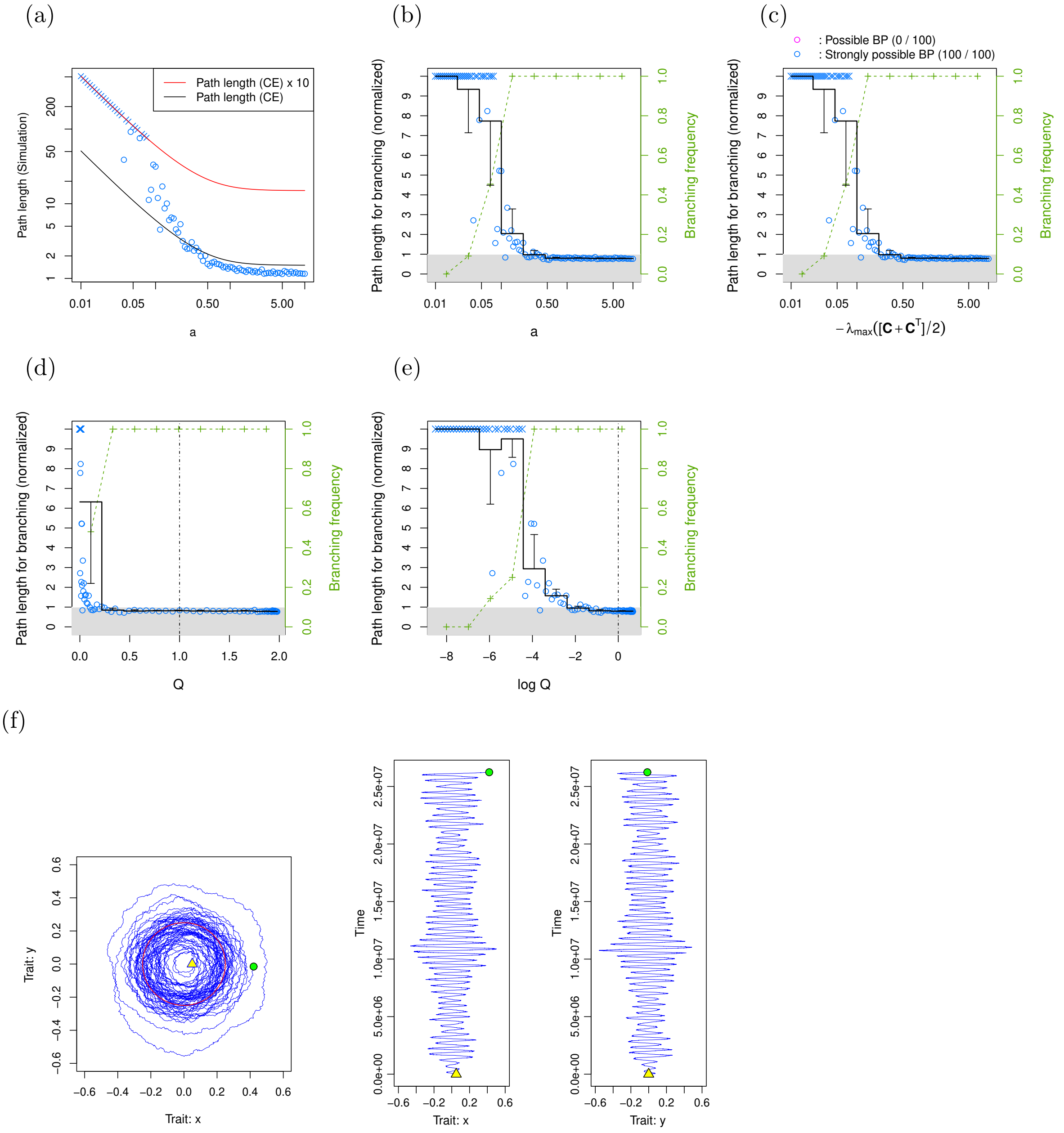

**Figure S3.** Effect of the imaginary parts of eigenvalues of $\mathbf{C}$ on likelihood of evolutionary branching in trait substitution sequences induced by a strongly possible branching point in a two-dimensional trait space. $\mathbf{C}$, $\mathbf{D}$, and $\mathbf{V}_\mu$ are given by equation (S6.1) with 100 different values for $a$ chosen from 0.01 to 10 in a geometric manner. The initial monomorphic resident was fixed at $\mathbf{s}_0 = (100\,\sigma, 0)^T$. Also, $\sigma$ was fixed at $\sigma = 0.005$. Occurrence of evolutionary branching was judged by the emergence of dimorphic residents with their phenotypic difference larger than $\Delta\,s = 100\,\sigma$. Each simulation has ended when evolutionary branching has occurred or when the path length ($\sigma\,K$ with $K$ describing the numbers of invasions) has exceeded $10\,l_{\rm CE}(\mathbf{s}_0, \Delta\,s)$, where $l_{\rm CE}(\mathbf{s}_0, \Delta\,s)$ describes the path length calculated with the canonical equation from the initial monomorphic resident $\mathbf{s}_0$ to the emergence of dimorphism with $\Delta\,s$, defined in equation (S.2.70).

In panel (a), blue small "$\circ$"s and "$\times$"s indicate occurrence and non-occurrence of evolutionary branching with $a$ for the horizontal axis and the non-normalized path length for vertical axis, where red and black curves indicate $10\,l_{\rm CE}(\mathbf{s}_0, \Delta\,s)$ and $l_{\rm CE}(\mathbf{s}_0, \Delta\,s)$, respectively. In panel (b), blue small "$\circ$"s and "$\times$"s indicate occurrence and non-occurrence of evolutionary branching with $a$ for the horizontal axis and the normalized path lengths ($\sigma\,K\,/\,l_{\rm CE}(\mathbf{s}_0, \Delta\,s)$) for the vertical axis. The black staircase plot and the green small "+"s indicate the mean path length (with errorbars indicating standard deviations) and the branching frequency, respectively, for subdivided regions of the horizontal axis. Panels (c-f) are plotted in the same manner as panel (b) except that the horizontal axes is $-\lambda_{\max}([\mathbf{C}+\mathbf{C}^T]\,/\,2)$ for (c), $Q$ for (d), $\log Q$ for (e), where $\lambda_{\max}(\circ)$ is the maximum value among the real parts of the eigenvalues of matrix $\circ$, and where $Q$ is defined in equation (S6.17). Vertical dashed lines in (d)

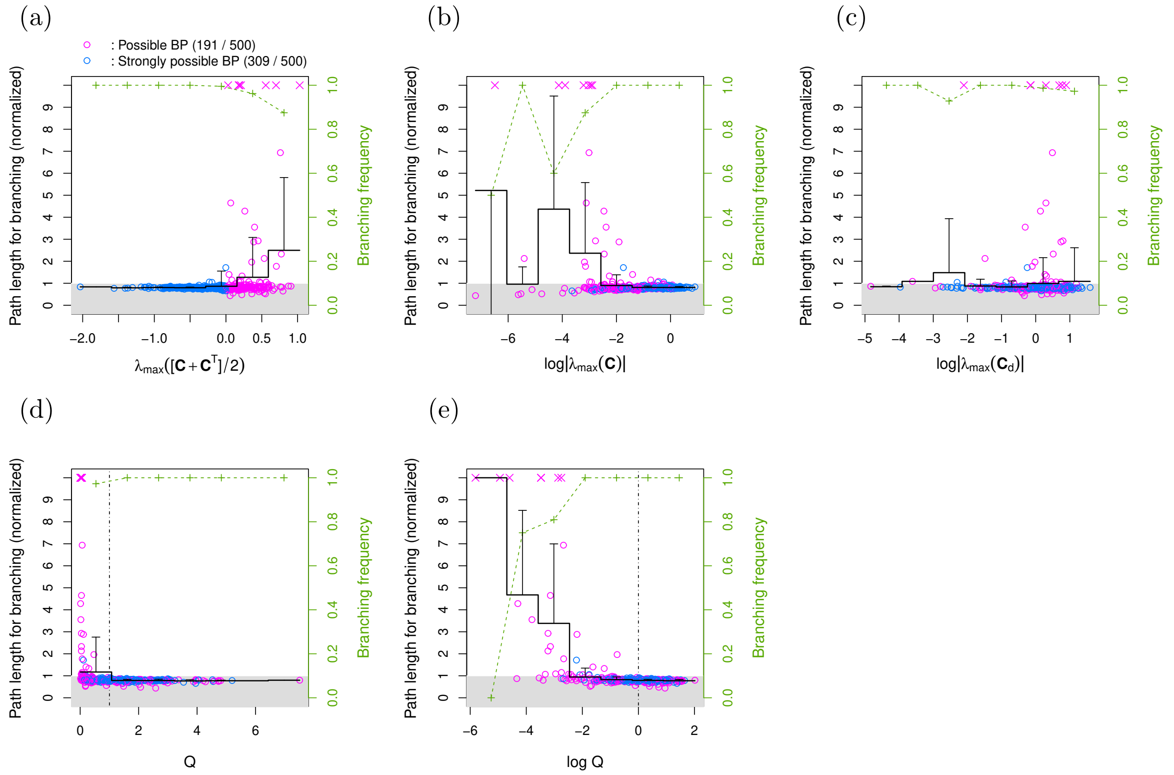

**Figure S4.** Examination of normalized path lengths for the occurrence of evolutionary branching in simulated evolution as trait substitution sequences (two-dimensional trait spaces, i.e., $L = 2$). Plotting formats for panels (a-g) are the same as those shown in panels (b-e) of figure S3, except that blue and magenta colors indicate strongly possible branching points and possible branching points that are not strongly convergence stable, respectively . The simulation was repeated 500 times, where each simulation was conducted in the same manner with the simulation shown in figure S3, except that the initial phenotype $\mathbf{s}_0$ and the set of $\mathbf{C}$ and $\mathbf{D}$ (satisfying the branching possibility conditions) were randomly chosen as explained below. $\mathbf{s}_0$ was chosen from an isotropic $L$-dimensional Gaussian distribution with its mean at the origin and covariance matrix given by $200\,\sigma^2\,\mathbf{I}$. Regarding $\mathbf{C}$ and $\mathbf{D}$, the first 500 sets of $\mathbf{C}$ and $\mathbf{D}$ satisfying the branching possibility conditions, equation (15) in the main text, were chosen from a large number of randomly generated matrices. For each random generation of $\mathbf{C}$, its each entry was chosen from a one-dimensional Gaussian distribution with mean 0 and standard deviation 1, denoted by $\mathcal{N}(0, 1)$. For each random generation of $\mathbf{D}$, which was assumed to be a diagonal matrix without loss of generality, its each diagonal entry was chosen from $\mathcal{N}(0, 1)$, and then the diagonal entries were sorted in descending order.

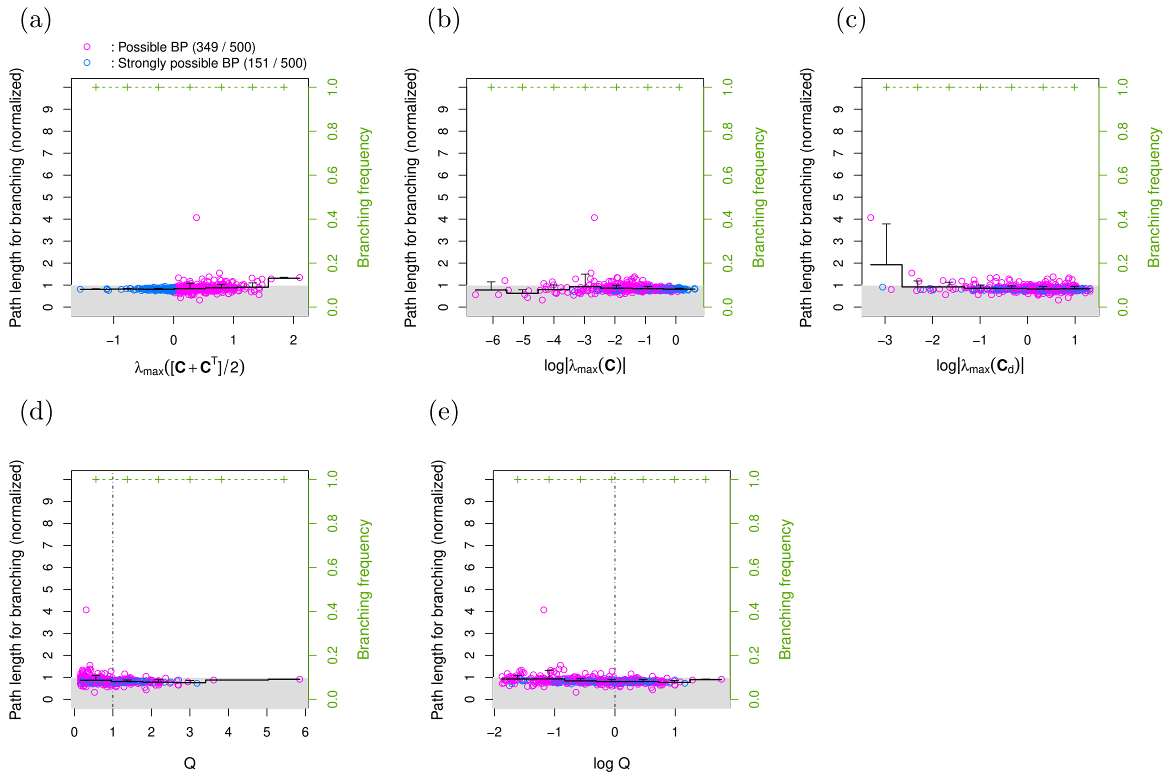

**Figure S5.** Normalized path lengths for the occurrence of evolutionary branching in simulated evolution as trait substitution sequences (three-dimensional trait spaces, i.e., $L=3$). Plotting formats are the same as those shown in figure S4. The simulation was conducted in the same manner as the simulation shown in figure. S4.

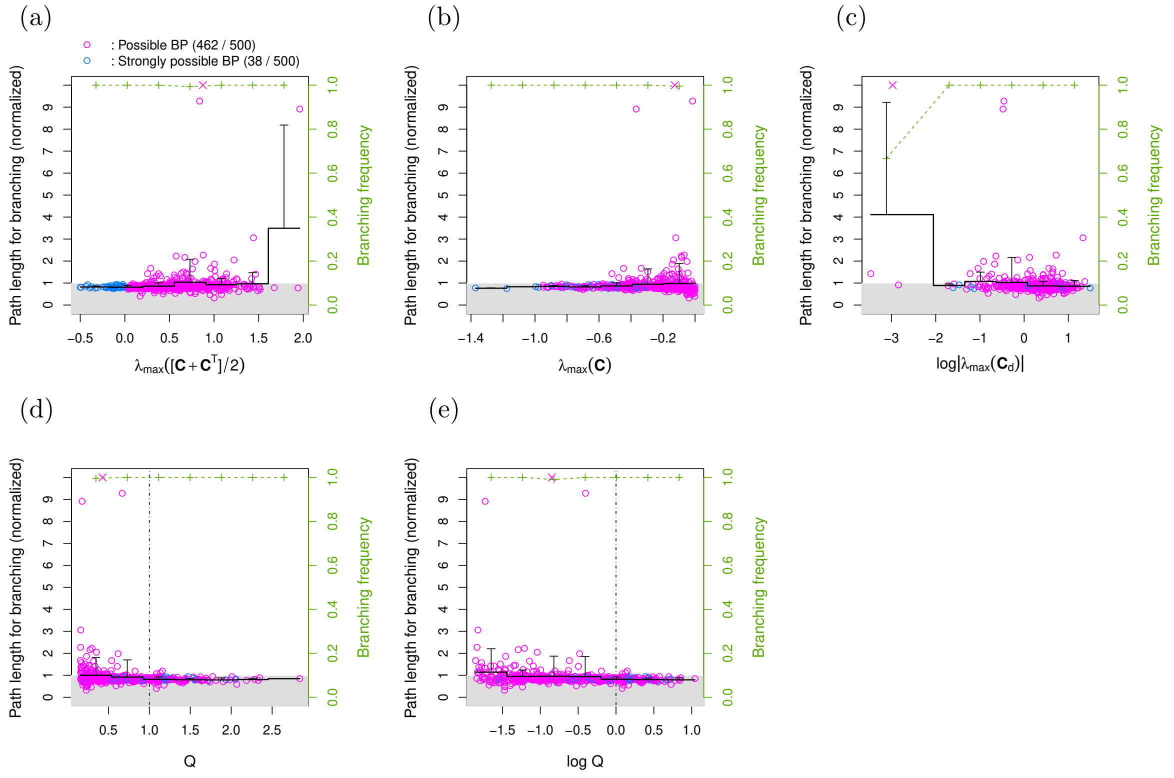

**Figure S6.** Normalized path lengths for occurrence of evolutionary branching in simulated evolution as trait substitution sequences (four-dimensional trait spaces, i.e., $L=4$). Plotting formats are the same as those shown in Fig. S2. The simulation was conducted in the same manner as the simulation shown in Fig. S2.

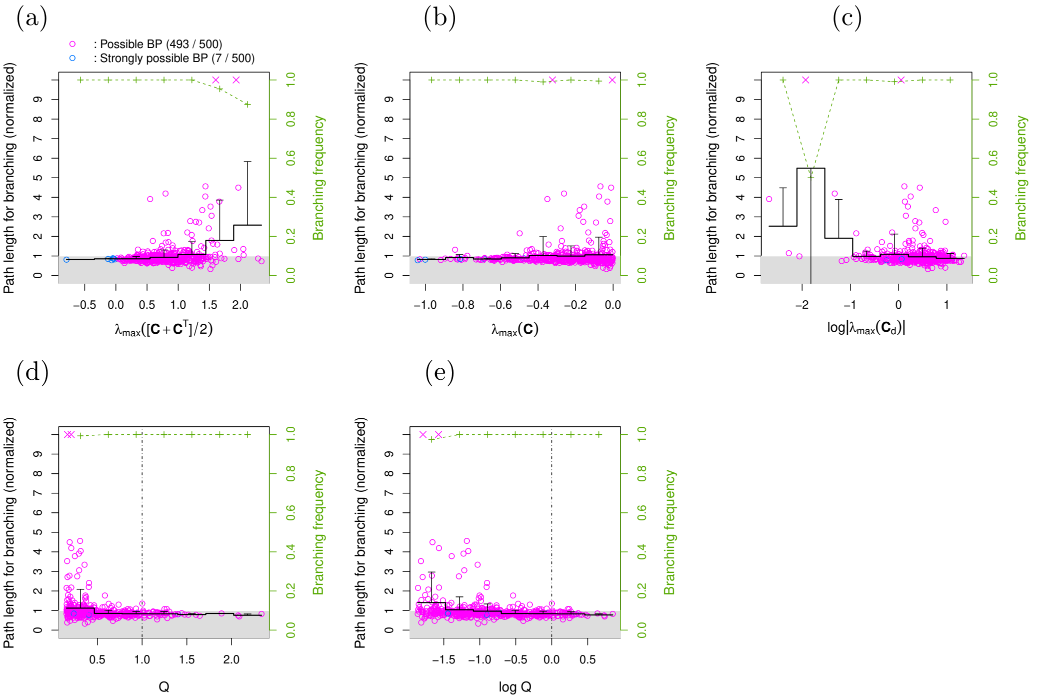

**Figure S7.** Normalized path lengths for the occurrence of evolutionary branching in simulated e ^ volution (five-dimensional trait spaces, i.e., $L=5$). Plotting formats are the same as those shown in figure S4. The simulation was conducted in the same manner as the simulation shown in figure S4.

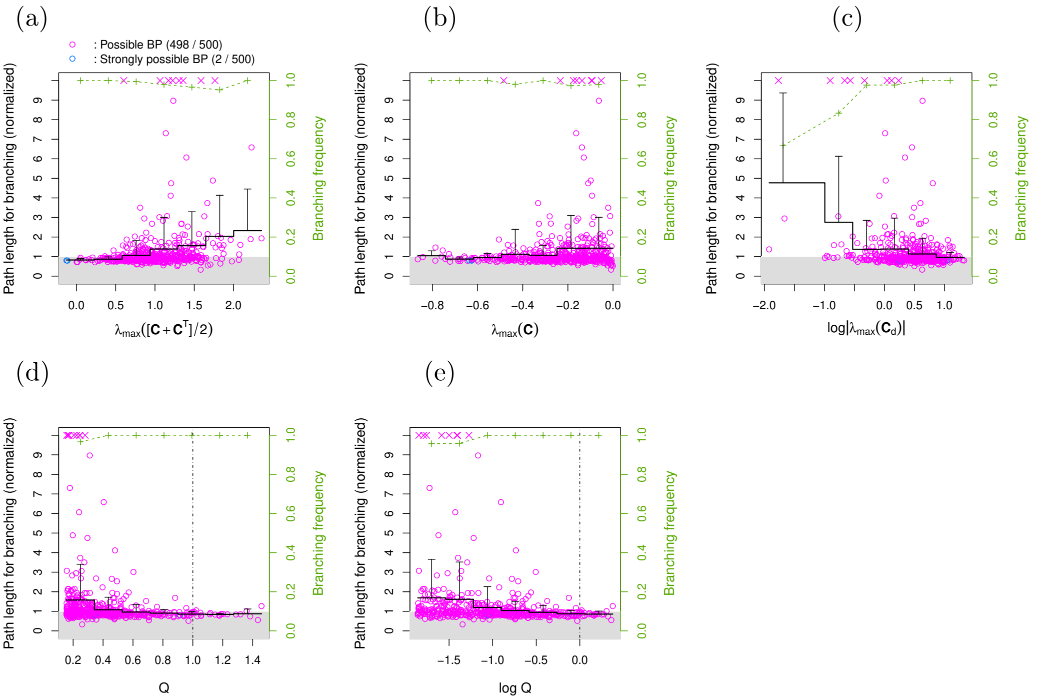

**Figure S8.** Normalized path lengths for the occurrence of evolutionary branching in simulated evolution as trait substitution sequences (six-dimensional trait spaces, i.e., $L=6$). Plotting formats are the same as those shown in figure S4. The simulation was conducted in the same manner as the simulation shown in figure S4.

### S7. Application example

#### S7.1. Calculation of trophic levels

We calculated trophic levels $T_1$, ..., $T_M$ for $M$ coexisting phenotypes analogously to the isotope-based method (Vander-Zanden et al., 1999), by solving the following simultaneous equations

$$T_i = 1 + \frac{\sum_{j=0}^{M} g_{i,j} T_j}{\sum_{j=0}^{M} g_{i,j}} \tag{S7.1}$$

for $i = 1$, ..., $M$ with $T_0 = 1$ (i.e., the trophic level of the main resource is assumed to be 1 for simplicity), where $g_{i,j}$ describes the assimilation amount of the $i$th phenotype through its predation on the $j$th phenotype, given by

$$g_{i,j} = c\rho\beta(x_i - y_j)n_j, \tag{S7.2}$$

and where $g_{i,0}$ describes the assimilation amount from the main resource, given by

$$g_{i,0} = r(x_i, y_i) - \sum_{j=1}^{M} \alpha(x_i, x_j)n_j. \tag{S7.3}$$

#### S7.2. Derivation of equations (22) and (23)

From the invasion fitness for a mutant $\mathbf{s}' = (x', y')$ under a monomorphic resident $\mathbf{s} = (x, y)$, given by equation (21) in the main text,

$$f(\mathbf{s}', \mathbf{s}) = 1 + \frac{1}{2} r_{\mathrm{xx}} x'^2 + \frac{1}{2} r_{\mathrm{yy}} y'^2 + [-\alpha(x' - x) + \rho c\beta(x' - y) - c\beta(x - y')]\,\hat{n}, \tag{S7.4}$$

the fitness gradients at $\mathbf{s}$ along $x$ and $y$ are given by

$$\begin{aligned}
g_x(x,y) &= \left[\frac{\partial f(\mathbf{s}',\mathbf{s})}{\partial x'}\right]_{\mathbf{s}'=\mathbf{s}} = r_{\rm xx}\,x + \left[-\frac{\partial \alpha(x'-x)}{\partial x'} + \rho c \frac{\partial \beta(x'-y)}{\partial x'}\right]_{\mathbf{s}'=\mathbf{s}} \hat{n} \\
&= r_{\rm xx}\,x + \rho c\,\hat{n} \left[\beta(x'-y)\,\frac{\partial\left(-\frac{[x'-y]^2}{2\,\sigma_\beta^2}\right)}{\partial x'}\right]_{\mathbf{s}'=\mathbf{s}} \\
&= r_{\rm xx}\,x - \rho c A\,[x-y]\,\beta(x-y), \\
g_y(x,y) &= \left[\frac{\partial f(\mathbf{s}',\mathbf{s})}{\partial y'}\right]_{\mathbf{s}'=\mathbf{s}} = r_{\rm yy}\,y - \left[c\frac{\partial \beta(x-y')}{\partial y'}\right]_{\mathbf{s}'=\mathbf{s}} \hat{n} \\
&= r_{\rm yy}\,y - c\,\hat{n} \left[\beta(x-y')\,\frac{\partial\left(-\frac{[x-y']^2}{2\,\sigma_\beta^2}\right)}{\partial y'}\right]_{\mathbf{s}'=\mathbf{s}} \\
&= r_{\rm yy}\,y - cA[x-y]\,\beta(x-y) \qquad \text{(S7.5)}
\end{aligned}$$

with $A = \hat{n}/\sigma_\beta^2$. From equation (S7.2) we get

$$\begin{aligned}
C_{\rm xx} &= \left[\frac{\partial g_x(x,y)}{\partial x}\right]_{\mathbf{s}=\mathbf{0}} = \left[\frac{\partial [r_{\rm xx}\,x - \rho c A[x-y]\,\beta(x-y)]}{\partial x}\right]_{\mathbf{s}=0} = r_{\rm xx} - c\rho A \\
C_{\rm xy} &= \left[\frac{\partial g_y(x,y)}{\partial x}\right]_{\mathbf{s}=\mathbf{0}} = \left[\frac{\partial r_{\rm yy}\,y - cA[x-y]\,\beta(x-y)}{\partial x}\right]_{\mathbf{s}=0} = -cA \\
C_{\rm yx} &= \left[\frac{\partial g_x(x,y)}{\partial y}\right]_{\mathbf{s}=\mathbf{0}} = \left[\frac{\partial [r_{\rm xx}\,x - \rho c A[x-y]\,\beta(x-y)]}{\partial y}\right]_{\mathbf{s}=0} = c\rho A \\
C_{\rm yy} &= \left[\frac{\partial g_y(x,y)}{\partial y}\right]_{\mathbf{s}=\mathbf{0}} = \left[\frac{\partial r_{\rm yy}\,y - cA[x-y]\,\beta(x-y)}{\partial y}\right]_{\mathbf{s}=0} = r_{\rm yy} + cA. \qquad \text{(S7.6)}
\end{aligned}$$

From equation (S7.4) we get

$$\begin{aligned}
D_{\rm xx} &= \left[\frac{\partial^2 f(\mathbf{s}',\mathbf{s})}{\partial x'^2}\right]_{\mathbf{s}'=\mathbf{s}=\mathbf{0}} = \frac{\partial}{\partial x'}\left[r_{\rm xx}\,x' + \left[-\frac{\partial \alpha(x'-x)}{\partial x'} + \rho c\frac{\partial \beta(x'-y)}{\partial x'}\right]\hat{n}\right]_{\mathbf{s}'=\mathbf{s}=0} \\
&= r_{\rm xx} + \left[-\frac{\partial^2 \alpha(x'-x)}{\partial x'^2} + \rho c\frac{\partial^2 \beta(x'-y)}{\partial x'^2}\right]\hat{n} \\
&= r_{\rm xx} + \left[\frac{1}{\sigma_\alpha^2} - \rho c\frac{1}{\sigma_\beta^2}\right]\hat{n} \\
&= r_{\rm xx} + \left[\frac{\sigma_\beta^2}{\sigma_\alpha^2} - c\rho\right]A, \\
D_{\rm xy} &= \left[\frac{\partial^2 f(\mathbf{s}',\mathbf{s})}{\partial x'\partial y'}\right]_{\mathbf{s}'=\mathbf{s}=\mathbf{0}} = \frac{\partial}{\partial y'}\left[r_{\rm xx}\,x' + \left[-\frac{\partial \alpha(x'-x)}{\partial x'} + \rho c\frac{\partial \beta(x'-y)}{\partial x'}\right]\hat{n}\right]_{\mathbf{s}'=\mathbf{s}=0} = 0 \\
D_{\rm yy} &= \left[\frac{\partial^2 f(\mathbf{s}',\mathbf{s})}{\partial y'^2}\right]_{\mathbf{s}'=\mathbf{s}=\mathbf{0}} = \frac{\partial}{\partial y'}\left[r_{\rm yy}\,y' - \left[c\frac{\partial \beta(x-y')}{\partial y'}\right]\hat{n}\right]_{\mathbf{s}'=\mathbf{s}=0} = r_{\rm yy} + cA. \qquad \text{(S7.7)}
\end{aligned}$$

#### S7.3. Derivation of equations (24-28)

##### Possible branching point conditions

According to table 1 in the main text, the conditions for $\mathbf{s}^* = \mathbf{0}$ to be a possible branching point under $\mathbf{V}_\mu = \sigma^2\,\mathbf{I}$ are simplified into

$$\begin{aligned}
\lambda_{\max}(\mathbf{C}) &< 0, \qquad \text{(S7.8)} \\
\lambda_{\max}(\mathbf{D}) &> 0, \qquad \text{(S7.9)} \\
\mathbf{e}^T[\mathbf{D}-\mathbf{C}]\,\mathbf{e} &> 0 \qquad \text{(S7.10)}
\end{aligned}$$

with $\mathbf{e}=\mathbf{v}_{\lambda\max}(\mathbf{D})$. We see from equations (S7.6) and (S7.7) that equation (S7.8) can be expressed as

$$\det(\mathbf{C}) = C_{\rm xx}C_{\rm yy}-C_{\rm xy}C_{\rm yx}=[r_{\rm xx}-c\rho A][r_{\rm yy}+cA]+\rho c^2A^2>0, \tag{S7.11}$$
$$\mathrm{tr}(\mathbf{C}) = C_{\rm xx}+C_{\rm yy}=[r_{\rm xx}-c\rho A]+[r_{\rm yy}+cA]<0. \tag{S7.12}$$

We see from equations (S7.6) and (S7.7) that $\lambda_{\max}(\mathbf{D})$ and $\mathbf{e}$ are given by

$$\lambda_{\max}(\mathbf{D}) = \max\{D_{\rm xx},D_{\rm yy}\} \tag{S7.13}$$
$$\mathbf{e} = \begin{cases}(1,0)^T & \text{for } \lambda_{\max}(\mathbf{D})=D_{\rm xx}\\ (0,1)^T & \text{for } \lambda_{\max}(\mathbf{D})=D_{\rm yy}\end{cases}. \tag{S7.14}$$

Note that $\lambda_{\max}(\mathbf{D})=D_{\rm yy}$ gives $\mathbf{e}=(0,1)^T$ and hence gives $\mathbf{e}^T[\mathbf{D}-\mathbf{C}]\,\mathbf{e}=D_{\rm yy}-C_{\rm yy}=0$. Therefore, equation (S7.10) requires $\lambda_{\max}(\mathbf{D})=D_{\rm xx}$, i.e.,

$$D_{\rm xx}-D_{\rm yy} = r_{\rm xx}-r_{\rm yy}+\left[\frac{\sigma_\beta^2}{\sigma_\alpha^2}-c(\rho+1)\right]A>0. \tag{S7.15}$$

In this case, equation (S7.6) becomes

$$D_{\rm xx} = r_{\rm xx}+\left[\frac{\sigma_\beta^2}{\sigma_\alpha^2}-c\rho\right]A>0, \tag{S7.16}$$

which gives $\mathbf{e}^T[\mathbf{D}-\mathbf{C}]\,\mathbf{e}=D_{\rm xx}-C_{\rm xx}=\frac{\sigma_\beta^2}{\sigma_\alpha^2}A>0$. Therefore, the conditions for $\mathbf{s}^*=\mathbf{0}$ to be a possible branching point are obtained as equations (7.11), (S7.12), (S7.15), and (S7.16), which are identical to equations (24) and (25) in the main text.

**Strongly possible branching point conditions**

According to table 1 in the main text, the conditions for $\mathbf{s}^*=\mathbf{0}$ being a strongly possible branching point are given by

$$\lambda_{\max}([\mathbf{C}+\mathbf{C}^T]/2) < 0, \tag{S7.17}$$
$$\lambda_{\max}(\mathbf{D}) > 0, \tag{S7.18}$$

with $\mathbf{e}=\mathbf{v}_{\lambda\max}(\mathbf{D})$. We see from equations (S7.6) and (S7.7) that equation (S7.17) can be expressed as

$$\det([\mathbf{C}+\mathbf{C}^T]/2) = C_{\rm xx}C_{\rm yy}-\frac{[C_{\rm xy}+C_{\rm yx}]^2}{4}=[r_{\rm xx}-c\rho A][r_{\rm yy}+cA]-\frac{1}{4}[1-\rho]^2c^2A^2>0, \tag{S7.19}$$
$$\mathrm{tr}([\mathbf{C}+\mathbf{C}^T]/2) = C_{\rm xx}+C_{\rm yy}=[r_{\rm xx}-c\rho A]+[r_{\rm yy}+cA]<0. \tag{S7.20}$$

In the same manner with equations (S7.13-S7.16), the conditions for equation (S7.18) are given by equations (S7.15), and (S7.16). Therefore, the conditions for $\mathbf{s}^*=\mathbf{0}$ to be a strongly possible branching point are obtained as equations (S7.19), (S7.20), (S7.15), and (S7.16), which are identical to equations (26) and (25) in the main text.

**Inevitable branching point conditions**

According to table 1 in the main text, the conditions for $\mathbf{s}^*=\mathbf{0}$ to be an inevitable branching point are given by

$$\mathbf{C}-\mathbf{C}^T = 0, \tag{S7.21}$$
$$\lambda_{\max}(\mathbf{C}) < 0, \tag{S7.22}$$
$$\lambda_{\max}(\mathbf{D}) > 0. \tag{S7.23}$$

We see from equations (S7.6) and (S7.7) that equation (S7.21) can be expressed as

$$C_{\mathrm{yx}} - C_{\mathrm{xy}} \;=\; c[1+\rho]\,A = 0, \tag{S7.24}$$

resulting in

$$c \;=\; 0. \tag{S7.25}$$

In this case, equation (S7.22) is expressed as

$$\det(\mathbf{C}) \;=\; C_{\mathrm{xx}}\,C_{\mathrm{yy}} - C_{\mathrm{xy}}\,C_{\mathrm{yx}} = r_{\mathrm{xx}}\,r_{\mathrm{yy}} > 0, \tag{S7.26}$$

$$\mathrm{tr}(\mathbf{C}) \;=\; C_{\mathrm{xx}} + C_{\mathrm{yy}} = r_{\mathrm{xx}} + r_{\mathrm{yy}} < 0, \tag{S7.27}$$

resulting in

$$r_{\mathrm{xx}} < 0 \quad , \quad r_{\mathrm{yy}} < 0. \tag{S7.28}$$

Because equations (S7.22) and (S7.25) give $D_{\mathrm{yy}} = r_{\mathrm{yy}} < 0$, we can transform equation (S7.23) into

$$D_{\mathrm{xx}} \;=\; r_{\mathrm{xx}} + \frac{\sigma_\beta^2}{\sigma_\alpha^2}\,A > 0. \tag{S7.29}$$

Therefore, the conditions for $\mathbf{s}^* = \mathbf{0}$ to be an inevitable branching point are obtained as equations (S7.25), (S7.28), and (S7.29), which are identical to equations (27) and (28) in the main text.
